# Complex genomic landscape of inversion polymorphism in Europe’s most destructive forest pest

**DOI:** 10.1101/2023.10.10.561670

**Authors:** Anastasiia Mykhailenko, Piotr Zieliński, Aleksandra Bednarz, Fredrik Schlyter, Martin N. Andersson, Bernardo Antunes, Zbigniew Borowski, Paal Krokene, Markus Melin, Julia Morales-García, Jörg Müller, Zuzanna Nowak, Martin Schebeck, Christian Stauffer, Heli Viiri, Julia Zaborowska, Wiesław Babik, Krystyna Nadachowska-Brzyska

## Abstract

In many species, polymorphic inversions underlie complex phenotypic polymorphisms and facilitate local adaptation in the face of gene flow. Multiple polymorphic inversions can co-occur in a genome, but the prevalence, evolutionary significance, and limits to complexity of genomic inversion landscapes remain poorly understood. Here, we examine genome-wide variation in one of Europe’s most destructive forest pests, the spruce bark beetle *Ips typographus*, scan for polymorphic inversions, and test whether inversions are involved in key adaptations in this species. We analyzed 240 individuals from 18 populations across the species’ European range and, using a whole-genome resequencing approach, identified 27 polymorphic inversions covering approximately 28% of the genome. The inversions vary in size and in levels of intra-inversion recombination, are highly polymorphic across the species range, and often overlap, forming a complex genomic architecture. We test several mechanisms, including directional selection, overdominance and associative overdominance that can contribute to the maintenance of inversion polymorphisms in the genome. We show that the heterogeneous inversion landscape is likely maintained by the combined action of several evolutionary forces and that inversions are enriched in odorant receptor genes encoding key elements of recognition pathways for host plants, mates, and symbiotic fungi. Our results indicate that the genome of this major forest pest of growing social, political, and economic importance harbors one of the most complex inversion landscapes described to date posing a question about limits of genomic architecture complexity.

## Introduction

Large structural variants (SVs), such as chromosomal inversions, translocations, insertions, and duplications, were among the earliest mutations described in natural populations (McClung 1905; Sturtevant 1921), but their systematic identification and in-depth analysis has only become possible with recent advances in sequencing technology (Wellenreuther et al. 2019). The increasing accumulation of genomic data has revealed not only the presence of structural variation between and within many species but also its important role in species adaptation and speciation (Lamichhaney et al. 2015; Hof et al. 2016; Cheng et al. 2018; Wellenreuther and Bernatchez 2018; Faria, Chaube, et al. 2019; Todesco et al. 2020; Mérot et al. 2021; Harringmeyer and Hoekstra 2022; Saitou et al. 2022). In particular, polymorphic chromosomal inversions have been at the center of recent debate about their potential to redirect the evolutionary process (Berdan et al. 2023).

Polymorphic inversions are chromosomal segments that occur in two orientations within populations: collinear and inverted haplotypes/arrangements. Inversions have been shown to be involved in speciation, local adaptation, and maintenance of complex phenotypes (Lamichhaney et al. 2015; Lohse et al. 2015; Wellenreuther and Bernatchez 2018; Fuller et al. 2019). This is due to a key property of inversions: they suppress recombination within heterozygotes and thereby prevent separation of coadapted variants. Because of their role as recombination modifiers, inversions can act as supergenes, i.e., large elements of genomic architecture with multiple linked functional elements (Thompson and Jiggins 2014). Supergenes keep coadapted alleles together in the face of gene flow and prevent the formation of maladaptive recombinant genotypes.

Classic examples of supergenes include inversions associated with different mating strategies in ruffs (*Calidris pugnax*; Lamichhaney et al. 2015), mimicry phenotypes in *Heliconius* butterflies (Joron et al. 2011), and social organization in fire ants (*Solenopsis* spp.; Purcell et al. 2014; Gutiérrez-Valencia et al. 2021). In many other species, polymorphic inversions define locally adapted ecotypes (Lowry and Willis 2010; Kirubakaran et al. 2016; Koch et al. 2021; Matschiner et al. 2022; Reeve et al. 2023) or exhibit spatial frequency differences, e.g. by forming geographic and climatic gradients (Ayala et al. 2014; Kapun et al. 2016; Ayala et al. 2017; Mérot et al. 2021). While the vast majority of described cases are organisms with one or a few inversions, several recent studies have reported species with many polymorphic inversions (Faria, Chaube, et al. 2019; Todesco et al. 2020; Harringmeyer and Hoekstra 2022; Porubsky et al. 2022; Reeve et al. 2023). These recent findings raise questions about the prevalence and evolutionary significance of polymorphic inversions in natural populations. Are inversion-rich genomes the exception or the rule? How much of the genome can be situated within polymorphic inversions and, consequently, how large can the fraction of the genome with reduced recombination be? The latter question is particularly important because, in addition to suppressing recombination and keeping coadapted alleles together, inversion heterozygotes will also prevent the formation of new allelic combinations and thus reduce the efficacy of natural selection (Roesti et al. 2022).

Given the importance of recombination in purging deleterious mutations (Keightley and Otto 2006), a substantial fraction of the genome that is situated within polymorphic inversions raises other questions. First, as the degree of recombination suppression depends on the number of heterozygous genotypes, what is the probability of an individual being heterozygous at multiple inversion regions? Second, are polymorphic inversions where both arrangements are equally common across the species’ range more common than inversions with one common and one rare haplotype? Third, how common are mechanisms that potentially can mitigate negative consequences of reduced recombination? Such mechanisms include double crossover-events and gene conversion in heterozygotes, which can reshuffle allelic content between large parts of two inversion arrangements and thereby reduce mutational load and create new haplotypes (Berdan et al. 2021; Charlesworth 2023).

Long-term persistence of two inversion haplotypes can be facilitated by two main types of selection: divergent and balancing (Faria, Johannesson, et al. 2019). Divergent selection can favor different inversion genotypes in different environments and, when coupled with reduced migration between divergent populations, can lead to speciation. Even when intraspecific gene flow is high, divergent selection can lead to divergent ecotypes associated with locally advantageous inversion genotypes. Alternatively, balancing selection may maintain balanced inversion polymorphisms over time via several, not mutually exclusive, mechanisms, such as overdominance, negative frequency dependence, antagonistic pleiotropy, and spatially or temporally varying selection (Connallon and Clark 2014; Faria, Johannesson, et al. 2019).

Importantly, regardless of the selection mechanism, inversions will accumulate mutations independently since recombination is suppressed in inversion heterozygotes. This will lead to differentiated allelic content and increased differentiation between inversion haplotypes over time (Faria, Johannesson, et al. 2019). Each inversion haplotype can thus be treated as a separate “population” with a size that corresponds to the frequency of that arrangement within studied population or species. Rare inversion arrangements will experience a high mutational load due to reduced recombination and limited purging because of rarity of heterozygotes. However, deleterious mutations can also accumulate on more frequent inversion haplotypes (Berdan et al. 2021) and lead to associative overdominance. This contributes to the maintenance of inversion polymorphisms since independent accumulation of mutations continues over time but recessive deleterious alleles private to an inversion arrangement are not visible to selection in inversion heterozygotes.

The Eurasian spruce bark beetle (*Ips typographus* (L.): Curculionidae: Scolytinae; hereafter the spruce bark beetle), plays a key role in Eurasian forest ecosystems. Under endemic conditions, it attacks weakened Norway spruce (*Picea abies*) trees. However, if spruce resistance is compromised by certain abiotic disturbances (e.g. snowbreaks, windfalls, high temperatures, drought), an increased availability of stressed trees can trigger mass-propagation, leading to rapid population increase and devastating outbreaks. The extent of recent outbreaks is unprecedented, and impacts will likely increase in the coming decades in response to climate change (Biedermann et al. 2019; Bentz et al. 2021; Müller et al. 2022). For example, during the first decade of the 21st century the spruce bark beetle killed an estimated 14.5 million m^3^ of timber per year on average. The Czech Republic provides a particularly striking example of the beetles’ destructive potential. During the peak outbreak years 2017-2019 the beetles killed annually 3.1-5.4% of the country’s growing stock of Norway spruce, which in 2019 translated to 23 million m^3^. Historically, spruce forests in Central Europe have been most heavily affected by bark beetle outbreaks, while in northern Europe outbreaks have been less frequent and destructive (Hlásny et al. 2019). However, this may change with climate warming that probably will make the boreal forests of northern Europe more vulnerable to bark beetle outbreaks (Lange et al. 2006; Jönsson et al. 2007; Müller 2011; Bentz et al. 2021; Müller et al. 2022). As an example, heatwaves and severe summer drought in Sweden in 2018 initiated a bark beetle outbreak killing over 30 million m^3^ Norway spruce in the next years (Öhrn et al. 2021; Wulff and Roberge 2022).

Increasing bark beetle attacks and other forest disturbances have already triggered social and political conflicts in parts of Europe and have highlighted the urgent need for improved management strategies (Müller 2011; Vega and Hofstetter 2015; Tomáš Hlásny et al. 2021). Indeed, a rapidly growing body of research focuses on the species’ ecology, and the causes and consequences of outbreaks (reviewed in: Wermelinger 2004; Vega and Hofstetter 2015; Biedermann et al. 2019; Wermelinger and Jakoby 2022). Despite this enormous interest, one aspect of the species’ biology remains largely unexplored: we know almost nothing about the species’ genome-wide variation and the evolutionary mechanisms that shape this variation. The lack of population genomics studies restricts our understanding of the genomic basis of adaptation and adaptive potential in the spruce bark beetle. Such information could provide a critical missing link between applied and basic research and serve as a foundation for effective management. This is particularly important because, as revealed in this study, the spruce bark beetle genome harbors a complex inversion polymorphism landscape that may play a critical role in many evolutionary processes, including key species adaptations (Kirkpatrick and Barton 2006; Wellenreuther and Bernatchez 2018; Berdan et al. 2021).

Here, we investigated genome-wide variation across spruce bark beetle populations with a special focus on chromosomal inversion polymorphisms. We found one of the most complex polymorphic inversion architectures described to date and investigated several evolutionary mechanisms, including directional selection, overdominance and associative overdominance, that can maintain inversion polymorphisms in the genome. We also tested associations between inversion polymorphisms and two key fitness-related traits - diapause and olfaction. Our results suggest that inversions are involved in key species adaptations and raise questions about the prevalence and role of inversion polymorphisms in species characterized by large effective population sizes, little geographic subdivision, and no clear phenotypic variation across the species range.

## Results and Discussion

We resequenced whole genomes of the spruce bark beetles from 18 European populations (Figure 1, Table S1). After initial raw data processing from 253 individuals (including data quality check, trimming and GATK guided variant calling, genotyping, recalibration and filtering; for details see Supplementary Information), we used 240 individuals in downstream analyses. The mean per individual sequencing depth was 23.2x (range: 5 – 53x). Sequencing coverage on IpsContig9 was consistently lower in individuals sexed as males (on average 0.57 individual coverage). Thus, we considered IpsContig 9 to be a sex (X) chromosome (females carry XX and males XYp chromosomes). After quality filtering, we retained 5.245 million SNPs covering the entire length of the genome assembly (236.8 Mb) but analyzed a subset of 5.067 million SNPs located on the 36 longest contigs and that represented 78% of the assembly.

**Figure 1.**
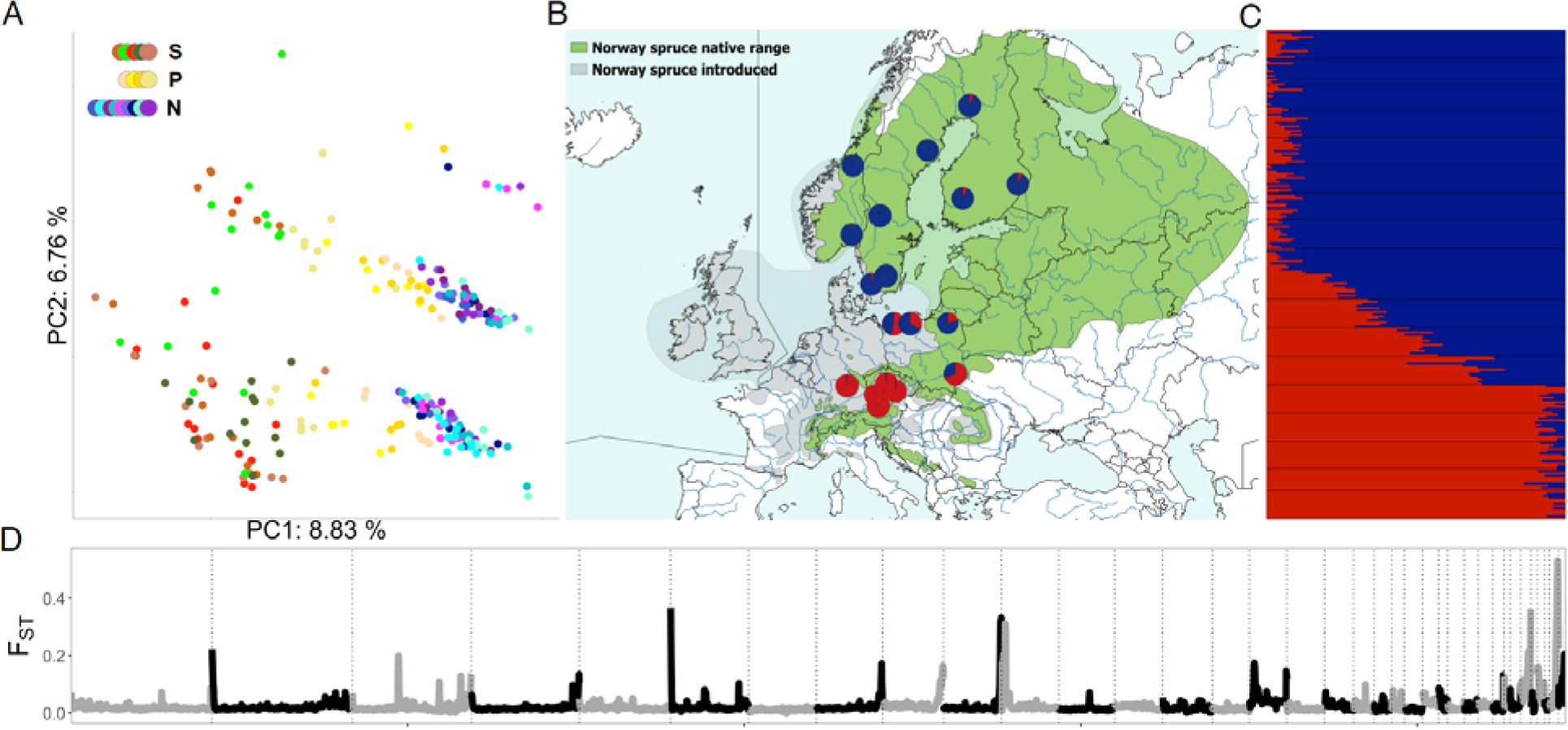
Genomic structure and differentiation in *Ips typographus*. (A) Whole-genome PCA, where colors correspond to what population an individual belongs to. Legend indicates geographical grouping of populations: S-South, P-Poland, N-North. (B, C) Geographical distribution and genetic differentiation of the 18 beetle populations analyzed. Blue and red are the colors of the main genetic groups: southern (red), northern (blue). (D) Genome-wide genetic differentiation (*F_ST_*) calculated between northern and southern groups. Vertical lines separate different contigs shown in grey and black.

Alignment of the *Ips typographus* and *Ips nitidus* genome assemblies indicated no major misassembly problems (*Ips typographus* contigs align to single *Ips nitidus* chromosomes) and high synteny between karyotypes of both species (Figure S1). Given these results, contig sizes, genome assembly quality scores and species karyotype, it is likely that many spruce bark beetle contigs represent entire chromosome arms. Such genome quality is sufficient for all downstream analyses, in particular SNP-based polymorphic inversion identification (Mérot et al. 2021; Reeve et al. 2023).

### Complex genomic inversion landscape

Twenty-nine candidate inversions (“inversions” henceforth) were identified following the criteria described in the Methods section (Table 1, Figure 2, Figure S2, and Figure S3). Briefly, we considered a genomic region to harbor a polymorphic inversion if local PCA analysis identified the region as an outlier, and/or the region exhibited high linkage disequilibrium (LD) and PCA performed on SNPs from this region separated individuals not by geography but into three distinct groups (with a group classified as putative heterozygous individuals characterized by higher heterozygosity compared to the other groups). Two putative inversions (Inv16.1 and Inv23.1) were most likely part of the same inversion: they are in strong LD (Figure S4) and the same beetles were genotyped as homo- and heterozygotes in both, which is unlikely to occur for two unlinked inversions. The same situation was found for Inv16.2 and Inv23.2. No other inversions were in strong LD with each other, which would suggest co-segregation (Figure S4). Thus, overall, we found 27 inversions in 17 contigs, including one located on the X chromosome (Inv9). Approximate inversion sizes varied from 0.1 to 10.8 Mb (Table 1) and inversions constituted approximately 28% of the analyzed part of the genome and at least 18.6% of the entire assembly. Estimated inversion ages ranged from 0.5 to 2.6 My (Table 1, assuming a mutation rate of 2.9 × 10^-9^; see Table S1 for results for different mutation rates) and were strongly negatively correlated with the major inversion haplotype frequency (Figure S5). Inversion regions exhibited a reduced population recombination rate in heterozygous individuals (Figure S2) and moderate to high genetic differentiation between inversion arrangements (*F_ST_* between homozygous individuals was 0.15-0.64; Figure 1; Figure S2).

**Figure 2.**
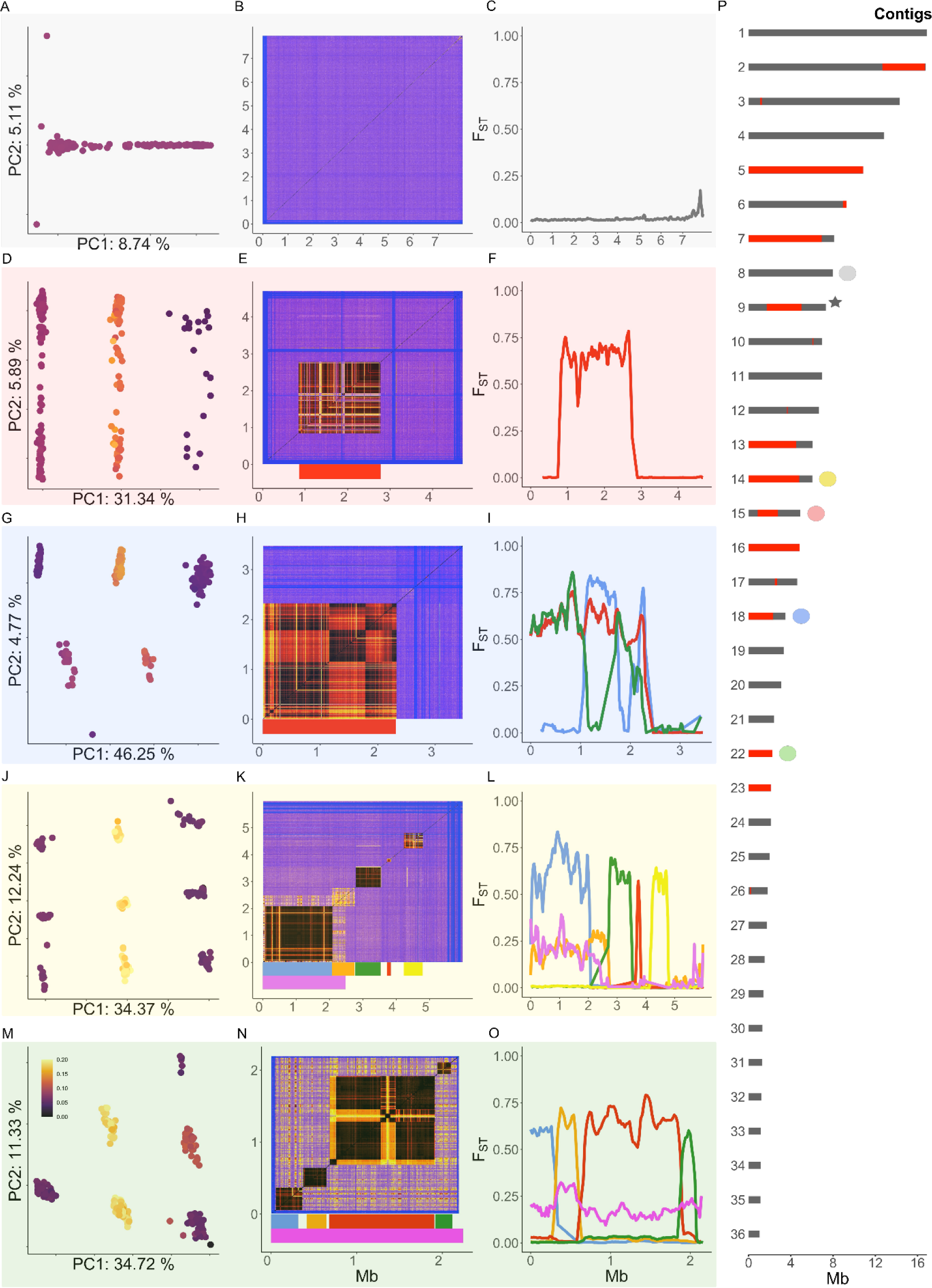
Identification of chromosomal inversions in *Ips typographus*. Each row of the figure panels shows the results of per contig PCA, per contig linkage disequilibrium analysis, and genetic differentiation (*F*_ST_) analysis for a selected contig. Panels A-C show results for a contig with no inversions (IpsContig8) and little differentiation between southern and northern populations. Panels D-O show different contigs with increasingly complex inversion patterns: a single inversion (D-F, Inv15); a single inversion with a double-crossover signal (G-I, Inv18); multiple adjacent inversions with one inversion overlapping with the first two inversions on the contig (J-L, Inv14.1, Inv14.2, Inv14.3, Inv14.4, Inv14.5, and Inv.14.6); multiple adjacent inversions with one large inversion overlapping with several smaller ones (M-O, Inv22.1, Inv22.2, Inv22.3, Inv22.4, and Inv22.5). The large overlapping inversion is visible as a yellow background in the LD plot (N). The PCA grouping shown in (M) corresponds to genotypes of the largest inversion on IpsContig22 (Inv22.3), including genotypes that include haplotypes produced with the double crossover event. The last panel (P) shows the 36 largest contigs of the spruce bark beetle genome with inversions indicated in red. Colored dots indicate the inversions that are presented in detail in panels A-O. Colored bars indicate the position of each inversion. Colors of the bars in plots E, K and N correspond to adjacent *F*_ST_ plots (panels F, L and O), where *F*_ST_ is calculated between haplotypes of indicated inversion along the contig in question. In the plot I *F*_ST_ is calculated between three haplotypes: homozygotes for two arrangements and the homozygotes for recombinant haplotype created after putative double crossover event. Dots in the PCA plots represent individual beetles and are colored according to the heterozygosity of the individual (darker color represents low heterozygosity). Both axes in LD plots (B, E, H, K, N) represent positions along the contig in megabases (Mb); low levels of linkage are shown in blue and higher levels in yellow to dark red. Sex chromosome is indicated with a grey star in P panel.

**Table 1.**
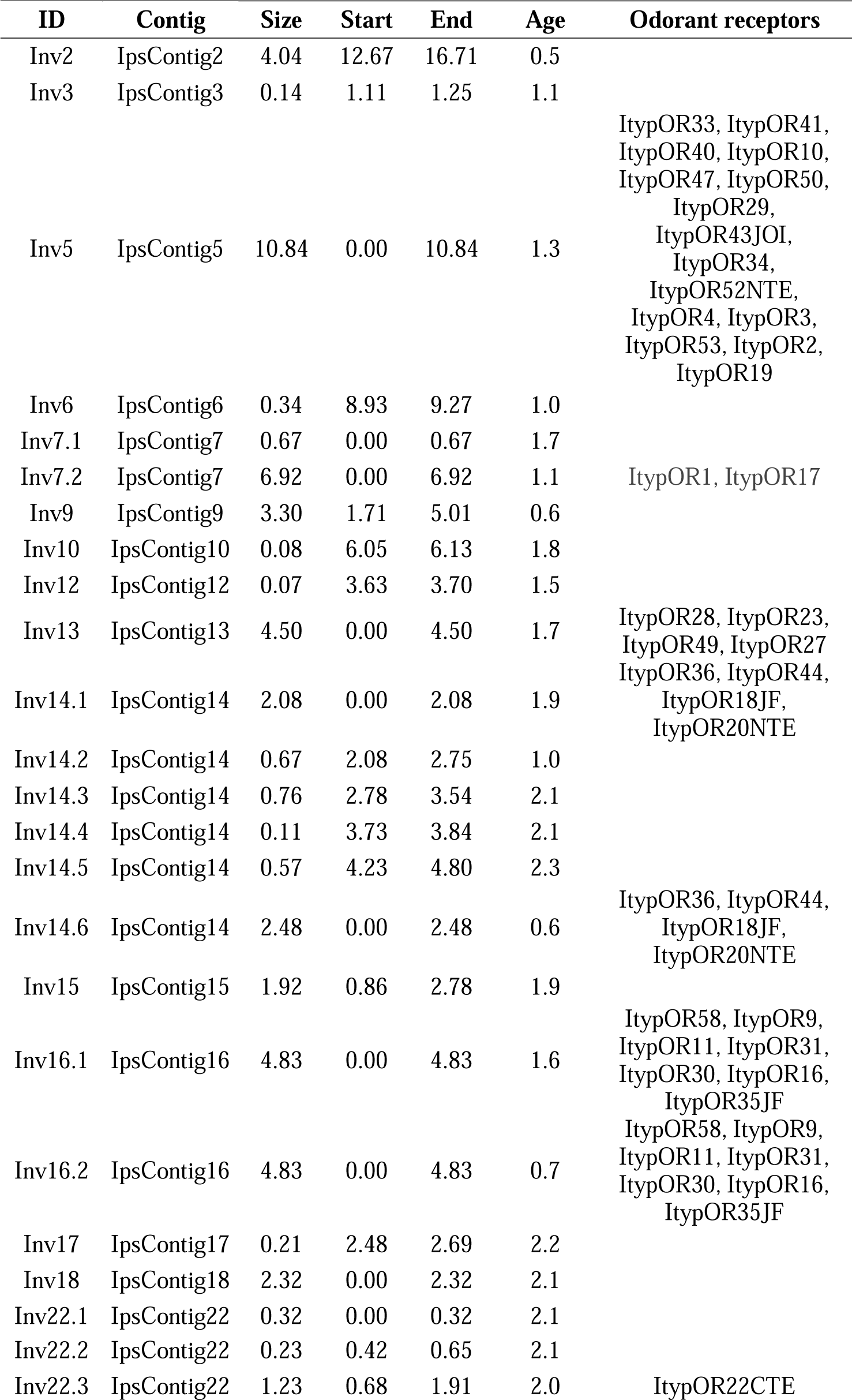
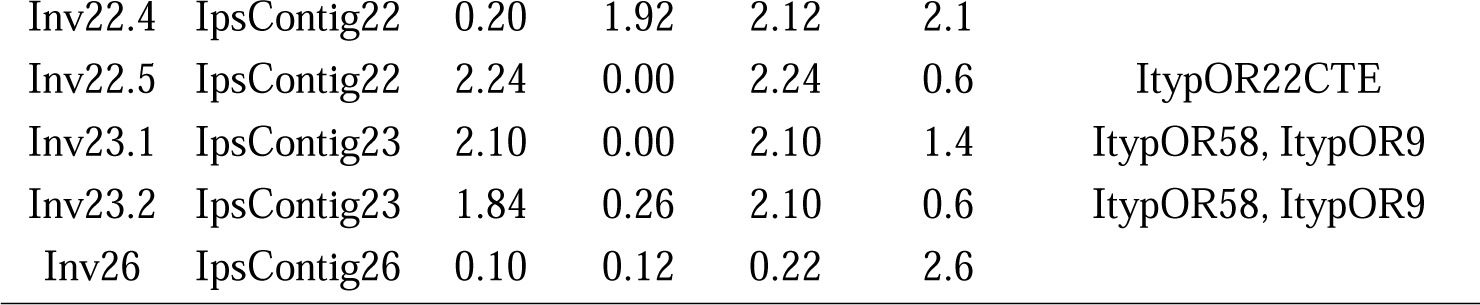
List of identified chromosomal inversions in the *Ips typographus* genome. ID: inversion name; Contig: contig name; Size: size of the inversions (Mb); Start and End: coordinates of the inversion (Mb); Age: approximate age of the inversion in Myr; Odorant receptors: odorant receptors present within inversion (in the order they appear along the contig sequence). Note that Inv16.1 and Inv23.1 are parts of the same inversion and Inv16.2 and Inv23.2 are a part of another single inversion.

While 12 contigs contained single inversions, five contigs showed patterns consistent with multiple adjacent or overlapping inversions (Figure 2, Figure S2, Figure S3). IpsContig14 and IpsContig22 contained complexes of multiple adjacent and sometimes overlapping inversions (Figure 1 J-O), and three other contigs contained two overlapping inversions each (IpsContigs: 7, 16 and 23, Figure S2 and Figure S3). Additionally, four putative double-crossover events were identified within four inversions (Inv5, Inv18, Inv22.3, Inv22.4; Figure 1, Figure S2 and Figure S3).

While the variation patterns observed on contigs with single inversions were clear and leave no doubt about the presence of polymorphic inversions, the complex patterns (overlapping inversions and inversions with double crossover events) were more difficult to interpret and deserve additional explanation and caution. Putative overlapping inversions were identified on the basis of overlapping clusters of high LD, PCA clustering into three distinct groups along PC2 or PC3 (in addition to the three distinct groups along PC1), and moderate to high *F_ST_* between individuals classified as alternatively homozygous (Figure 2 and Figure S3). In the case of putative double-crossover events, the observed LD clusters were separated by regions of lower LD and intermediate groups of individuals were visible between the main clusters along the PC1 (Figure S3). The *F_ST_* between individuals classified as alternatively homozygotes was significantly reduced in the region of the putative double-crossover (Figure 2 and Figure S2, S3). The most difficult to disentangle patterns were associated with Inv2, Inv5, Inv7.1 and Inv7.2.

Putative Inv2 showed a signature of overlapping LD clusters (overlapping inversions) but ambiguous PCA clustering (Figure S3). The patterns were clear and consistent with inversion polymorphisms when only populations from the southern region were analyzed (Figure S6; thus, Inv2 was genotyped only in this part of the species range), suggesting other structural differences between the southern and northern regions or unidentified genome assembly problems. Principal component analysis of Inv5 identified three clusters that most likely correspond to three inversion genotypes, but also several smaller clusters that may indicate multiple independent recombination events (double-crossover events that occurred at different locations within Inv5, Figure S2 and Figure S3) or some assembly problems within Inv5 (however, IpsContig5 aligns perfectly with *Ips nitidus* chromosome 9, Figure S1). Thus, Inv5 was only genotyped for three major genotype groups for which multiple lines of evidence (LD, heterozygosity, *F_ST_*) suggested an inversion polymorphism. Inv7.1 and Inv7.2 are characterized by overlapping LD clusters, and PCA and *F_ST_* patterns consistent with overlapping inversions. However, Inv7.2 showed low LD in the middle of the inversion and unexpected clustering of individuals in one of the clusters along PC1 – the cluster was split into two distinct groups with individuals clustered according to their geographic location within each of these groups (Figure S2 and Figure S3). This suggests possible misassembly and it is likely that Inv7.2 is shorter than predicted from the size of the LD boundaries. Genotyping of Inv16.2 and Inv23.2 was only possible in part of the species range, as PCA clustering into three genotype groups was only visible when analyzing the northern group only (Figure S6).

Even though we are not able to rule out that some of the patterns were created or disturbed by potential misassembly or additional structural rearrangements (e.g., duplications; Kim et al. 2022), the evidence for the vast majority of polymorphic inversions in the spruce bark beetle is strong. Extensive collinearity with *Ips nitidus* suggested no major misassembly problems and further supports our identification of polymorphic inversions in the spruce bark beetle genome (Figure S1; Wang et al. 2023). Long-read sequencing and Hi-C data is needed to shed light on the problematic/complex cases as well as to precisely identify the inversion boundaries. It was not surprising that we detected polymorphic inversions in the spruce bark beetle genome, as there are many well-known examples of polymorphic inversions in natural populations (Wellenreuther and Bernatchez 2018). What was striking, however, was the complex genomic landscape of polymorphic inversions we found in this species. The spruce bark beetle has at least 27 large inversions covering a substantial part of the genome (28% of the analyzed part of the genome). Numerous (a dozen or more) polymorphic inversions have been described for several species (e.g. Faria, Chaube, et al. 2019; Tigano et al. 2021; Harringmeyer and Hoekstra 2022) but it remains an open question whether many polymorphic inversions within species is the exception or the rule.

The exceptionally complex inversion architecture we found in the spruce bark beetle, with multiple adjacent and often overlapping inversions, resembles well known examples from *Heliconius* butterflies (Jay et al. 2021) or fire ants (Wang et al. 2013). In these insects, multiple adjacent inversions are the basis for mimicry phenotypes and complex social organization, respectively. The presence of clusters of adjacent inversions and inversion overlaps are consistent with theoretical expectations of stepwise extension of recombination suppression on supergenes (Jay et al. 2022) and with a highly polygenic architecture of adaptation (Schaal et al. 2022).

### Genome-wide variation and its geographic structuring – collinear vs. inversion regions

Analyses based on both the whole genome and collinear parts only revealed a clear latitudinal structuring of genetic variation in the spruce bark beetle (Figure 1). Both PCA and NGSadmix supported the presence of two distinct genetic groups corresponding to southern and northern populations, with Polish populations showing varying degrees of admixture between the two clusters. Based on these results, we divided the 18 studied populations into a northern, southern, and Polish group. Despite unambiguous NGSadmix division into two genetic clusters, the genome-wide genetic differentiation between the northern and southern group was very low (*F_ST_*= 0.021). Similarly, *F_ST_* between all population pairs showed low levels of differentiation, ranging from 0.000 to 0.035 (calculated excluding inversions; Table S2). Mean genome-wide nucleotide diversity was moderate (π = 0.0062) and per population π ranged from 0.0055 to 0.0066 (Figure S7). There was a weak negative correlation between nucleotide diversity and latitude (r^2^ = 0.32, p = 0.033, Figure S8), as northern populations had slightly lower genetic variation than southern populations (π_southern_= 0.0065; π _northern_ = 0.0061; π _Polish_= 0.0066).

Southern populations had an excess of rare alleles and, consequently, had more negative Tajima’s D values along the genome than northern populations (mean Tajima’s D was −0.458 and −0.062 in the southern and northern group, respectively; Figure S7). All these results are consistent with previous phylogeographic studies of the spruce bark beetle that analyzed a much smaller number of genetic markers. The data suggests high levels of connectivity among populations and a very recent differentiation into two genetic clusters (Stauffer et al. 1999; Sallé et al. 2007; Bertheau et al. 2013; Mayer et al. 2015). More recent RADseq data confirms a very weak genetic structure in the spruce bark beetle across much of Sweden (Ellerstrand et al. 2022), as is expected in a species with high dispersal (Nilssen 1984) and/or recent divergence. Tajima’s D values indicate different demographic history of southern and northern group (e.g., recent expansion of southern populations and bottleneck in northern populations, Figure S7).

Inversion regions in the spruce bark beetle did not structure in the same way as the collinear part of the genome and in most of the cases do not show any clustering into a southern and northern group (Figure S2). Frequencies of five inversions show significant correlation with latitude (Figure 3; r^2^ ranged from 0.34 to 0.68; see more discussion below). Almost all identified inversions were polymorphic across the European species range, except for one inversion (Inv9) that was polymorphic only in northern populations. For three inversions (Inv2, Inv5, Inv16.2+Inv23.2) unambiguous genotyping was only possible across part of the species range (see above). The differentiation between inversion haplotypes was moderate or high (mean *F_ST_* range 0.15-0.64), (Figure 2; Figure S2) and, according to our age estimates, the origin of the inversions in the spruce bark beetle predates the Last Glacial Period. Many inversions may be several million years old, likely also predating the within-species differentiation into a southern and northern group. This was not unexpected, as many known inversion polymorphisms have been segregating within species for hundreds of thousands or millions of years (see table in (Wellenreuther and Bernatchez 2018)), sometimes even persisting through multiple speciation events (Brelsford et al. 2020).

**Figure 3.**
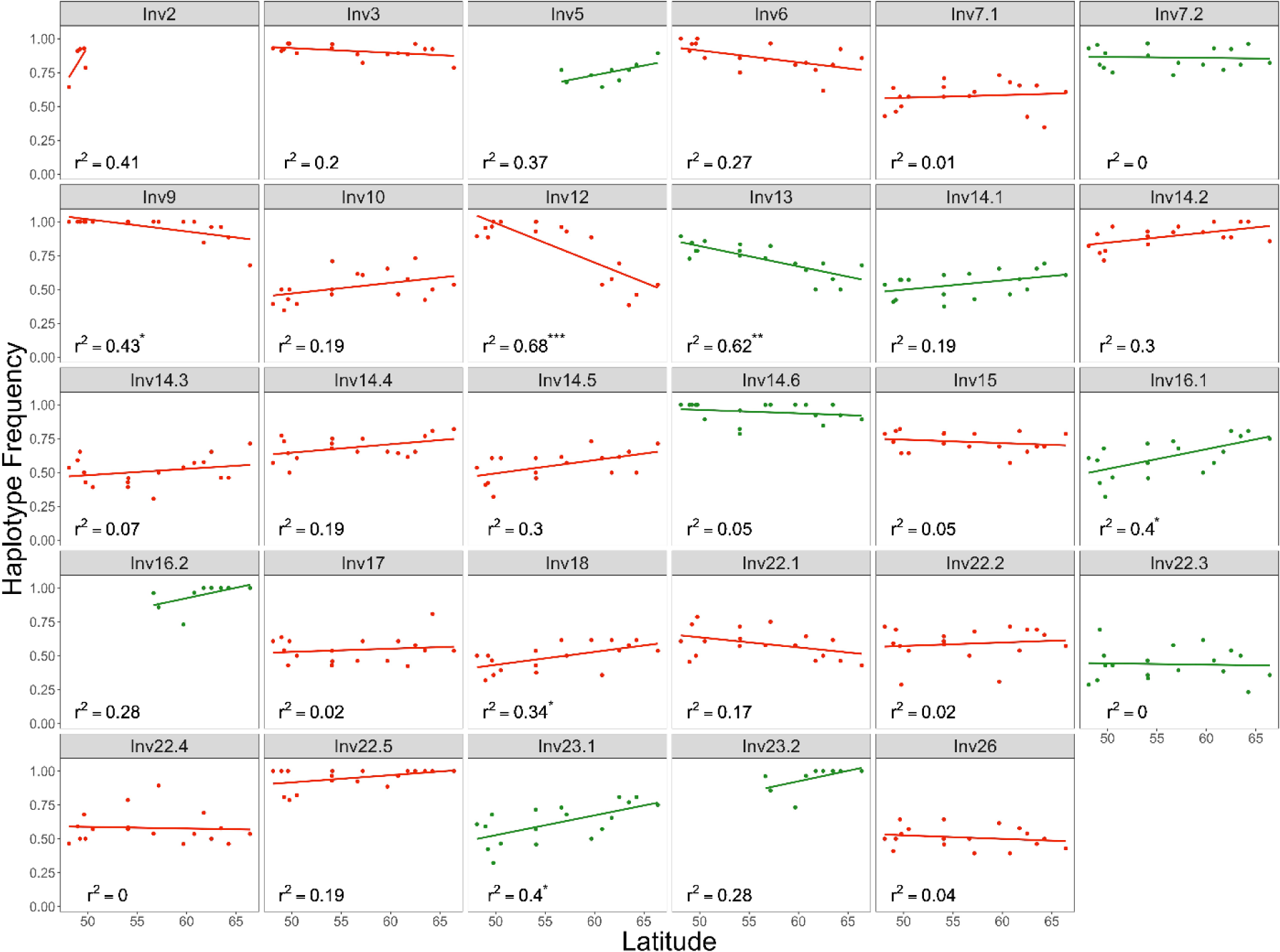
Correlation between inversion haplotype frequency and population latitude for chromosomal inversions detected in European *Ips typographus* populations. Significant correlations are indicated with asterisks (*: p < 0.05; **: p < 0.01; ***: p < 0.001). Inversions harboring odorant receptors are indicated in green.

### Inversions and key fitness-related traits

Inversion polymorphism is often associated with the maintenance of complex polymorphic phenotypes (Schwander et al. 2014; Lamichhaney et al. 2015). Although the spruce bark beetle does not exhibit easily identifiable phenotypes, such as distinct color patterns or mating types, we were able to test for associations between inversions and two complex traits of key importance for many insects: diapause and olfaction. Diapause allows species to suspend development during unfavorable conditions. While multiple environmental factors can influence this complex process, many aspects of diapause, such as its induction and termination, are heritable (Roff 1996) and may be controlled by a small number of loci or be highly polygenic (Paolucci et al. 2016; Pruisscher et al. 2018; Kozak et al. 2019; Denlinger 2022). The spruce bark beetle exhibits two diapause strategies that could be associated with polymorphic inversions: a facultative photoperiod-regulated diapause and an obligate photoperiod-independent diapause (Schebeck et al. 2022). However, we found no association between diapause phenotypes and inversion genotypes (exact G test, Table S3, Figure S9), nor did we find any highly differentiated genomic regions between facultatively and obligately diapausing individuals (Figure S10). This suggests a polygenic nature of diapause phenotypes in the spruce bark beetle.

Olfaction is another key fitness-related trait in many insects, including bark beetles. Insect odorant receptors (ORs) are encoded by a large and dynamically evolving gene family. Some of the receptors are evolutionarily conserved across species within insect orders, however, many are species- or genus-specific. In the spruce bark beetle, detection of odorants is essential for host and mate finding, as well as recognition and maintenance of symbiosis with specific fungi (Hansson and Stensmyr 2011; Kandasamy et al. 2019). We examined 73 antennally expressed ORs (Yuvaraj et al. 2021) and found that 46% of these were located within inverted regions (Table 1), even though inverted regions constituted only 28% of the analyzed part of the genome. A permutation test confirmed this over-representation of OR genes within inversions (p = 0.03). In addition, several *Ips*-specific ORs (5 out of 7 ORs from an *Ips*-specific OR clade (Hou et al. 2021)) were located in inverted regions, specifically on IpsContig13 (4 out of 7 ORs). These 5 ORs (ItypOR23, ItypOR27, ItypOR28, ItypOR29, and ItypOR49) have been functionally characterized, responding to compounds primarily produced by beetles (pheromones), the host tree, or fungal symbionts (Hou et al. 2021; Powell et al. 2021). We found no difference in the OR number (no OR deletions) between inversion haplotypes. The majority of ORs located in inversions harbor multiple nonsynonymous variants segregating within the spruce bark beetle populations (Table S4) and nine ORs (Table S5) had at least one fixed or nearly fixed (*d_xy_*>0.9) nonsynonymous variant between inversion haplotypes. These diverged ORs were located at Inv5 (ItypOR43JOI, ItypOR29), Inv13 (ItypOR23, ItypOR27), Inv14.1 (ItypOR20NTE, ItypOR36) and Inv16.1 (ItypOR30, ItypOR31, ItypOR16). Among these ORs only ItypOR30 had dN/dS >1 (2 nonsynonymous substitutions, McDonald-Kreitman test p=0.55).

Two inversions harboring multiple OR genes showed significant latitudinal variation (Figure 3; Inv13 and Inv16.1+23.1, Table 1). Olfactory receptor activity and odorant binding were also among the enriched gene ontology categories in inversions that showed a correlation between inversion haplotype frequency and latitude (Figure 3, Table S6; 0.01 < p < 0.05). In *Drosophila pseudoobscura* odorant receptors were also associated with inversions (Fuller et al. 2016; Fuller et al. 2017). Interestingly, one of the inversions, Inv13, includes a gene encoding ItypOR23, receptor that has been previously shown to primarily respond to an odor from fungi (Hou et al. 2021). This OR was also among the most divergent between alternative inversion haplotypes (Table S5). We hypothesize that different OR alleles associated with these inversions may be involved in spruce bark beetle interactions with fungal associates present in different parts of Europe. As many other bark beetle species, *I. typographus* is associated with several, spatially differentiated, symbiotic fungal species that can help beetles to exhaust and overcome tree’s defense system (Lieutier et al. 2009). Interestingly, recent studies suggested that beetles can recognize volatile organic compounds of specific beneficial fungi that can be located and picked up and that individual beetles may also have preferences for different fungal species (Kandasamy et al. 2019; Zhao et al. 2019; Kandasamy et al. 2023).German spruce bark beetles have, for example, been shown to be more attracted to fungal species that are common in Germany (*Grosmannia penicillata, Endoconidiophora polonica*) than to rarer species (*Leptographium europhioides*). Unpublished data from Swedish beetles suggests that they are more attracted to *L. europhioides*, which is common in Sweden (personal communication D. Kandasamy). Although preliminary, these observations suggest that beetle preferences may be tuned to the local fungal community, and we speculate that inversions may be involved in recognition of region-specific fungal species.

Several other interesting behavioral strategies are polymorphic among spruce bark beetle individuals, including the existence of pioneer individuals that are the first to infest host trees, and re-emergence of females after egg laying to establish so-called sister broods in new trees. Other less obvious/visible phenotypes could also be associated with inversion polymorphisms. Gene ontology enrichment analysis provided a list of categories of interest (Table S7) but comprehensive research, both bottom-up and top-down, is needed to understand the relationship between spruce bark beetle phenotypes and the inversion polymorphism landscape.

### Evolutionary mechanisms maintaining inversion polymorphism in the spruce bark beetle

Several not mutually exclusive evolutionary mechanisms can maintain polymorphic inversions within species, in particular divergent and balancing selection (Wellenreuther and Bernatchez 2018; Faria, Johannesson, et al. 2019). The importance of divergent selection has been postulated based on allele frequency patterns and associations of polymorphic inversions with local adaptations that persist despite extensive intraspecific gene flow (Tigano and Friesen 2016). For example, a recent study of deer mice (*Peromyscus maniculatus*) (Harringmeyer and Hoekstra 2022) identified multiple polymorphic inversions with clinal variation across environmental gradients in two distinct habitats. Such frequency changes have also been reported across hybrid zones (Faria, Chaube, et al. 2019) or latitudinal gradients (Mérot et al. 2021). Although the spruce bark beetle does not occupy distinct environmental niches, it inhabits forests across a very wide latitudinal gradient (spanning at least 16 degrees). It is also a species with high dispersal capacity and extensive gene flow, as indicated by low *F_ST_*across its range. We found a significant correlation between the frequency of inversion haplotypes in populations and geographic location (latitude) for five inversions (Figure 3; r^2^ ranged from 0.34 to 0.68). There were no significant correlations between haplotype frequencies and longitude (Figure S11). In most cases, differences in inversion haplotype frequencies were small (Figure 3), except for the two inversions with the strongest correlations (r^2^ > 0.6; Inv12 and Inv13). Climate and land cover variation (summarized via PCA) was associated with allele frequency changes of multiple SNPs (59), however, no such association was found for inversion genotypes (Figure 4). The identified SNPs were in the proximity of 14 genes, five of which had assigned gene ontology categories (Table S8). All the above results indicated that probably there is selection across environmental gradients in the spruce bark beetle but that this is not the only, or even a major, force maintaining inversion polymorphism within the species.

**Figure 4.**
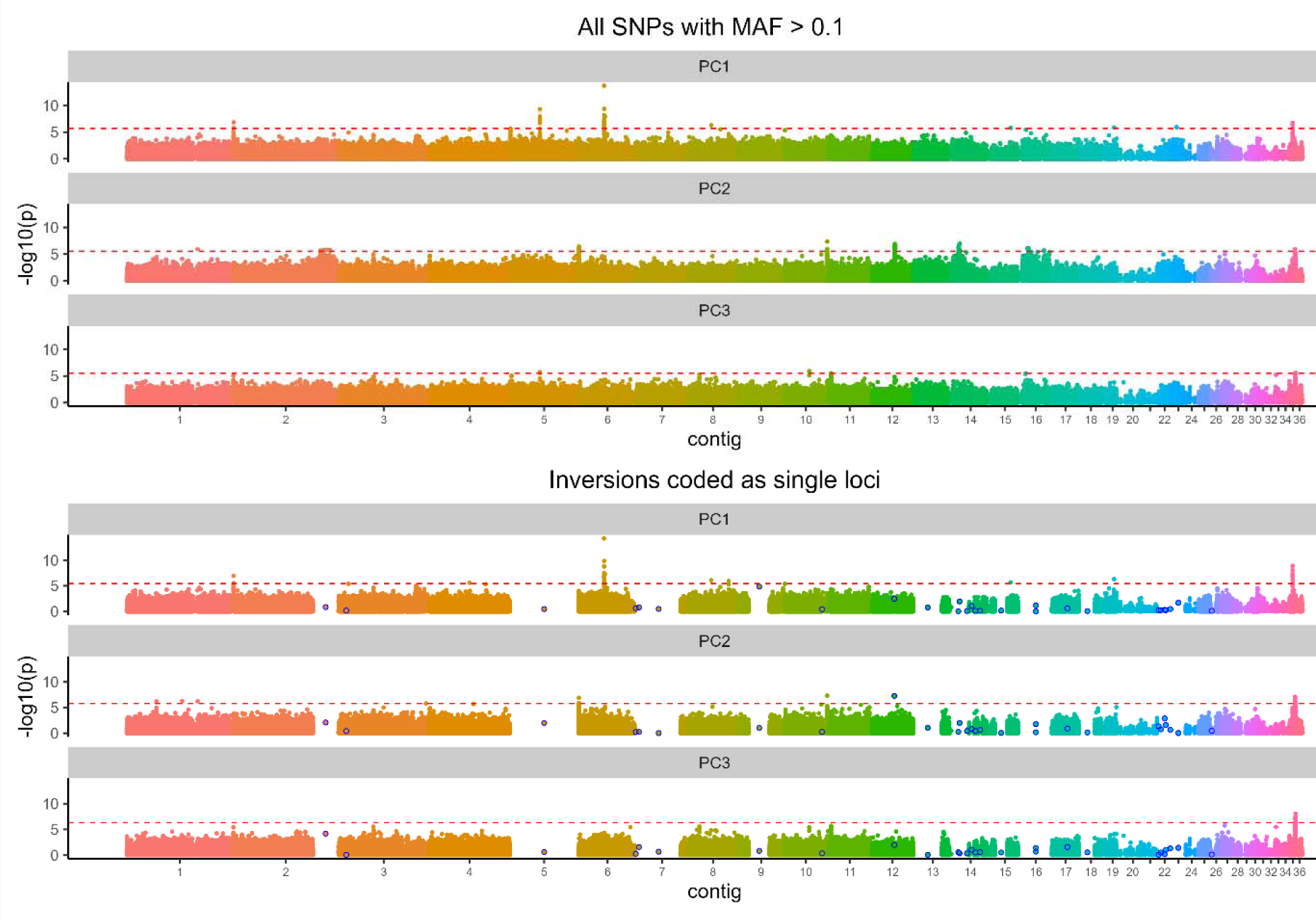
Genotype-environment associations across 36 contigs in the *Ips typographus* genome. Different colors represent different contigs. Each dot represents a single SNP. The upper three panels show results for all SNPs and the lower three panels show results using data where each inversion was coded as a single locus (shown in a blue circle). MAF: Minor Allele Frequency. PC1, PC2, and PC3 represent the first three of PCA used to summarize the environmental variation among populations.

While the ‘local adaptation’ is a major hypothesis proposed to explain inversion polymorphism (Christmas et al. 2019; Akopyan et al. 2022; Harringmeyer and Hoekstra 2022), balancing selection and related mechanisms may also be important in maintaining polymorphic arrangements (Faria, Johannesson, et al. 2019; Mérot et al. 2020; Berdan et al. 2021). Such mechanisms include overdominance, associative overdominance, frequency-dependent selection, and spatially and temporally varying selection. We found no support for overdominance playing a role in the spruce bark beetle, as no excess of inversion heterozygotes compared to Hardy-Weinberg expectations was detected in any of the populations. The only significant deviations from Hardy-Weinberg expectations were detected in Inv12 and Inv22.4 in geographic regions and/or across the whole species range (Table S9). We therefore looked more closely at mutation load, which can say something about the role of associative overdominance in the maintenance of inversion polymorphism. Theory predicts that recessive deleterious mutations will accumulate on both inversion arrangements but that most of these mutations will be private to only one arrangement (Navarro et al. 2000; Faria, Johannesson, et al. 2019; Berdan et al. 2021). This would lead to associative overdominance, as in heterozygotes the effects of deleterious recessive alleles on one arrangement would be masked by the wild-type alleles on the other arrangement. The result would be long-term maintenance of the inversion polymorphism, resulting in strong divergence between inversion haplotypes (Navarro et al. 2000; Guerrero et al. 2012; Berdan et al. 2021).

Interestingly, several stable evolutionary scenarios that maintain polymorphic inversions are possible (for details see Figure 4 in (Berdan et al. 2021)). These scenarios differ in the expected mutation load, fitness, and frequency of the corresponding genotypes. Given the haplotype frequencies we observed in spruce bark beetle inversions, two scenarios are likely. First, that minority arrangements experience higher mutation load (due to reduced recombination and lower population size) but are maintained in the population at low frequency due to, e.g., associative overdominance. Such a mechanism would favor balanced inversion polymorphisms of intermediate to large sizes (harboring hundreds of genes; (Ohta 1971; Connallon and Olito 2022) and has been shown to play a role in maintaining polymorphic inversions in several insect species (Yang et al. 2002; Jay et al. 2021). Second, mutation load may accumulate on one or both inversion arrangements but be mitigated by the haplotype structuring process, i.e., the existence of multiple diverged sub-haplotypes among inversion homozygotes that reduces the mutation load within homozygotes. If this process operates within one or both inversion arrangements it may result in more equal frequencies of alternative inversion haplotypes. However, such a mechanism is only possible when genetic variation and mutation load is high (Berdan et al. 2021).

In contrast to these theoretical expectations, we observed no sign of increased mutation load (measured by the π_N_/π_S_ ratio) in inversion regions compared to the collinear part of the spruce bark beetle genome (Table S10; Figure 5). We also found no sign of haplotype structuring capable of reducing mutation load within inversion homozygotes (Figure S12). The only significant within-homozygote clustering we observed was in a few inversion haplotypes.

**Figure 5.**
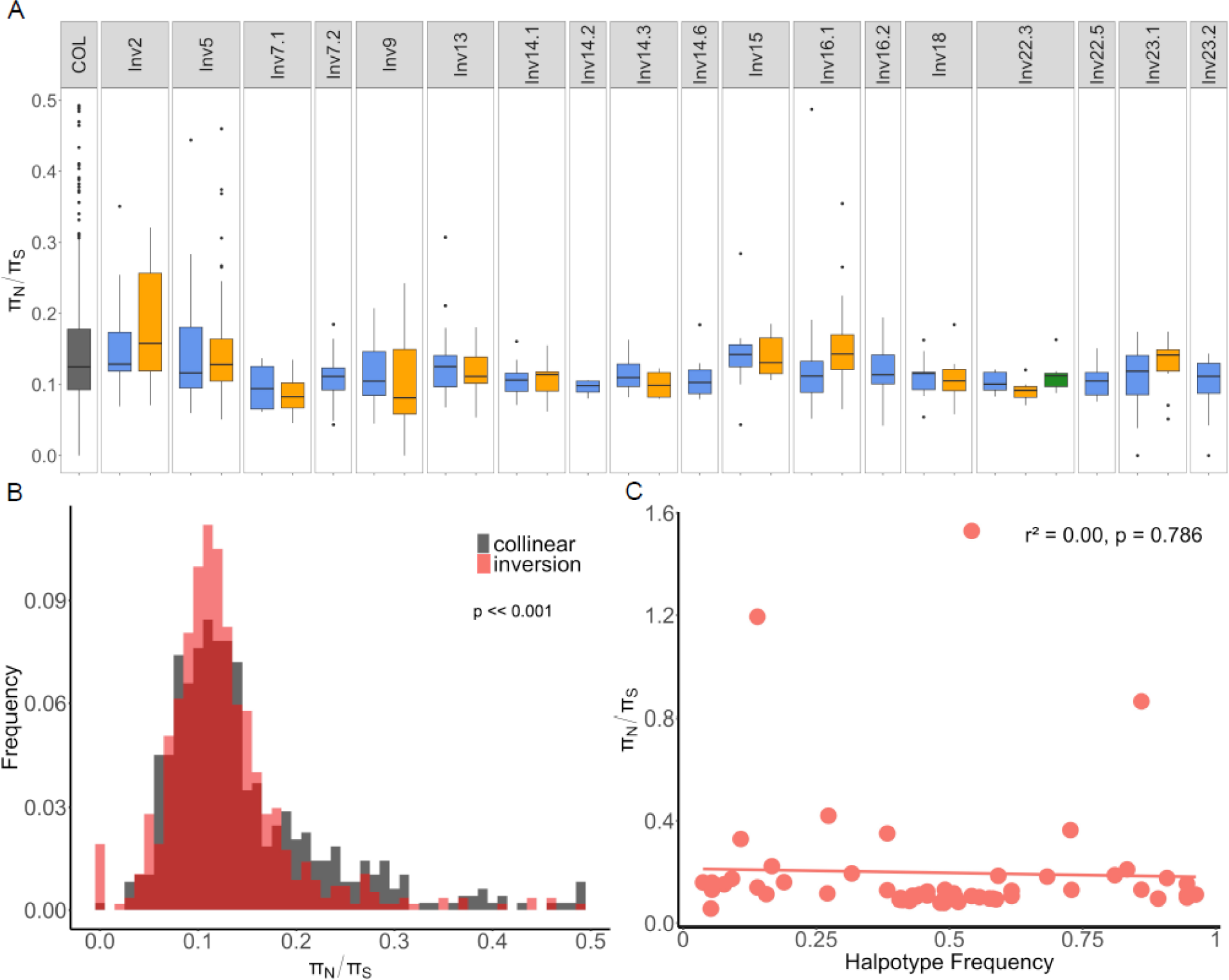
Mutation load analysis in *Ips typographus*. (A) The ratio of nonsynonymous to synonymous nucleotide diversity π_N_/π_S_computed for 200 kb windows along collinear parts of the genome (COL) and different inversion haplotypes (blue: major haplotype; yellow: minor haplotype; green: haplotype produced by double crossover events between minor and major haplotypes). n = 4 to 55 windows per inversion haplotype and n = 531 for the collinear genome. Inversion haplotypes that were present in less than four individuals, did not include four or more 200 kb windows or included fewer than five genes were not included in the analysis. For better visibility, π_N_/π_S_ outliers above 0.5 are not shown (a total of 58 π_N_/π_S_ values, 45 of them in COL). The lower and upper box hinges correspond to the first and third quartiles, whiskers show 1.5 * the inter-quartile range. (B) Distribution of π_N_/π_S_ values per 200 kb window (C) Correlation between mean π_N_/π_S_ for all inversion haplotypes and their frequency across spruce bark beetle populations.

However, the clustering divided individuals into southern and northern clades suggesting either neutral differentiation or divergent selection acting on one of the inversion arrangements, rather than haplotype structuring being associated with mutation load (Figure S12). These results are consistent with observations of no significant mutation load in other species where inversion haplotypes are subject to divergent selection that results in geographic structuring (Harringmeyer and Hoekstra 2022; Huang et al. 2022). In such cases, inversions facilitate adaptive divergence but do not tend to accumulate a mutation load. However, geographic clustering of inversion polymorphisms is weak in the spruce bark beetle, and many inversions appear to have a slightly lower mutation load associated with the more common inversion haplotype (Figure 5, Table S11). It is possible that accumulation of a mutation load in the spruce bark beetle is mitigated by high effective population size in this species and/or gene conversion and double crossover events, which despite their apparent rarity have been detected in several spruce bark beetle inversions.

Overall, the absence of heterozygote excess does not support a role of overdominance in the maintenance of inversion polymorphism in the spruce bark beetle. Likewise, the absence of an elevated mutation load in inverted regions does not support a role of associative overdominance either. However, since we only have genomic data available, we cannot conclusively rule out that other potential mechanisms, such as negative frequency-dependent selection or antagonistic pleiotropy, could maintain balanced inversion polymorphisms. Additional temporal data are needed to test whether temporally varying selection has affected the frequencies of inversion haplotypes in the spruce bark beetle and further research is essential to determine the role of different mechanisms maintaining inversion polymorphisms within the species.

### Far-reaching consequences of having an inversion-rich genome

The presence of multiple polymorphic inversions can have significant consequences for the evolution of a species, as well as for evolutionary inferences based on genome-wide polymorphism data. Importantly, polymorphic inversions are a reservoir of genetic variation that can facilitate adaptation to rapidly changing environments. Indeed, several studies have shown that polymorphic inversions support rapid adaptation to changing climatic conditions (Rane et al. 2015; Kapun and Flatt 2019; McCulloch and Waters 2023) or adaptive tracking of fluctuating environments (Nunez et al. 2023). Spruce bark beetle populations are subject to seasonal weather changes and a rapidly changing environment due to strong anthropogenic pressures (Hofmann et al. 2024). Warmer weather and drought periods have been associated with a predicted intensification of bark beetle outbreaks (Biedermann et al. 2019; Bentz et al. 2021; Hlásny et al. 2021), which may act as a strong selection factor within bark beetle populations. Whether inversions are involved in rapid adaptations in the spruce bark beetle is an open question that requires further investigation.

Abundant polymorphic inversions within the genome can have far-reaching consequences for inferences about e.g. demographic history and selection. It is well known that non-equilibrium demography and selection can leave similar genomic signatures. Traditionally, demographic analyses have used non-coding parts of the genome, based on the assumption that directional selection mostly affects protein-coding regions. However, growing evidence for the importance of background selection in shaping genome-wide diversity is moving the field towards incorporating linked selection into inferences of demographic history (Li et al. 2012; Johri et al. 2021). We believe that new approaches should also consider the potential influence of polymorphic inversion landscapes, because variation patterns of inverted regions can be shaped by different types of balancing selection. In addition, genomics scans for selection in inversion-rich genomes may be biased due to reduced recombination within inversion regions. Importantly, the effect of reduced recombination may extend outside inversions (Adrion et al. 2020; Koury 2023; Li et al. 2023). The influence of inversions and inversion-rich genomes on various evolutionary inference methods may become a standard scenario to test their robustness (Novo et al. 2023).

## Materials and methods

### Species genome and its quality

The spruce bark beetle karyotype is n = 14 + XYp with male being a heterogametic sex. The species genome assembly consists of 272 contigs, and 36 largest (>1Mb) contigs constitute the majority of the assembly (78% of 236.8 Mb long assembly, N50 = 6.65 Mb; Powell et al. 2021). Telomeric motifs (sequences present at the end of chromosomes) were identified at the end of eight contigs (including five largest contigs). BUSCO analysis indicated that 99.5 % genes present in the insect database (insects_odb9) were also present among the predicted *Ips typographus* genes. To further investigate the quality of the *Ips typographus* assembly we aligned it to *Ips nitidius* reference genome (chromosome-level assembly, 16 chromosomes; 2.3 Mya divergence from *Ips typographus*; (Wang et al. 2023). The alignment was conducted with minimap2 (Li 2018; *asm20* preset recommended for sequences expected to differ in up to 20%). We used *Ips typographus* contigs as either an alignment target or as a query.

### Sampling

Adult spruce bark beetles were collected with pheromone-baited traps in the spring and summer of 2020. In total, we sampled 18 populations throughout Europe with 13-14 individuals per locality (253 individuals in total) (Figure 2; Table S5). Throughout the text, we use the term ‘population’ to refer to a particular site or a collection of closely situated sites (within about 50 km). In Austria, we pooled individuals from three localities that were up to 120 km apart because of small sample sizes (five beetles or less per site). Populations from Fennoscandia are referred to as northern populations (or the northern group) and populations from central Europe will be referred to as southern populations (or the southern group). Polish populations are considered separately from other central European populations (due to high admixture proportions from northern group identified in downstream analysis). Beetles were brought alive to the laboratory, kept on a paper diet for several days, dissected, sexed based on genitalia morphology, and subjected to DNA extraction (described below).

### DNA extraction and genome re-sequencing

DNA was extracted from the whole body of dissected beetles using the Wizard Genomic DNA Purification Kit (Promega). The concentration of extracted DNA was estimated using a Qubit fluorimeter (Thermo Fisher Scientific). Genomic libraries were prepared with NEBNext Ultra II FS DNA Library Prep with Beads (New England Biolabs), with single indexes. Individual libraries were combined into three pools and 2×150 bp paired-end sequenced in three lanes of a S4 flowcell using the NovaSeq 6000 instrument and v1 sequencing chemistry (Illumina Inc.). Sequencing was done by the National Genomics Infrastructure, SNP&SEQ Technology Platform (Uppsala, Sweden). To assess the overall genotyping error, we prepared and sequenced duplicate libraries for nine individuals.

### Data preparation and filtering

Details of raw data processing and filtering are described in the Supplementary Files. Shortly, raw reads were mapped to the reference genome (Powell et al. 2021) using Bowtie 2 version 2.4.2 (Langmead and Salzberg 2012). Duplicated reads were removed using Picard MarkDuplicates version 2.24.1 (Broad Institute 2019). To detect and correct systematic errors in base quality scores, recalibration was done using the Genome Analysis Toolkit (GATK) version 4.1.9.0, BaseRecalibrator, and ApplyBQSR (McKenna et al. 2010; Depristo et al. 2011). Variant calling and genotyping was done using GATK HaplotypeCaller, CombineGVCFs, and GenotypeGVCFs. GATK VariantRecalibrator and ApplyVQSR were used to calculate and filter (by variant) quality score log-odds (VQSLOD). Bcftools version 1.11 (Danecek et al. 2021) was used to remove insertions and deletions (indels) as well polymorphisms five bases up- and downstream. GATK VariantFiltration was applied to mask all genotypes with low sequencing depth or low genotype quality (McKenna et al. 2010; Depristo et al. 2011). Variants which were not biallelic single nucleotide polymorphisms or did not meet the recommended hard filtering thresholds (GATK Team, see Supplementary materials) were filtered out. To filter out polymorphisms that could come from duplicated regions we removed variants located within repeat-masked regions of the genome (Powell et al. 2021), variants with excessive overall coverage (mean + 1 SD), and variants with heterozygote excess. Variants for which genotypes could be detected in less than half of the individuals were also removed. We used PLINK version 1.90b6.18 (Purcell et al. 2007) to detect sample contaminations, swaps and duplications, and familial relationships (e.g. sibling pairs present in the data) which might bias downstream analyses. Individuals with excessive coverage were removed, as these could be a result of human errors during library preparation or pooling. We focused on contigs longer than 1 Mb that together constituted 78% of the genome assembly, i.e. a total of 186 Mb. Since part of IpsContig33 had high similarity to mtDNA this contig was not included in the downstream analysis. Genotyping error was assessed using GATK Genotype Concordance.

### Genome-wide genetic variation and its geographic structuring

Genome-wide genetic structuring was explored by PCA using PLINK. The most likely number of genetic clusters and admixture proportions was estimated using NGSadmix (Skotte et al. 2013). The analysis was run for five different K-values (1-5; 10 replicates per K-value), using a minor allele frequency (MAF) filter of 0.05 and 10,000 iterations, and a SNP dataset that was pruned for linkage disequilibrium using PLINK (–indep-pairwise 50 10 0.1 option). To choose the most likely number of genetic clusters, the results were examined using CLUMPAK (http://clumpak.tau.ac.il/index.html). To examine the influence of inversions on genetic clustering and to facilitate inversion genotyping, NGSadmix was run separately for 1) all autosomal contigs without potential inversions and 2) each potential inversion.

To assess genetic differentiation among different spruce bark beetle populations Weir and Cockerham’s (Weir and Cockerham 1984) *F*_ST_ was estimated using VCFtools version 0.1.16 (Danecek et al. 2011). *F*_ST_ was calculated between population groups identified by NGSadmix, as well as among all population pairs. Additionally, we summarized *F*_ST_ values in 100 kb overlapping windows (using 20 kb steps) along the contigs. Window-based analyses were done for each contig and each population pair. In addition, absolute sequence divergence (*d*_xy_), nucleotide diversity, and Tajima’s D statistic were estimated and summarized for 100 kb non-overlapping windows using ANGSD version 0.935-44-g02a07fc (Korneliussen et al. 2014).

These statistics were calculated for each population separately and were based on a maximum likelihood estimate of the folded site frequency spectrum (SFS). We excluded sites with mapping quality below 30 phred, base quality score below 20, and coverage less than three times the population sample size and more than three times the average coverage, following the approach used in Delmore et al. (Delmore et al. 2018). The ANGSD calculations were based on allele frequencies estimated from genotype likelihoods (Li 2011) and ngsPopGen scripts (https://github.com/mfumagalli/ngsPopGen).

### Identification of inversions, their geographic distribution, and variation patterns

To identify inversions, we followed standard population genetic approach (see e.g. Huang et al. 2020; Mérot et al. 2021; Reeve et al. 2023). Potential chromosomal inversion regions were identified based on: i) per contig PCAs, ii) local PCAs, iii) patterns of heterozygosity, and iv) LD clustering. Per contig PCAs were performed as a first screen for inversion existence using PLINK (Purcell et al. 2007). Local PCAs were performed to identify inversions approximate positions using the *lostruct* R package (Li and Ralph 2019). Analysis was performed using two independent approaches: localPCA analysis using all analyzed contigs and localPCA analysis for each contig independently. The first analysis was run following the approach described in Huang et al. (2020)Windows were considered outliers if their MDS scores were greater than 2SD from the mean across all windows and the maximum number of windows between what was considered separate inversions was set to 20. In the second analysis separate contigs were subject to k-means clustering algorithm (R) to define groups of windows that formed a single inversion. K-means clustering algorithm was performed on the first MDS scores. For each contig different k (number of window clusters) were tested so that each structurally different part of the contig was isolated into one cluster. For both approaches, different window sizes (1, 10 and 100kb) were tested.

Linkage disequilibrium among SNPs (thinned by selecting one SNP every 10 kb; MAF > 5%) was calculated for each contig using PLINK. We considered a genomic region to be an inversion region if 1) local PCA analysis identified the region as an outlier, and/or 2) the region exhibited high LD (most SNPs having r^2^ > 0.4), and 3) PCA performed on SNPs from this region separated individuals into three distinct groups with heterozygosity patterns matching the expectation of the middle group of the PCA presenting higher heterozygosity.

Genotyping of individual beetles with respect to the inversion haplotypes they carried was done based on NGSadmix clustering (with K = 2 inversion heterozygotes having mixed ancestry in approximate 50/50 proportions; Figure S13). In a few more complex cases (putative overlapping inversions and double-crossover events) genotyping was done by clustering individuals based on the PCA groups visible along PC1 and PC2 (for more details see Figure S3 and related text). Contigs with less than 10,000 SNPs provided ambiguous results and were excluded from genotyping. Approximate inversion boundaries were defined based on local PCAs and sharp borders detected in LD clusters. The population genetic approach used in this study is a standard approach to detect putative large polymorphic inversions but is not able to provide details on inversion breakpoints.

To further support inversion evidence, we estimated population recombination rates (rho) for each putative inversion region using individuals with a particular inversion genotype (separately for each homozygous and heterozygous individuals). We followed the approach used in Jones et al. (2019) and sampled 20 individuals from each genotype group in cases when more individuals were available. Rho was not calculated if there were less than 10 individuals of a particular genotype. Watterson theta estimates were used to create a custom likelihood lookup table using the Ldhat 2.2 program *complete* (McVean et al. 2002). The *interval* program was used to estimate the population recombination rate across investigated inversion (and adjacent regions in the same contig). The *interval* algorithm was run for 2 million iterations and the chain was sampled every 10,000 iterations with a burn-in of 100,000 generations (*stat* program in Ldhat package, block penalty = 5). Population recombination rates were summarized in non-overlapping 100 kb windows using a custom perl script.

Inversion genotype and haplotype frequencies were calculated using an in-house R-script. Frequencies were calculated 1) within each population, 2) within the southern and northern group, and 3) for all sampled populations combined. Deviations from Hardy-Weinberg equilibrium were estimated for all three datasets. Inversions with only two haplotypes were tested using Fisher’s exact tests. Inversions with more than two haplotypes (including recombinant haplotypes between two inversion arrangements) were tested using permutation test, and sex chromosome inversions were tested as described in Graffelman & Weir (Graffelman and Weir 2018). All tests were done using the R package *HardyWeinberg* (Graffelman 2015). To investigate if inversion haplotypes differed in frequency along environmental gradients, Pearson correlation between inversion haplotype frequencies and latitude/longitude was calculated. To assess levels of genetic differentiation between inversion haplotypes, *F*_ST_ and *d*_xy_ between alternative inversion haplotypes (AA, BB) were estimated following the approach described above. Deletions present in alternative inversion haplotypes were not included in *d*_xy_ calculations but were summarized separately using custom scripts.

### Testing for enrichment of gene ontology terms in inversions

We used genes annotations from Powell et. al (2021) and tested for overrepresentation of gene ontology (GO) terms among genes located within inversions using R package topGO (Alexa and Rahnenführer 2022), applying Fisher’s exact test and “weight01” algorithm (Alexa et al. 2006) to deal with the GO graph structure; only GO categories with at least 10 members among the SNP associated genes were considered.

### Inversion age estimation

Absolute sequence divergence between alternative inversion haplotypes was used to calculate the approximate time of divergence of inverted and non-inverted haplotypes. We used the equation T = *d_xy_*/2µ, where T is the divergence time in generations, µ is the mutation rate per site per generation, and *d_xy_* is a mean *d_xy_* calculated based on per SNP values estimated in ANGSD. Since mutation rates of the spruce bark beetle are unknown, we used a range of mutation rate estimates available for some diploid, sexually reproducing insects including *Drosophila melanogaster* (Krasovec 2021)*, Heliconius melpomene* (Keightley et al. 2015), and *Chironomus riparius* (Oppold and Pfenninger 2017). Per generation per bp mutation rate estimates varied from 2.1 to 11.7 × 10^-9^. This approach could only give us rough inversion age estimates due to the uncertainty of the mutation rate estimates, probable intraspecific variation in mutation rate (Krasovec 2021), and a (likely) substantial influence of selection and gene flux (Charlesworth 2023).

### Mutation load estimation

To estimate mutation load we calculated the ratio of nucleotide diversity at non-synonymous sites (π_N_) vs. synonymous sites (π_S_). Mutation load (π_N_/π_S_) was calculated separately for each inversion homozygote and for the collinear part of the genome (all individuals). We computed nucleotide diversity for each site using SNPGenie (Nelson et al. 2015). The π_N_/π_S_ ratio was estimated in 200 kb windows using an in-house R script. To account for the fact that inversions can greatly suppress recombination in surrounding parts of the genome (Koury 2023) the collinear part of the genome was divided into two groups: 1) a group including all collinear 200 kb windows outside inversions and 2) a group including all collinear windows outside inversions but excluding windows that came from contigs with inversions (so called strict filtering). Both collinear datasets were used to test for overall differences in mutation load between inversions and the collinear part of the genome (using two-sided t-test). One-sided t-tests were used to test whether minor (less frequent) homokaryotypes had higher mutation loads than major (more frequent) homokaryotypes. Homokaryotypes that contained fewer than four 200 kb windows and were present in few individuals (two thresholds were tested: < 4 and < 10 individuals) were excluded from the analysis. Additionally, windows with a small number of genes were excluded (two thresholds were tested: < 5 and < 10 genes).

Haplotype structuring, i.e., the presence of two or more distinct sub-haplotypes among inversion homozygote haplotypes, can prevent fitness degeneration on one or both inversion haplotypes by carrying partially complementary sets of deleterious recessive alleles (Charlesworth and Charlesworth 1997; Berdan et al. 2021). To check if any inversion homozygotes exhibited haplotype structuring we first phased the data using Beagle 5.2 (default settings (Browning and Browning 2007)). Next, in each inverted region and homokaryotype we filtered out all variants with MAF < 0.1 and used PGDSpider 2.1.1.0 (Lischer and Excoffier 2012) to convert variant call formats (VCF) to full length sequences. Finally, we constructed neighbor-joining trees for alleles within haplotypes using MEGA7 and investigated if topologies showed the presence of distinct clusters within homokaryotype groups (Kumar et al. 2016).

### Phenotype-genotype associations

To test whether inversion polymorphisms were associated with diapause phenotypes we resequenced whole genomes of 18 individuals from a spruce bark beetle diapause study by Schebeck et al. (Schebeck et al. 2022) (10 beetles expressing facultative diapause and 8 beetles expressing obligatory diapause). DNA sequencing was performed using the DNBseq platform (BGI Tech Solutions, Poland) to a mean coverage of 20×. The data was processed in the same way as described above and combined with other sequenced individuals before performing PCA in PLINK. To test for differentiation in inversion haplotype frequencies between diapause phenotypes an exact G test was run using Genepop 4.1.2 (Raymond and Rousset 1995). *F*_ST_ between individuals expressing facultative and obligate diapause was estimated using VCFtools version 0.1.16 (Danecek et al. 2011) and summarized in 100 kb overlapping windows (20 kb steps).

To check if spruce bark beetle odorant receptors (ORs) genes were associated with inversions we examined 73 OR genes recently annotated by Yuvaraj et al. (Yuvaraj et al. 2021). OR sequences were mapped to the bark beetle reference genome using minimap2 (Li 2018), and 71 out of 73 ORs were located in the 186 Mb covered by the 36 contigs we analyzed. Three ORs mapped to more than one contig. ItypOR9 and ItypOR58 mapped to the end of IpsContig16 and 23, suggesting possible assembly error and duplication of end-of-contig sequences, which are difficult to assemble. ItypOR59NTE mapped to three nearby locations on IpsContig6, suggesting either assembly error or recent duplication. For these three ORs we used one randomly chosen location in downstream analyses. To test if inversions were enriched in OR genes, we ran permutation tests (10,000 iterations; permutating inversion locations). To check if alternative inversion arrangements harbored different numbers of OR genes (e.g. that one arrangement carried a deletion) we compared sequence coverage within ORs in individual beetles identified as inversion homozygotes. Additionally, we counted nonsynonymous segregating variants present in ORs situated within inversions and checked if any of these SNPs are highly diverged between inversion haplotypes (by calculating *d_xy_*). Finally, to investigate whether natural selection was involved in shaping ORs nonsynonymous variation patterns between inversion haplotypes we calculated d_N_/d_S_ ratio for ORs with multiple highly diverged nonsynonymous variants and performed MacDonald-Kreitman test for ORs with d_N_/d_S_ > 1. *D_xy_*was calculated using GATK VariantsToTable outputs and custom perl scripts and dN/dS was calculated using pairwise distances option in MEGA11 (Syn-Nonsynonymous substitution model, Kumar method). MacDonald-Kreitman test was performed in DnaSP v6 (Rozas et al. 2017).

### Genotype-environment association

Genotype-environment association (GEA) analyses were done to test if allele frequency changes in SNPs were associated with the beetle populations’ local environment. The analysis was performed using 1) all SNPs present in the analyzed part of the genome and 2) excluding SNPs present within inversion regions and coding inversions as single SNPs. Many different environmental variables were summarized along two principal components (see below). To control for the confounding effect of the overall genetic differentiation, we used Latent Factors Mixed Models (LFMM (Caye et al. 2019)) as implemented in the lfmm2() function from the R package LEA (Gain and François 2021). We used only SNPs with < 20% missing data and MAF >= 0.1, and that occurred in individuals with < 30% missing data. Because LFMM cannot handle missing data, we imputed missing genotypes with impute() from LEA. We ran LFMM for 1) all SNPs that passed the filters described above and 2) for a dataset where each inversion was represented as a single “SNP” inversion genotype. We used five latent factors (K = 5) in lfmm2(). P-values were calculated using lfmm2.test() from LEA and false discovery rate (FDR)-corrected using the p.adjust() R function with method = “fdr”.

Each beetle population’s local environment was characterized according to climate and land-cover data. We used all 19 bioclimatic variables from WorldClim version 2.1. (Fick and Hijmans 2017) with a resolution of ∼1 km^2^. Environmental variables included annual mean temperature, mean diurnal range, isothermality, temperature seasonality, maximum temperature of warmest month, minimum temperature of coldest month, temperature annual range, mean temperature of wettest quarter, mean temperature of driest quarter, mean temperature of warmest quarter, mean temperature of coldest quarter, annual precipitation, precipitation of wettest month, precipitation of driest month, precipitation seasonality, precipitation of wettest quarter, precipitation of driest quarter, precipitation of warmest quarter, and precipitation of coldest quarter. These variables are averaged over the years 1970-2000. Proportions of forest, cropland, and built-up areas were downloaded from https://lcviewer.vito.be/ for 2015 with a spatial resolution of ∼100 m^2^ (Buchhorn et al. 2019). These global land-cover maps are part of the Copernicus Land Service, derived from PROBA-V satellite observations, and have an accuracy of 80% as measured by the CEOS land product validation subgroup. We also included the proportion of land area covered by spruce trees (genus *Picea*) at a resolution of ∼1 km^2^, as obtained by Brus et al. (2012) using a statistical mapping approach. All environmental variables were reprojected to a final resolution of ∼1 km^2^ using the Lambert azimuthal equal area method. We then extracted mean values for all environmental variables (Figure S14) within a 50 km radius from each population location using the R package terra (Hijmans 2023). Finally, PCA was used to summarize the multi-scale environmental variation among populations. The first two PCA components (PC1 and PC2) explained 25% and 23% of the environmental variation, respectively, and were used as the final input for the GEA analyses. PC1 represented environmental variation mainly related to latitude, with northern populations showing higher values indicative of higher temperature variation and lower temperatures during the coldest months. PC2 represented environmental variation mostly related to temperature and amount of cropland, with higher values representing localities with higher temperatures during the warmest months and a higher proportion of cropland (and conversely less forest cover and spruce) (Figure S15). Additionally, we identified genes closest to SNPs that showed significant association with PC.

## Data Availability

All DNA sequences have been deposited to the National Center for Biotechnology Information Sequence Read Archive under the BioProject ID PRJNA1013983.

## Acknowledgments

We thank A. Hietala, E. Stengel, K. Zub, Å. Lindelöw, O. Langvall, M. Holmlund, E. Kristensen, U. Johansson, R. Modlinger, J. Reisenberger, K. Szreder, W. Skowroński L. Stanecki, and M. Ahlström for help in sampling spruce bark beetles across Europe. T. Mokrzycki helped with beetle sexing and identification. We thank members of the Genomics and Experimental Evolution Group at Jagiellonian University for their help in improving this manuscript. We thank Tomasz Gaczorek for the help in writing scripts and optimizing data analysis. This work was funded by a Polish National Science Center 2018/30/E/NZ8/00105 grant to K.N.B, and the Foundation in Memory of Oscar and Lili Lamm to M.N.A.

## Author contributions

A.M., P.Z. and K.N.B. conceived the study, performed the main analyses, and wrote the manuscript. A.B. managed sample shipment, sexed beetles, isolated DNA, and prepared samples for sequencing. F.S., P.K. and M.N.A. provided an unpublished version of the spruce bark beetle genome and valuable insights on the species’ ecology and sensory biology. M.M., J.M., Z.B., M.S., P.K., and C.S. organized sampling, provided beetles for analysis and insights on sampled populations. B.A. and W.B. performed GEA analysis. Z.N. analyzed diapause data. J.M. phased the data. M.S. provided diapause samples. J.Z performed *Ips nitidus* analysis. W.B. helped in data interpretation and provided feedback on all manuscript versions. All authors read and approved the final manuscript.

## References

Adrion JR, Galloway JG, Kern AD. 2020. Predicting the landscape of recombination using deep learning. Mol. Biol. Evol. 37:1790–1808.

Akopyan M, Tigano A, Jacobs A, Wilder AP, Baumann H, Therkildsen NO. 2022. Comparative linkage mapping uncovers recombination suppression across massive chromosomal inversions associated with local adaptation in Atlantic silversides. Mol. Ecol. 31:3323– 3341.

Alexa A, Rahnenführer J. 2022. topGO: Enrichment analysis for gene ontology (2.46. 0)[computer software]. Bioconductor version: Release (3.14)e.

Alexa A, Rahnenführer J, Lengauer T. 2006. Improved scoring of functional groups from gene expression data by decorrelating GO graph structure. Bioinformatics 22:1600–1607.

Ayala D, Acevedo P, Pombi M, Dia I, Boccolini D, Costantini C, Fontenille D. 2017. Chromosome inversions and ecological plasticity in the main African malaria mosquitoes. Evolution 71:686–701.

Ayala D, Ullastres A, González J, Chain FJJ, Planck M. 2014. Adaptation through chromosomal inversions in Anopheles. Front. Genet. 5:1–10.

Bentz BJ, Hansen EM, Davenport M, Soderberg D. 2021. Complexities in predicting mountain pine beetle and spruce beetle response to climate change. In: Bark beetle management, ecology and climate change. Elsevier.

Berdan EL, Barton NH, Butlin R, Charlesworth B, Faria R, Fragata I, Gilbert KJ, Jay P, Kapun M, Lotterhos KE, et al. 2023. How chromosomal inversions reorient the evolutionary process. J. Evol. Biol. 36:1761–1782.

Berdan EL, Blanckaert A, Butlin RK, Bank C. 2021. Deleterious mutation accumulation and the long-term fate of chromosomal inversions. PLoS Genet. 17: e1009411.

Bertheau C, Schuler H, Arthofer W, Avtzis DN, Mayer F, Krumböck S, Moodley Y, Stauffer C. 2013. Divergent evolutionary histories of two sympatric spruce bark beetle species. Mol. Ecol. 22:3318–3332.

Biedermann PHW, Müller J, Grégoire JC, Gruppe A, Hagge J, Hammerbacher A, Hofstetter RW, Kandasamy D, Kolarik M, Kostovcik M, et al. 2019. Bark Beetle Population Dynamics in the Anthropocene: Challenges and Solutions. Trends Ecol. Evol. 34:914–924.

Brelsford A, Purcell J, Avril A, Tran Van P, Zhang J, Brütsch T, Sundström L, Helanterä H, Chapuisat M. 2020. An Ancient and Eroded Social Supergene Is Widespread across Formica Ants. Curr. Biol. 30:304–311.e4.

Browning SR, Browning BL. 2007. Rapid and accurate haplotype phasing and missing-data inference for whole-genome association studies by use of localized haplotype clustering. Am. J. Hum. Genet. 81:1084–1097.

Brus DJ, Hengeveld GM, Walvoort DJJ, Goedhart PW, Heidema AH, Nabuurs GJ, Gunia K. 2012. Statistical mapping of tree species over Europe. Eur. J. For. Res. 131:145–157.

Buchhorn M, Smets B, Bertels L, Lesiv M, Tsendbazar, N. E., Herold M, Fritz S. 2019. Copernicus global land service: Land cover 100m: Epoch 2015: Globe. Version V2. 0.2.

Caye K, Jumentier B, Lepeule J, François O. 2019. LFMM 2: Fast and accurate inference of gene-environment associations in genome-wide studies. Mol. Biol. Evol. 36:852–860.

Charlesworth B. 2023. The effects of inversion polymorphisms on patterns of neutral genetic diversity. Genetics 224:1–21.

Charlesworth B, Charlesworth D. 1997. Rapid fixation of deleterious alleles can be caused by Muller’s ratchet. Genet. Res. 70:63–73.

Cheng C, Tan JC, Hahn MW, Besansky NJ. 2018. Systems genetic analysis of inversion polymorphisms in the malaria mosquito Anopheles gambiae. Proc. Natl. Acad. Sci. U. S. A. 115:E7005–E7014.

Christmas MJ, Wallberg A, Bunikis I, Olsson A, Wallerman O, Webster MT. 2019. Chromosomal inversions associated with environmental adaptation in honeybees. Mol. Ecol. 28:1358–1374.

Connallon T, Clark AG. 2014. Balancing selection in species with separate sexes: Insights from fisher’s geometric model. Genetics 197:991–1006.

Connallon T, Olito C. 2022. Natural selection and the distribution of chromosomal inversion lengths. Mol. Ecol. 31:3627–3641.

Danecek P, Auton A, Abecasis G, Albers CA, Banks E, DePristo MA, Handsaker RE, Lunter G, Marth GT, Sherry ST, et al. 2011. The variant call format and VCFtools. Bioinformatics 27:2156–2158.

Danecek P, Bonfield JK, Liddle J, Marshall J, Ohan V, Pollard MO, Whitwham A, Keane T, McCarthy SA, Davies RM. 2021. Twelve years of SAMtools and BCFtools. Gigascience 10:1–4.

Delmore KE, Lugo Ramos JS, Van Doren BM, Lundberg M, Bensch S, Irwin DE, Liedvogel M. 2018. Comparative analysis examining patterns of genomic differentiation across multiple episodes of population divergence in birds. Evol. Lett. 2:76–87.

Denlinger D. 2022. Insect diapause. Cambridge University Press.

Depristo MA, Banks E, Poplin R, Garimella K V., Maguire JR, Hartl C, Philippakis AA, Del Angel G, Rivas MA, Hanna M, et al. 2011. A framework for variation discovery and genotyping using next-generation DNA sequencing data. Nat. Genet. 43:491–501.

Ellerstrand SJ, Choudhury S, Svensson K, Andersson MN, Kirkeby C, Powell D, Schlyter F, Jönsson AM, Brydegaard M, Hansson B, et al. 2022. Weak population genetic structure in Eurasian spruce bark beetle over large regional scales in Sweden. Ecol. Evol. 12:1–12.

Faria R, Chaube P, Morales HE, Larsson T, Lemmon AR, Lemmon EM, Rafajlović M, Panova M, Ravinet M, Johannesson K, et al. 2019. Multiple chromosomal rearrangements in a hybrid zone between Littorina saxatilis ecotypes. Mol. Ecol. 28:1375–1393.

Faria R, Johannesson K, Butlin RK, Westram AM. 2019. Evolving Inversions. Trends Ecol. Evol. 34:239–248.

Fick SE, Hijmans RJ. 2017. WorldClim 2: new 1-km spatial resolution climate surfaces for global land areas. Int. J. Climatol. 37:4302–4315.

Fuller ZL, Haynes GD, Richards S, Schaeffer SW. 2016. Genomics of natural populations: How differentially expressed genes shape the evolution of chromosomal inversions in Drosophila pseudoobscura. Genetics 204:287–301.

Fuller ZL, Haynes GD, Richards S, Schaeffer SW. 2017. Genomics of natural populations: Evolutionary forces that establish and maintain gene arrangements in Drosophila pseudoobscura. Mol. Ecol. 26:6539–6562.

Fuller ZL, Koury SA, Phadnis N, Schaeffer SW. 2019. How chromosomal rearrangements shape adaptation and speciation: Case studies in Drosophila pseudoobscura and its sibling species Drosophila persimilis. Mol. Ecol. 28:1283–1301.

Gain C, François O. 2021. LEA 3: Factor models in population genetics and ecological genomics with R. Mol. Ecol. Resour. 21:2738–2748.

Graffelman J. 2015. Exploring diallelic genetic markers: The HardyWeinberg package. J. Stat. Softw. 64:1–23.

Graffelman J, Weir BS. 2018. Multi-allelic exact tests for Hardy–Weinberg equilibrium that account for gender. Mol. Ecol. Resour. 18:461–473.

Guerrero RF, Rousset F, Kirkpatrick M. 2012. Coalescent patterns for chromosomal inversions in divergent populations. Philos. Trans. R. Soc. B Biol. Sci. 367:430–438.

Gutiérrez-Valencia J, Hughes PW, Berdan EL, Slotte T. 2021. The Genomic Architecture and Evolutionary Fates of Supergenes. Genome Biol. Evol. 13:1–19.

Hansson BS, Stensmyr MC. 2011. Evolution of insect olfaction. Neuron 72:698–711.

Harringmeyer OS, Hoekstra HE. 2022. Chromosomal inversion polymorphisms shape the genomic landscape of deer mice. Nat. Ecol. Evol. 6:1965–1979.

Hijmans R. 2023. terra: Spatial Data Analysis. R package version 1.7–18.: https://CRAN.R-project.org/package=terra.

Hlásny T, König L, Krokene P, Lindner M, Montagné-Huck C, Müller J, Qin H, Raffa KF, Schelhaas MJ, Svoboda M, et al. 2021. Bark Beetle Outbreaks in Europe: State of Knowledge and Ways Forward for Management. Curr. For. Reports 7:138–165.

Hof AEV t., Campagne P, Rigden DJ, Yung CJ, Lingley J, Quail MA, Hall N, Darby AC, Saccheri IJ. 2016. The industrial melanism mutation in British peppered moths is a transposable element. Nature 534:102–105.

Hofmann S, Schebeck M, Kautz M. 2024. Diurnal temperature fluctuations improve predictions of developmental rates in the spruce bark beetle Ips typographus. J. Pest Sci. 2024.

Hou XQ, Yuvaraj JK, Roberts RE, Zhang DD, Unelius CR, Löfstedt C, Andersson MN. 2021. Functional Evolution of a Bark Beetle Odorant Receptor Clade Detecting Monoterpenoids of Different Ecological Origins. Mol. Biol. Evol. 38:4934–4947.

Huang K, Andrew RL, Owens GL, Ostevik KL, Rieseberg LH. 2020. Multiple chromosomal inversions contribute to adaptive divergence of a dune sunflower ecotype. Mol. Ecol. 29:2535–2549.

Huang K, Ostevik KL, Elphinstone C, Todesco M, Bercovich N, Owens GL, Rieseberg LH. 2022. Mutation Load in Sunflower Inversions Is Negatively Correlated with Inversion Heterozygosity. Mol. Biol. Evol. 39:1–17.

Jay P, Chouteau M, Whibley A, Bastide H, Parrinello H, Llaurens V, Joron M. 2021. Mutation load at a mimicry supergene sheds new light on the evolution of inversion polymorphisms. Nat. Genet. 53:288–293.

Jay P, Tezenas E, Véber A, Giraud T. 2022. Sheltering of deleterious mutations explains the stepwise extension of recombination suppression on sex chromosomes and other supergenes.

Johri P, Riall K, Becher H, Excoffier L, Charlesworth B, Jensen JD. 2021. The Impact of Purifying and Background Selection on the Inference of Population History: Problems and Prospects. Mol. Biol. Evol. 38:2986–3003.

Jones JC, Wallberg A, Christmas MJ, Kapheim KM, Webster MT, Singh N. 2019. Extreme Differences in Recombination Rate between the Genomes of a Solitary and a Social Bee. Mol. Biol. Evol. 36:2277–2291.

Joron M, Frezal L, Jones RT, Chamberlain NL, Lee SF, Haag CR, Whibley A, Becuwe M, Baxter SW, Ferguson L, et al. 2011. Chromosomal rearrangements maintain a polymorphic supergene controlling butterfly mimicry. Nature 477:203–206.

Kandasamy D, Gershenzon J, Andersson MN, Hammerbacher A. 2019. Volatile organic compounds influence the interaction of the Eurasian spruce bark beetle (*Ips typographus*) with its fungal symbionts. ISME J. 13:1788–1800.

Kandasamy D, Zaman R, Nakamura Y, Zhao T, Hartmann H, Andersson MN, Hammerbacher A, Gershenzon J. 2023. Conifer-killing bark beetles locate fungal symbionts by detecting volatile fungal metabolites of host tree resin monoterpenes.

Kapun M, Fabian DK, Goudet J, Flatt T. 2016. Genomic Evidence for Adaptive Inversion Clines in Drosophila melanogaster. Mol. Biol. Evol. 33:1317–1336.

Kapun M, Flatt T. 2019. The adaptive significance of chromosomal inversion polymorphisms in Drosophila melanogaster. Mol. Ecol. 28:1263–1282.

Keightley PD, Otto SP. 2006. Interference among deleterious mutations favours sex and recombination in finite populations. Nature 443:89–92.

Keightley PD, Pinharanda A, Ness RW, Simpson F, Dasmahapatra KK, Mallet J, Davey JW, Jiggins CD. 2015. Estimation of the Spontaneous Mutation Rate in Heliconius melpomene. Mol. Biol. Evol. 32:239–243.

Kim KW, De-Kayne R, Gordon IJ, Omufwoko KS, Martins DJ, Ffrench-Constant R, Martin SH. 2022. Stepwise evolution of a butterfly supergene via duplication and inversion. Philos. Trans. R. Soc. B Biol. Sci. 377.

Kirubakaran TG, Grove H, Kent MP, Sandve SR, Baranski M, Nome T, De Rosa MC, Righino B, Johansen T, Otterå H, et al. 2016. Two adjacent inversions maintain genomic differentiation between migratory and stationary ecotypes of Atlantic cod. Mol. Ecol. 25:2130–2143.

Koch EL, Morales HE, Larsson J, Westram AM, Faria R, Lemmon AR, Lemmon EM, Johannesson K. 2021. Genetic variation for adaptive traits is associated with polymorphic inversions in *Littorina saxatilis*. Evol. Letters 5:196–213.

Korneliussen TS, Albrechtsen A, Nielsen R. 2014. ANGSD: Analysis of Next Generation Sequencing Data. BMC Bioinformatics 15:1–13.

Koury SA. 2023. Predicting recombination suppression outside chromosomal inversions in Drosophila melanogaster using crossover interference theory. Heridity 130:196–208.

Kozak GM, Wadsworth CB, Kahne SC, Bogdanowicz SM, Harrison RG, Coates BS, Dopman EB. 2019. Genomic Basis of Circannual Rhythm in the European Corn Borer Moth. Curr. Biol. 29:3501–3509.e5.

Krasovec M. 2021. The spontaneous mutation rate of Drosophila pseudoobscura. G3 Genes, Genomes, Genet. 11: jkab151.

Kumar S, Stecher G, Tamura K. 2016. MEGA7: Molecular Evolutionary Genetics Analysis Version 7.0 for Bigger Datasets. Mol. Biol. Evol. 33:1870–1874.

Lamichhaney S, Fan G, Widemo F, Gunnarsson U, Thalmann DS, Hoeppner MP, Kerje S, Gustafson U, Shi C, Zhang H, et al. 2015. Structural genomic changes underlie alternative reproductive strategies in the ruff (*Philomachus pugnax*). Nat. Genet. 48:84–88.

Langmead B, Salzberg SL. 2012. Fast gapped-read alignment with Bowtie 2. Nat. Methods 9:357–359.

Li H. 2011. A statistical framework for SNP calling, mutation discovery, association mapping and population genetical parameter estimation from sequencing data. Bioinformatics 27:2987–2993.

Li H. 2018. Minimap2: Pairwise alignment for nucleotide sequences. Bioinformatics 34:3094– 3100.

Li H, Berent E, Hadjipanteli S, Galey M, Muhammad-Lahbabi N, Miller DE, Crown KN. 2023. Heterozygous inversion breakpoints suppress meiotic crossovers by altering recombination repair outcomes. PLoS Genet. 19:1–20.

Li H, Ralph P. 2019. Local PCA shows how the effect of population structure differs along the genome. Genetics 211:289–304.

Li J, Li H, Jakobsson M, Li S, Sjödin P, Lascoux M. 2012. Joint analysis of demography and selection in population genetics: Where do we stand and where could we go? Mol. Ecol. 21:28–44.

Lieutier F, Yart A, Salle A. 2009. Stimulation of tree defenses by Ophiostomatoid fungi can explain attack success of bark beetles on conifers. Ann. For. Sci. 66:801.

Lischer HEL, Excoffier L. 2012. PGDSpider: An automated data conversion tool for connecting population genetics and genomics programs. Bioinformatics 28:298–299.

Lohse K, Clarke M, Ritchie MG, Etges WJ. 2015. Genome-wide tests for introgression between cactophilic Drosophila implicate a role of inversions during speciation. Evolution 69:1178– 1190.

Lowry DB, Willis JH. 2010. A widespread chromosomal inversion polymorphism contributes to a major life-history transition, local adaptation, and reproductive isolation. PLoS Biol. 8:e1000500.

Matschiner M, Maria J, Barth I, Tørresen OK, Star B, Baalsrud HT, Servane M, Brieuc O, Pampoulie C, Bradbury I, et al. 2022. Supergene origin and maintenance in Atlantic cod. *Nat*. Ecol. Evol. 6:469–481.

Mayer F, Piel FB, Cassel-Lundhagen A, Kirichenko N, Grumiau L, Økland B, Bertheau C, Grégoire JC, Mardulyn P. 2015. Comparative multilocus phylogeography of two Palaearctic spruce bark beetles: Influence of contrasting ecological strategies on genetic variation. Mol. Ecol. 24:1292–1310.

McClung CE. 1905. The chromosome complex of orthopteran spermatocytes. Biol. Bull. 9:304– 340.

McCulloch GA, Waters JM. 2023. Rapid adaptation in a fast-changing world: Emerging insights from insect genomics. Glob. Chang. Biol. 29:943–954.

McKenna A, Hanna M, Banks E, Andrey Sivachenko KC, Kernytsky A, Garimella K, Altshuler D, Gabriel S, Daly M, DePristo MA. 2010. The Genome Analysis Toolkit: A MapReduce framework for analyzing next-generation DNA sequencing data. Genome Res. 20:1297– 1303.

McVean G, Awadalla P, Fearnhead P. 2002. A coalescent-based method for detecting and estimating recombination from gene sequences. Genetics 160:1231–1241.

Mérot C, Berdan EL, Cayuela H, Djambazian H, Ferchaud AL, Laporte M, Normandeau E, Ragoussis J, Wellenreuther M, Bernatchez L. 2021. Locally Adaptive Inversions Modulate Genetic Variation at Different Geographic Scales in a Seaweed Fly. Mol. Biol. Evol. 38:3953–3971.

Mérot C, Llaurens V, Normandeau E, Bernatchez L, Wellenreuther M. 2020. Balancing selection via life-history trade-offs maintains an inversion polymorphism in a seaweed fly. Nat. Commun. 11:1–11.

Navarro A, Barbadilla A, Ruiz A. 2000. Effect of inversion polymorphism on the neutral nucleotide variability of linked chromosomal regions in drosophila. Genetics 155:685–698.

Nelson CW, Moncla LH, Hughes AL. 2015. SNPGenie: Estimating evolutionary parameters to detect natural selection using pooled next-generation sequencing data. Bioinformatics 31:3709–3711.

Nilssen AC. 1984. Long-range aerial dispersal of bark beetles and bark weevils (Coleoptera, Scolytidae and Curculionidae) in northern Finland. Ann. Entomol. Fenn. 50:37–42.

Novo I, Ordás P, Moraga N, Santiago E, Quesada H, Caballero A. 2023. Impact of population structure in the estimation of recent historical effective population size by the software GONE. Genetics Selection Evolution 55:1–15.

Nunez JCB, Lenhart BA, Bangerter A, Murray CS, Mazzeo GR, Yu Y, Nystrom TL, Tern C, Erickson PA, Bergland AO. 2023. A cosmopolitan inversion facilitates seasonal adaptation in overwintering Drosophila. Genetics 226: iyad207.

Ohta T. 1971. Associative overdominance caused by linked detrimental mutations. Genet. Res. 18:277–286.

Oppold AM, Pfenninger M. 2017. Direct estimation of the spontaneous mutation rate by short-term mutation accumulation lines in Chironomus riparius. Evol. Lett. 1:86–92.

Paolucci S, Salis L, Vermeulen CJ, Beukeboom LW, van de Zande L. 2016. QTL analysis of the photoperiodic response and clinal distribution of period alleles in Nasonia vitripennis. Mol. Ecol. 25:4805–4817.

Porubsky D, Höps W, Ashraf H, Hsieh PH, Rodriguez-Martin B, Yilmaz F, Ebler J, Hallast P, Maria Maggiolini FA, Harvey WT, et al. 2022. Recurrent inversion polymorphisms in humans associate with genetic instability and genomic disorders. Cell 185:1986–2005.e26.

Powell D, Groβe-Wilde E, Krokene P, Roy A, Chakraborty A, Löfstedt C, Vogel H, Andersson MN, Schlyter F. 2021. A highly-contiguous genome assembly of the Eurasian spruce bark beetle, Ips typographus, provides insight into a major forest pest. Commun. Biol. 4:1–9.

Pruisscher P, Nylin S, Gotthard K, Wheat CW. 2018. Genetic variation underlying local adaptation of diapause induction along a cline in a butterfly. Mol. Ecol. 27:3613–3626.

Purcell J, Brelsford A, Wurm Y, Perrin N, Chapuisat M. 2014. Convergent genetic architecture underlies social organization in ants. Curr. Biol. 24:2728–2732.

Purcell S, Neale B, Todd-Brown K, Thomas L, Ferreira MAR, Bender D, Maller J, Sklar P, De Bakker PIW, Daly MJ, et al. 2007. PLINK: A tool set for whole-genome association and population-based linkage analyses. Am. J. Hum. Genet. 81:559–575.

Rane R V., Rako L, Kapun M, Lee SF, Hoffmann AA. 2015. Genomic evidence for role of inversion 3RP of Drosophila melanogaster in facilitating climate change adaptation. Mol. Ecol. 24:2423–2432.

Raymond M, Rousset F. 1995. An Exact Test for Population Differentiation. Evolution 49:1280.

Reeve J, Butlin RK, Koch EL, Stankowski S, Faria R. 2023. Chromosomal inversion polymorphisms are widespread across the species ranges of rough periwinkles (*Littorina saxatilis* and *L. arcana*). Early view.

Roesti M, Gilbert KJ, Samuk K. 2022. Chromosomal inversions can limit adaptation to new environments. 31:1–16.

Roff D a. 1996. The evolution of threshold traits in animals. Q. Rev. Biol. 71:3–35.

Rozas J, Ferrer-Mata A, Sanchez-DelBarrio JC, Guirao-Rico S, Librado P, Ramos-Onsins SE, Sanchez-Gracia A. 2017. DnaSP 6: DNA sequence polymorphism analysis of large data sets. Mol. Biol. Evol. 34:3299–3302.

Saitou M, Masuda N, Gokcumen O. 2022. Similarity-based analysis of allele frequency distribution among multiple populations identifies adaptive genomic structural variants. Mol. Biol. Evol. 39:1–19.

Sallé A, Arthofer W, Lieutier F, Stauffer C, Kerdelhué C. 2007. Phylogeography of a host-specific insect: Genetic structure of Ips typographus in Europe does not reflect past fragmentation of its host. Biol. J. Linn. Soc. 90:239–246.

Schaal SM, Haller BC, Lotterhos KE. 2022. Inversion invasions: When the genetic basis of local adaptation is concentrated within inversions in the face of gene flow. Philos. Trans. R. Soc. B Biol. Sci. 377.

Schebeck M, Dobart N, Ragland GJ, Schopf A, Stauffer C. 2022. Facultative and obligate diapause phenotypes in populations of the European spruce bark beetle Ips typographus. J. Pest Sci. 95:889–899.

Schwander T, Libbrecht R, Keller L. 2014. Supergenes and complex phenotypes. Curr. Biol. 24:R288–R294.

Skotte L, Korneliussen TS, Albrechtsen A. 2013. Estimating individual admixture proportions from next generation sequencing data. Genetics 195:693–702.

Stauffer C, Lakatos F, Hewitt GM. 1999. Phylogeography and postglacial colonization routes of Ips typographus L. (Coleoptera, Scolytidae). Mol. Ecol. 8:763–773.

Sturtevant AH. 1921. Linkage variation and chromosome maps. Proc. Natl. Acad. Sci. USA 7:181–183.

Thompson MJ, Jiggins CD. 2014. Supergenes and their role in evolution. Heredity (Edinb). 113:1–8.

Tigano A, Friesen VL. 2016. Genomics of local adaptation with gene flow. Mol. Ecol. 25:2144– 2164.

Tigano A, Jacobs A, Wilder AP, Nand A, Zhan Y, Dekker J, Therkildsen NO, Lohmueller K. 2021. Chromosome-Level Assembly of the Atlantic Silverside Genome Reveals Extreme Levels of Sequence Diversity and Structural Genetic Variation. Genome Biol. Evol. 13:1– 17.

Todesco M, Owens GL, Bercovich N, Légaré JS, Soudi S, Burge DO, Huang K, Ostevik KL, Drummond EBM, Imerovski I, et al. 2020. Massive haplotypes underlie ecotypic differentiation in sunflowers. Nature 584:602–607.

Wang J, Wurm Y, Nipitwattanaphon M, Riba-Grognuz O, Huang YC, Shoemaker D, Keller L. 2013. A Y-like social chromosome causes alternative colony organization in fire ants. Nature 493:664–668.

Wang Z, Liu Y, Wang H, Roy A, Liu H, Han F, Zhang X, Lu Q. 2023. Genome and transcriptome of Ips nitidus provide insights into high-altitude hypoxia adaptation and symbiosis. iScience 26:107793.

Weir BS, Cockerham CC. 1984. Estimating F-statistics for the analysis of population structure. Evolution 38:1358–1370.

Wellenreuther M, Bernatchez L. 2018. Eco-Evolutionary Genomics of Chromosomal Inversions. Trends Ecol. Evol. 33:427–440.

Wellenreuther M, Mérot C, Berdan E, Bernatchez L. 2019. Going beyond SNPs: The role of structural genomic variants in adaptive evolution and species diversification. Mol. Ecol. 28:1203–1209.

Yang YY, Lin FJ, Chang HY. 2002. Comparison of recessive lethal accumulation in inversion-bearing and inversion-free chromosomes in Drosophila. Zool. Stud. 41:271–282.

Yuvaraj JK, Roberts RE, Sonntag Y, Hou XQ, Grosse-Wilde E, Machara A, Zhang DD, Hansson BS, Johanson U, Löfstedt C, et al. 2021. Putative ligand binding sites of two functionally characterized bark beetle odorant receptors. BMC Biol. 19:1–21.

Zhao T, Kandasamy D, Krokene P, Chen J, Gershenzon J, Hammerbacher A. 2019. Fungal associates of the tree-killing bark beetle, Ips typographus, vary in virulence, ability to degrade conifer phenolics and in fl uence bark beetle tunneling behavior. Fungal Ecol. 38:71–79.

