## Supplemental Files for "Complex genomic landscape of inversion polymorphism in Europe’s most destructive forest pest"

### Details on resequencing, data preparation, and filtering

Raw data were first subjected to read quality control using FastQC 0.11.9 (Andrews 2010). Trimming of adaptors, low-quality bases and reads were done with Trimmomatic 0.39 (Bolger et al. 2014). Next, reads were mapped to the *Ips typographus* reference genome (Powell et al. 2021) using Bowtie 2 version 2.4.2 (Langmead and Salzberg 2012). Samtools 1.11 (Bonfield et al. 2021; Danecek et al. 2021) was used for alignment sorting and indexing. Mapping quality was assessed with Qualimap version 2.2.2 (Okonechnikov et al. 2016). To avoid allele amplification bias in variant calling, the Picard MarkDuplicates version 2.24.1 was used to remove duplicate reads (Broad Institute 2019). To detect and correct systematic errors in base quality scores recalibration was done using the Genome Analysis Toolkit (GATK) version 4.1.9.0 BaseRecalibrator and ApplyBQSR0 (McKenna et al. 2010; Depristo et al. 2011). Recalibration was performed separately for individual beetles sequenced on different lanes. This step provided recalibrated alignments which were used in the downstream analyses. A database of known variants was created from the same dataset using GATK hard filtering recommendations (GATK Team). Variants were filtered out when: QD < 2, FS > 60, MQ < 30, SOR > 3, MQRankSum < -12.5, ReadPosRankSum < -8, and, additionally for this purpose, we filtered out variants with MAF < 0.01 and > 50% missing genotypes. Variant calling and genotyping were performed with GATK HaplotypeCaller, CombineGVCFs, and GenotypeGVCFs. HaplotypeCaller was run in GVCF mode using default parameter settings, except for –heterozygosity (value used to compute prior probability that a locus is non-reference) which was set to 0.01 – since we expected increased levels of polymorphisms compared to the human genome. Next, variant quality score log-odds (VQSLOD) were calculated using GATK VariantRecalibrator and ApplyVQSR with the --truth-sensitivity-filter-level set to 99.0, to take into account properties of the variants not captured in the QUAL score. Bcftools version 1.11 (Bonfield et al. 2021; Danecek et al. 2021) was used to remove insertions and deletions (indels) as well as polymorphic sites five bases up- and downstream from indels. GATK VariantFiltration was applied to mask all genotypes with low sequencing depth (DP < 8) or low genotype quality (GQ < 20). Variants which were not biallelic single nucleotide polymorphisms or did not meet the hard filtering recommendations (see above) were filtered out. To filter out polymorphisms which may come from duplicated regions we removed variants located within masked regions of the genome (Powell et al. 2021), variants with excessive overall coverage (mean + 1 SD), and variants that exhibited significant heterozygote excess (ExcessHet > 54.69). Variants genotyped in less than half of the individuals were also removed. We used PLINK version 1.90b6.18 (--genome; Purcell et al. 2007) to detect sample contaminations, swaps, and duplications, as well as pedigree errors and unknown familial relationships (e.g. sibling pairs present in the data) which might bias downstream analyses. We also removed individuals with excessive coverage (coverage > mean*3) as these could have been affected by manual mistakes during library preparation or pooling. We focused on the 36 largest contigs (longer than 1 Mb) that constituted 78% of the genome assembly (i.e., 186 Mb). Since parts of IpsContig33 had high similarity to mtDNA this contig was not included in the downstream analyses. Genotyping error was assessed using GATK GenotypeConcordance.

After careful filtering of the whole genome re-sequencing data consisting of 253 libraries, we used 240 individuals in downstream analyses. Two individuals were removed due to high relatedness and another two due to extremely high sequencing coverage, indicating potential mistakes during library preparation. Nine individuals that had been sequenced twice allowed us to calculate genotyping errors, particularly Non-Reference Discrepancy which equaled 1.5% and Overall Genotype Concordance which was 99.8%. The mean coverage per individual varied from 5x to 53x (mean 23.2; median 23). Coverage at one contig (IpsContig9) was consistently lower (0.57 of average individual coverage) in individuals sexed as males (with only two mismatches (a 0.8% error rate) between morphologically identified sex and IpsContig9 coverage). Thus, IpsContig9 was treated as a sex (X) chromosome. After filtering we retained 5.245 million SNPs in the whole genome but we analyzed only 5.067 million SNPs present in the 35 analyzed contigs that constituted 75% of the whole genome assembly).

References


Andrews S. 2010. FastQC: a quality control tool for high throughput sequence data.

Bolger AM, Lohse M, Usadel B. 2014. Genome analysis Trimmomatic : a flexible trimmer for Illumina sequence data. 30:2114–2120.

Bonfield JK, Marshall J, Danecek P, Li H, Ohan V, Whitwham A, Keane T. 2021. HTSlib: C library for reading/writing high-Throughput sequencing data. *Gigascience* 10:1–6.

Cabanettes F, Klopp C. 2018. D-GENIES: Dot plot large genomes in an interactive, efficient and simple way. *PeerJ* 2018.

Danecek P, Bonfield JK, Liddle J, Marshall J, Ohan V, Pollard MO, Whitwham A, Keane T, McCarthy SA, Davies RM. 2021. Twelve years of SAMtools and BCFtools. *Gigascience* 10:1–4.

Depristo MA, Banks E, Poplin R, Garimella K V., Maguire JR, Hartl C, Philippakis AA, Del Angel G, Rivas MA, Hanna M, et al. 2011. A framework for variation discovery and genotyping using next-generation DNA sequencing data. *Nat. Genet.* 43:491–501.

Langmead B, Salzberg SL. 2012. Fast gapped-read alignment with Bowtie 2. *Nat. Methods* 9:357–359.

McKenna A, Hanna M, Banks E, Andrey Sivachenko KC, Kernytsky A, Garimella K, Altshuler D, Gabriel S, Daly M, DePristo MA. 2010. The Genome Analysis Toolkit: A MapReduce framework for analyzing next-generation DNA sequencing data. *Genome Res.* 20:1297–1303.

Okonechnikov K, Conesa A, García-Alcalde F. 2016. Qualimap 2: Advanced multi-sample quality control for high-throughput sequencing data. *Bioinformatics* 32:292–294.

Powell D, Groβe-Wilde E, Krokene P, Roy A, Chakraborty A, Löfstedt C, Vogel H, Andersson MN, Schlyter F. 2021. A highly-contiguous genome assembly of the Eurasian spruce bark beetle, Ips typographus, provides insight into a major forest pest. *Commun. Biol.* 4:1–9.

Purcell S, Neale B, Todd-Brown K, Thomas L, Ferreira MAR, Bender D, Maller J, Sklar P, De Bakker PIW, Daly MJ, et al. 2007. PLINK: A tool set for whole-genome association and population-based linkage analyses. *Am. J. Hum. Genet.* 81:559–575.

### Supplementary Tables:

**Table S1** Inversion age estimates in *Ips typographus*. The estimates are based on absolute genetic divergence ($d_{xy}$) calculated between two inversion haplotypes and using different mutation rates: 2.10 × $10^{-9}$ (Time 1), 2.90 × $10^{-9}$ (Time 2), 1.17 × $10^{-8}$ (Time 3); time in Myr.

| Inversion | Contig | $d_{xy}$ | Time 1 | Time 2 | Time 2 |
|---|---|---|---|---|---|
| Inv2 | IpsContig2 | 0.0030 | 0.70 | 0.51 | 0.13 |
| Inv3 | IpsContig3 | 0.0063 | 1.49 | 1.08 | 0.27 |
| Inv5 | IpsContig5 | 0.0076 | 1.82 | 1.32 | 0.33 |
| Inv6 | IpsContig6 | 0.0057 | 1.35 | 0.98 | 0.24 |
| Inv7.1 | IpsContig7 | 0.0100 | 2.39 | 1.73 | 0.43 |
| Inv7.2 | IpsContig7 | 0.0065 | 1.55 | 1.12 | 0.28 |
| Inv9 | IpsContig9 | 0.0033 | 0.78 | 0.56 | 0.14 |
| Inv10 | IpsContig10 | 0.0102 | 2.63 | 1.90 | 0.47 |
| Inv12 | IpsContig12 | 0.0090 | 1.55 | 1.12 | 0.28 |
| Inv13 | IpsContig13 | 0.0101 | 2.41 | 1.74 | 0.43 |
| Inv14.1 | IpsContig14 | 0.0109 | 2.59 | 1.87 | 0.46 |
| Inv14.2 | IpsContig14 | 0.0056 | 1.34 | 0.97 | 0.24 |
| Inv14.3 | IpsContig14 | 0.0123 | 2.92 | 2.12 | 0.52 |
| Inv14.4 | IpsContig14 | 0.0125 | 2.59 | 1.88 | 0.47 |
| Inv14.5 | IpsContig14 | 0.0134 | 3.20 | 2.31 | 0.57 |
| Inv14.6 | IpsContig14 | 0.0036 | 0.87 | 0.63 | 0.16 |
| Inv15 | IpsContig15 | 0.0112 | 2.67 | 1.94 | 0.48 |
| Inv16.1 | IpsContig16 | 0.0094 | 2.25 | 1.63 | 0.40 |
| Inv16.2 | IpsContig16 | 0.0041 | 0.97 | 0.70 | 0.17 |
| Inv17 | IpsContig17 | 0.0126 | 3.00 | 2.17 | 0.54 |
| Inv18 | IpsContig18 | 0.0119 | 2.83 | 2.05 | 0.51 |
| Inv22.1 | IpsContig22 | 0.0124 | 2.94 | 2.13 | 0.53 |
| Inv22.2 | IpsContig22 | 0.0123 | 2.94 | 2.13 | 0.53 |
| Inv22.3 | IpsContig22 | 0.0114 | 2.70 | 1.96 | 0.49 |
| Inv22.4 | IpsContig22 | 0.0121 | 2.88 | 2.09 | 0.52 |
| Inv22.5 | IpsContig22 | 0.0035 | 0.82 | 0.60 | 0.15 |
| Inv23.1 | IpsContig23 | 0.0082 | 1.96 | 1.42 | 0.35 |
| Inv23.2 | IpsContig23 | 0.0035 | 0.83 | 0.60 | 0.15 |
| Inv26 | IpsContig26 | 0.0151 | 3.59 | 2.60 | 0.64 |

**Table S2** Genetic differentiation ($F_{ST}$) between investigated *Ips typographus* populations. Population abbreviations are according to Table S5.

| | MEL | SVA | STJ | LAN | EFI | SIL | AAS | ASA | TON | GOS | LUB | BOR | ROZ | STE | BIL | BAW | TRE |
|---|---|---|---|---|---|---|---|---|---|---|---|---|---|---|---|---|---|
| **SVA** | 0.002 | | | | | | | | | | | | | | | | |
| **STJ** | 0.000 | -0.001 | | | | | | | | | | | | | | | |
| **LAN** | -0.001 | 0.001 | 0.000 | | | | | | | | | | | | | | |
| **EFI** | 0.000 | 0.003 | 0.000 | -0.001 | | | | | | | | | | | | | |
| **SIL** | 0.004 | 0.005 | 0.001 | 0.001 | 0.001 | | | | | | | | | | | | |
| **AAS** | 0.003 | 0.007 | 0.002 | 0.001 | 0.003 | 0.000 | | | | | | | | | | | |
| **ASA** | 0.004 | 0.008 | 0.003 | 0.003 | 0.001 | 0.000 | 0.002 | | | | | | | | | | |
| **TON** | 0.001 | 0.003 | -0.001 | 0.001 | 0.001 | 0.001 | -0.001 | 0.002 | | | | | | | | | |
| **GOS** | 0.006 | 0.010 | 0.007 | 0.005 | 0.003 | 0.005 | 0.007 | 0.005 | 0.005 | | | | | | | | |
| **LUB** | 0.006 | 0.010 | 0.006 | 0.004 | 0.003 | 0.003 | 0.004 | 0.005 | 0.004 | -0.001 | | | | | | | |
| **BOR** | 0.003 | 0.014 | 0.008 | 0.005 | 0.005 | 0.003 | 0.005 | 0.004 | 0.005 | 0.004 | 0.003 | | | | | | |
| **ROZ** | 0.014 | 0.020 | 0.015 | 0.014 | 0.014 | 0.011 | 0.012 | 0.012 | 0.012 | 0.002 | 0.003 | 0.007 | | | | | |
| **STE** | 0.017 | 0.025 | 0.021 | 0.020 | 0.020 | 0.023 | 0.022 | 0.022 | 0.018 | 0.009 | 0.014 | 0.015 | 0.008 | | | | |
| **BIL** | 0.029 | 0.035 | 0.029 | 0.026 | 0.026 | 0.021 | 0.021 | 0.025 | 0.024 | 0.009 | 0.011 | 0.018 | 0.003 | 0.011 | | | |
| **BAW** | 0.018 | 0.026 | 0.024 | 0.021 | 0.023 | 0.023 | 0.025 | 0.024 | 0.023 | 0.010 | 0.014 | 0.013 | 0.007 | 0.000 | 0.011 | | |
| **TRE** | 0.024 | 0.034 | 0.027 | 0.024 | 0.023 | 0.021 | 0.021 | 0.023 | 0.022 | 0.007 | 0.013 | 0.014 | 0.005 | 0.006 | 0.001 | 0.006 | |
| **LIN** | 0.019 | 0.027 | 0.023 | 0.023 | 0.023 | 0.025 | 0.024 | 0.022 | 0.021 | 0.008 | 0.014 | 0.015 | 0.006 | 0.000 | 0.007 | 0.000 | 0.004 |

**Table S3** Association between inversion haplotype and diapause phenotype in *Ips typographus*. Results of the genic differentiation test (exact G test). The test was done using individuals of known diapause phenotype. Only inversions that were genotyped for the whole species range were tested.

| Inversion | Contig | p value |
|---|---|---|
| Inv3 | IpsContig3 | 1.00 |
| Inv6 | IpsContig6 | not polymorphic |
| Inv7.1 | IpsContig7 | 0.74 |
| Inv7.2 | IpsContig7 | 1.00 |
| Inv9 | IpsContig9 | 0.44 |
| Inv10 | IpsContig10 | 0.50 |
| Inv12 | IpsContig12 | 0.15 |
| Inv13 | IpsContig13 | 1.00 |
| Inv14.1 | IpsContig14 | 0.75 |
| Inv14.2 | IpsContig14 | 0.57 |
| Inv14.3 | IpsContig14 | 0.30 |
| Inv14.4 | IpsContig14 | 0.48 |
| Inv14.5 | IpsContig14 | 1.00 |
| Inv14.6 | IpsContig14 | not polymorphic |
| Inv15 | IpsContig15 | 1.00 |
| Inv16.1 | IpsContig16 | 0.42 |
| Inv17 | IpsContig17 | 0.31 |
| Inv18 | IpsContig18 | 0.73 |
| Inv22.1 | IpsContig22 | 0.49 |
| Inv22.2 | IpsContig22 | 0.20 |
| Inv22.3 | IpsContig22 | 0.64 |
| Inv22.4 | IpsContig22 | 0.31 |
| Inv22.5 | IpsContig22 | not polymorphic |
| Inv23.1 | IpsContig23 | 0.43 |
| Inv26 | IpsContig26 | 0.32 |

**Table S4** Number of nonsynonymous and synonymous SNPs polymorphic in the spruce bark beetle populations. #NS – number of nonsynonymous SNPs; #SS – number of synonymous SNPs. Table sorted by #NS.

| OR ID | Contig ID | Inversion ID | #NS | #SS |
|---|---|---|---|---|
| ItypOR22CTE | IpsContig22 | Inv22.3 | 54 | 32 |
| ItypOR43JOI | IpsContig5 | Inv5 | 50 | 39 |
| ItypOR53 | IpsContig5 | Inv5 | 44 | 40 |
| ItypOR3 | IpsContig5 | Inv5 | 43 | 33 |
| ItypOR2 | IpsContig5 | Inv5 | 42 | 35 |
| ItypOR49 | IpsContig13 | Inv13 | 41 | 40 |
| ItypOR19 | IpsContig5 | Inv5 | 38 | 30 |
| ItypOR30 | IpsContig16 | Inv16.1, Inv16.2 | 38 | 14 |
| ItypOR4 | IpsContig5 | Inv5 | 38 | 37 |
| ItypOR36 | IpsContig14 | Inv14, Inv14.6 | 37 | 44 |
| ItypOR50 | IpsContig5 | Inv5 | 36 | 25 |
| ItypOR20NTE | IpsContig14 | Inv14, Inv14.6 | 35 | 37 |
| ItypOR1 | IpsContig7 | Inv7.2 | 31 | 22 |
| ItypOR17 | IpsContig7 | Inv7.2 | 31 | 37 |
| ItypOR33 | IpsContig5 | Inv5 | 31 | 26 |
| ItypOR40 | IpsContig5 | Inv5 | 29 | 16 |
| ItypOR23 | IpsContig13 | Inv13 | 28 | 24 |
| ItypOR28 | IpsContig13 | Inv13 | 28 | 37 |
| ItypOR31 | IpsContig16 | Inv16.1, Inv16.2 | 22 | 38 |
| ItypOR35JF | IpsContig16 | Inv16.1, Inv16.2 | 22 | 12 |
| ItypOR10 | IpsContig5 | Inv5 | 20 | 14 |
| ItypOR27 | IpsContig13 | Inv13 | 19 | 37 |
| ItypOR41 | IpsContig5 | Inv5 | 17 | 14 |
| ItypOR29 | IpsContig5 | Inv5 | 15 | 10 |
| ItypOR16 | IpsContig16 | Inv16.1, Inv16.2 | 12 | 13 |
| ItypOR11 | IpsContig16 | Inv16.1, Inv16.2 | 11 | 2 |

| | | | | |
|---|---|---|---|---|
| ItypOR34 | IpsContig5 | Inv5 | 10 | 12 |
| ItypOR52NTE | IpsContig5 | Inv5 | 7 | 6 |
| ItypOR18JF | IpsContig14 | Inv14, Inv14.6 | 0 | 0 |
| ItypOR44 | IpsContig14 | Inv14, Inv14.6 | 0 | 0 |
| ItypOR47 | IpsContig5 | Inv5 | 0 | 0 |
| ItypOR58 | IpsContig16 | Inv16.1, Inv16.2 | 0 | 0 |
| ItypOR58 | IpsContig23 | Inv23.1, Inv23.2 | 0 | 0 |
| ItypOR9 | IpsContig16 | Inv16.1, Inv16.2 | 0 | 0 |
| ItypOR9 | IpsContig23 | Inv23.1, Inv23.2 | 0 | 0 |

**Table S5** Nonsynonymous divergence between inversion haplotypes within OR coding sequence. Numbers indicate number of nonsynonymous SNPs with different $d_{xy}$ thresholds. Table sorted by number of SNPs with $d_{xy}$ >0.9.

| OR ID | Contig ID | Inversion ID | $d_{xy}$ >0.9 | $d_{xy}$ >0.8 | $d_{xy}$ >0.5 | $d_{xy}$ <0.1 |
|---|---|---|---|---|---|---|
| ItypOR43JOI | IpsContig5 | Inv5 | 7 | 7 | 7 | 41 |
| ItypOR30 | IpsContig16 | Inv16.1 | 4 | 4 | 4 | 30 |
| ItypOR20NTE | IpsContig14 | Inv14.1 | 3 | 3 | 5 | 25 |
| ItypOR36 | IpsContig14 | Inv14.1 | 3 | 4 | 4 | 31 |
| ItypOR23 | IpsContig13 | Inv13 | 2 | 2 | 5 | 18 |
| ItypOR31 | IpsContig16 | Inv16.1 | 2 | 4 | 4 | 17 |
| ItypOR16 | IpsContig16 | Inv16.1 | 1 | 1 | 1 | 11 |
| ItypOR27 | IpsContig13 | Inv13 | 1 | 2 | 3 | 11 |
| ItypOR29 | IpsContig5 | Inv5 | 1 | 1 | 1 | 12 |
| ItypOR1 | IpsContig7 | Inv7.2 | 0 | 0 | 1 | 26 |
| ItypOR10 | IpsContig5 | Inv5 | 0 | 0 | 1 | 17 |
| ItypOR11 | IpsContig16 | Inv16.2 | 0 | 0 | 0 | 9 |
| ItypOR11 | IpsContig16 | Inv16.1 | 0 | 0 | 2 | 9 |
| ItypOR16 | IpsContig16 | Inv16.2 | 0 | 0 | 1 | 11 |
| ItypOR17 | IpsContig7 | Inv7.2 | 0 | 0 | 0 | 21 |
| ItypOR19 | IpsContig5 | Inv5 | 0 | 0 | 1 | 33 |
| ItypOR2 | IpsContig5 | Inv5 | 0 | 1 | 1 | 35 |
| ItypOR20NTE | IpsContig14 | Inv14.6 | 0 | 0 | 1 | 27 |
| ItypOR22CTE | IpsContig22 | Inv22.3 | 0 | 4 | 14 | 23 |
| ItypOR28 | IpsContig13 | Inv13 | 0 | 0 | 2 | 24 |
| ItypOR3 | IpsContig5 | Inv5 | 0 | 1 | 11 | 32 |
| ItypOR30 | IpsContig16 | Inv16.2 | 0 | 0 | 3 | 32 |
| ItypOR31 | IpsContig16 | Inv16.2 | 0 | 0 | 2 | 17 |
| ItypOR33 | IpsContig5 | Inv5 | 0 | 0 | 1 | 28 |
| ItypOR34 | IpsContig5 | Inv5 | 0 | 0 | 0 | 8 |
| ItypOR35JF | IpsContig16 | Inv16.2 | 0 | 0 | 0 | 22 |

| | | | | | | |
|---|---|---|---|---|---|---|
| ItypOR35JF | IpsContig16 | Inv16 | 0 | 0 | 0 | 19 |
| ItypOR36 | IpsContig14 | Inv14.6 | 0 | 0 | 1 | 34 |
| ItypOR4 | IpsContig5 | Inv5 | 0 | 0 | 0 | 35 |
| ItypOR40 | IpsContig5 | Inv5 | 0 | 1 | 3 | 24 |
| ItypOR41 | IpsContig5 | Inv5 | 0 | 1 | 1 | 16 |
| ItypOR49 | IpsContig13 | Inv13 | 0 | 0 | 0 | 39 |
| ItypOR50 | IpsContig5 | Inv5 | 0 | 1 | 3 | 31 |
| ItypOR52NTE | IpsContig5 | Inv5 | 0 | 0 | 0 | 7 |
| ItypOR53 | IpsContig5 | Inv5 | 0 | 0 | 2 | 34 |

**Table S6** Gene Ontology categories significantly enriched in inversions that correlate with latitude. MF – Molecular function; BP – Biological process; CC – cellular component. Categories associated with olfaction are bolded.

| GO ID | Term | p-value | type |
|---|---|---|---|
| GO:0000287 | magnesium ion binding | 0.00048 | MF |
| GO:0006338 | chromatin remodeling | 0.00090 | BP |
| GO:0019438 | aromatic compound biosynthetic process | 0.00130 | BP |
| GO:1901362 | organic cyclic compound biosynthetic process | 0.00140 | BP |
| GO:0070603 | SWI/SNF superfamily-type complex | 0.00160 | CC |
| GO:0007264 | small GTPase mediated signal transduction | 0.00340 | BP |
| GO:0140657 | ATP-dependent activity | 0.00415 | MF |
| GO:0018130 | heterocycle biosynthetic process | 0.00560 | BP |
| GO:0050794 | regulation of cellular process | 0.00690 | BP |
| GO:0006886 | intracellular protein transport | 0.00700 | BP |
| GO:0140096 | catalytic activity. acting on a protein | 0.00921 | MF |
| GO:0043547 | positive regulation of GTPase activity | 0.00930 | BP |
| GO:0015914 | phospholipid transport | 0.00930 | BP |
| GO:0005096 | GTPase activator activity | 0.00941 | MF |
| GO:0030154 | cell differentiation | 0.01350 | BP |
| GO:0015078 | proton transmembrane transporter activity | 0.01435 | MF |
| GO:1902600 | proton transmembrane transport | 0.01490 | BP |
| GO:0045454 | cell redox homeostasis | 0.01490 | BP |
| GO:0008234 | cysteine-type peptidase activity | 0.01664 | MF |
| GO:0042254 | ribosome biogenesis | 0.01890 | BP |
| GO:0005319 | lipid transporter activity | 0.01912 | MF |
| GO:0048856 | anatomical structure development | 0.01990 | BP |
| **GO:0005549** | **odorant binding** | **0.02184** | **MF** |
| GO:0016853 | isomerase activity | 0.02201 | MF |
| GO:0022853 | active monoatomic ion transmembrane transporter activity | 0.02269 | MF |
| GO:0010629 | negative regulation of gene expression | 0.02340 | BP |
| GO:0055080 | monoatomic cation homeostasis | 0.02540 | BP |
| GO:0000413 | protein peptidyl-prolyl isomerization | 0.02540 | BP |
| GO:0003755 | peptidyl-prolyl cis-trans isomerase activity | 0.02566 | MF |
| GO:0008324 | monoatomic cation transmembrane transporter activity | 0.02929 | MF |
| GO:0006366 | transcription by RNA polymerase II | 0.02940 | BP |
| GO:0005524 | ATP binding | 0.03101 | MF |
| GO:0015031 | protein transport | 0.03220 | BP |
| GO:0006289 | nucleotide-excision repair | 0.03300 | BP |
| GO:0015291 | secondary active transmembrane transporter activity | 0.03339 | MF |
| GO:0098655 | monoatomic cation transmembrane transport | 0.03570 | BP |

| | | | |
|---|---|---|---|
| GO:0042302 | structural constituent of cuticle | 0.03589 | MF |
| GO:0065007 | biological regulation | 0.03780 | BP |
| **GO:0050911** | **detection of chemical stimulus involved in sensory perception of smell** | **0.04190** | **BP** |
| GO:0009966 | regulation of signal transduction | 0.04200 | BP |
| **GO:0004984** | **olfactory receptor activity** | **0.04232** | **MF** |
| GO:0044270 | cellular nitrogen compound catabolic process | 0.04340 | BP |
| GO:0046700 | heterocycle catabolic process | 0.04340 | BP |
| GO:0006401 | RNA catabolic process | 0.04410 | BP |
| GO:0045859 | regulation of protein kinase activity | 0.04410 | BP |
| GO:0032446 | protein modification by small protein conjugation | 0.04460 | BP |
| GO:0034220 | monoatomic ion transmembrane transport | 0.04660 | BP |
| GO:0043168 | anion binding | 0.04756 | MF |
| GO:0016773 | phosphotransferase activity. alcohol group as acceptor | 0.04925 | MF |

**Table S7** Gene Ontology categories significantly enriched in inversions. MF – Molecular function; BP – Biological process; CC – cellular component. Categories associated with olfaction indicated are bolded.

| GO ID | Term | p-value | type |
|---|---|---|---|
| GO:0005975 | carbohydrate metabolic process | 0.0000061 | BP |
| GO:0016491 | oxidoreductase activity | 0.0000091 | MF |
| GO:0071555 | cell wall organization | 0.000038 | BP |
| GO:0016853 | isomerase activity | 0.000065 | MF |
| GO:0030286 | dynein complex | 0.00022 | CC |
| GO:0005509 | calcium ion binding | 0.00082 | MF |
| GO:0006886 | intracellular protein transport | 0.0015 | BP |
| GO:0022857 | transmembrane transporter activity | 0.00158 | MF |
| GO:0005524 | ATP binding | 0.00225 | MF |
| GO:0006355 | regulation of DNA-templated transcription | 0.0027 | BP |
| GO:0000287 | magnesium ion binding | 0.0039 | MF |
| GO:0020037 | heme binding | 0.00405 | MF |
| GO:0043547 | positive regulation of GTPase activity | 0.0042 | BP |
| GO:0005096 | GTPase activator activity | 0.00435 | MF |
| GO:0015631 | tubulin binding | 0.00454 | MF |
| GO:0016747 | acyltransferase activity. transferring groups other than amino-acyl groups | 0.00522 | MF |
| GO:0006457 | protein folding | 0.0061 | BP |
| GO:0004553 | hydrolase activity. hydrolyzing O-glycosyl compounds | 0.00618 | MF |
| GO:0016705 | oxidoreductase activity. acting on paired donors. with incorporation or reduction of molecular oxygen | 0.00713 | MF |
| GO:0005506 | iron ion binding | 0.00714 | MF |
| GO:0003777 | microtubule motor activity | 0.00922 | MF |
| GO:0016810 | hydrolase activity. acting on carbon-nitrogen (but not peptide) bonds | 0.00929 | MF |
| GO:0006811 | monoatomic ion transport | 0.0101 | BP |
| GO:0003723 | RNA binding | 0.01011 | MF |
| GO:0016773 | phosphotransferase activity. alcohol group as acceptor | 0.01052 | MF |
| GO:0008234 | cysteine-type peptidase activity | 0.0111 | MF |
| GO:0055085 | transmembrane transport | 0.0125 | BP |
| GO:0035556 | intracellular signal transduction | 0.0148 | BP |
| GO:0019438 | aromatic compound biosynthetic process | 0.0165 | BP |
| GO:0005319 | lipid transporter activity | 0.01735 | MF |
| GO:0016667 | oxidoreductase activity. acting on a sulfur group of donors | 0.01735 | MF |
| GO:0006468 | protein phosphorylation | 0.0176 | BP |
| GO:0004812 | aminoacyl-tRNA ligase activity | 0.01774 | MF |

| | | | |
|---|---|---|---|
| GO:0005829 | cytosol | 0.01818 | CC |
| GO:1901576 | organic substance biosynthetic process | 0.0192 | BP |
| GO:0005783 | endoplasmic reticulum | 0.01972 | CC |
| GO:0050896 | response to stimulus | 0.0205 | BP |
| GO:0016579 | protein deubiquitination | 0.0206 | BP |
| **GO:0050911** | **detection of chemical stimulus involved in sensory perception of smell** | **0.0206** | **BP** |
| **GO:0004984** | **olfactory receptor activity** | **0.02136** | **MF** |
| GO:0004497 | monooxygenase activity | 0.02147 | MF |
| GO:0015914 | phospholipid transport | 0.0228 | BP |
| GO:0007155 | cell adhesion | 0.0228 | BP |
| GO:0140096 | catalytic activity. acting on a protein | 0.02711 | MF |
| GO:0016798 | hydrolase activity. acting on glycosyl bonds | 0.02754 | MF |
| GO:0015078 | proton transmembrane transporter activity | 0.02821 | MF |
| GO:0043565 | sequence-specific DNA binding | 0.02949 | MF |
| GO:0005794 | Golgi apparatus | 0.02985 | CC |
| GO:0030054 | cell junction | 0.03063 | CC |
| GO:0003700 | DNA-binding transcription factor activity | 0.03085 | MF |
| GO:0018130 | heterocycle biosynthetic process | 0.031 | BP |
| GO:0044249 | cellular biosynthetic process | 0.0327 | BP |
| GO:0006418 | tRNA aminoacylation for protein translation | 0.0344 | BP |
| GO:1901362 | organic cyclic compound biosynthetic process | 0.0361 | BP |
| GO:0005737 | cytoplasm | 0.03646 | CC |
| GO:0005886 | plasma membrane | 0.03892 | CC |
| GO:0046496 | nicotinamide nucleotide metabolic process | 0.0394 | BP |
| GO:0044272 | sulfur compound biosynthetic process | 0.0394 | BP |
| GO:0003690 | double-stranded DNA binding | 0.04048 | MF |
| GO:0050790 | regulation of catalytic activity | 0.0431 | BP |
| GO:0000398 | mRNA splicing. via spliceosome | 0.044 | BP |
| GO:0018193 | peptidyl-amino acid modification | 0.0442 | BP |
| GO:0008810 | cellulase activity | 0.04579 | MF |
| GO:0015291 | secondary active transmembrane transporter activity | 0.04579 | MF |
| GO:0006396 | RNA processing | 0.0469 | BP |
| GO:0140513 | nuclear protein-containing complex | 0.04851 | CC |
| GO:1902600 | proton transmembrane transport | 0.049 | BP |
| GO:0007264 | small GTPase-mediated signal transduction | 0.049 | BP |

**Table S8** Genes closest to SNPs significant in the genotype-environment association analysis.

| Gene | Description | GO ID |
|---|---|---|
| Ityp00693 | roundabout homolog 2-like | NA |
| Ityp07355 | PREDICTED: mitoferrin-1-like | GO:0016021 |
| Ityp07620 | U2 small nuclear ribonucleoprotein auxiliary factor 35 kDa subunit-related protein 1 | NA |
| Ityp08738 | poly [ADP-ribose] polymerase 12-like | NA |
| Ityp08741 | general transcription factor IIE subunit 2-like | GO:0003677.GO:0005673.GO:0006367 |
| Ityp09110 | inotocin receptor | GO:0005000.GO:0005887.GO:0007186.GO:0032870.GO:0042277 |
| Ityp14927 | copper-transporting ATPase 1 isoform X1 | NA |
| Ityp17124 | PREDICTED: uncharacterized protein LOC109541129 | NA |
| Ityp17174 | chondroitin sulfate proteoglycan 4 isoform X2 | GO:0016491.GO:0055114 |
| Ityp17175 | eukaryotic translation initiation factor 2-alpha kinase | NA |
| Ityp17177 | PREDICTED: uncharacterized protein LOC109540861 isoform X1 | NA |
| Ityp17181 | RNA-binding protein 24-A-like | GO:0003729 |
| Ityp17183 | solute carrier organic anion transporter family member 2A1-like | NA |
| Ityp17184 | PREDICTED: uncharacterized protein LOC109541131 | NA |

**Table S9.** Significant deviations from Hardy-Weinberg expectations for inversion genotypes at whole species range and within genetic groups (southern, northern and Polish)**.** Significance level at 0.05 after Bonferroni correction.

| Inversion | Data | Observed number of genotypes | | | | | | Expected number of genotypes | | | | | | P-value | Alfa at 0.05 |
|---|---|---|---|---|---|---|---|---|---|---|---|---|---|---|---|
| | | AA | AB | AC | BB | BC | CC | AA | AB | AC | BB | BC | CC | | |
| Inv12 | whole species range | 18 | 55 | | 167 | | | 9 | 74 | | 158 | | | 0.00022 | 0.001724 |
| Inv22.4 | | 46 | 101 | 88 | 3 | 2 | 0 | 82 | 64 | 53 | 12 | 20 | 8 | 0.00000 | 0.001724 |
| Inv22.4 | northern | 28 | 45 | 47 | 0 | 0 | 0 | 46 | 28 | 29 | 4 | 9 | 5 | 0.00000 | 0.000617 |
| Inv22.4 | polish | 8 | 20 | 24 | 2 | 0 | 0 | 17 | 13 | 13 | 3 | 5 | 3 | 0.00000 | 0.000617 |
| Inv22.4 | southern | 10 | 36 | 17 | 1 | 2 | 0 | 20 | 22 | 11 | 6 | 6 | 1 | 0.00000 | 0.000617 |

**Table S10** Mutation load analyses in *Ips typographus*.

The ratio of nonsynonymous to synonymous nucleotide diversity ($\pi_N/\pi_S$) was calculated in 200 kb windows along the genome. The windows were separated into windows inside or outside inversions. Windows of the collinear part of the genome were analyzed including (no strict filtering) or excluding windows that come from the contigs harboring inversions (strict filtering). Mutation load for inversions was calculated for all individuals (all), individuals that were homozygous for less common inversion haplotypes (minor), and individuals that were homozygous for more common inversion haplotypes (major). Different filtering criteria were tested. including minimum number of genes (gene), minimum number of individuals carrying a given haplotype (ind), and minimum number of windows (win). For further details see the main text.

| **Strict filtering for collinear genome** | **Inversion groups** | **Filtering criteria gene/ind/win** | **Median $\pi_N/\pi_S$** | | **P-value** |
|---|---|---|---|---|---|
| | | | **for collinear genome** | **for inversion group** | |
| no | all | 5/4/4 | 0.1311 | 0.1161 | 0.001 |
| | minor | | 0.1311 | 0.1224 | 0.049 |
| | major | | 0.1311 | 0.1131 | <0.001 |
| | all | 10/10/4 | 0.1293 | 0.1145 | 0.006 |
| | minor | | 0.1293 | 0.1161 | 0.537 |
| | major | | 0.1293 | 0.1129 | <0.001 |
| yes | all | 5/4/4 | 0.1285 | 0.1161 | 0.047 |
| | minor | | 0.1285 | 0.1224 | 0.216 |
| | major | | 0.1285 | 0.1131 | 0.016 |
| | all | 10/10/4 | 0.1284 | 0.1145 | 0.062 |
| | minor | | 0.1284 | 0.1161 | 0.663 |
| | major | | 0.1284 | 0.1129 | <0.001 |

**Table S11** Details about sites where *Ips typographus* were collected. ‘Group’ denotes in what part of the species’ distribution area beetles were collected: northern (N), southern (S) or Poland (P). ‘N’ gives the number of individuals collected per site.

| Population ID | Population name | Latitude | Longitude | Country | Group | N |
|---|---|---|---|---|---|---|
| AAS | Ås | 59.667 | 10.793 | Norway | N | 13 |
| ASA | Asa | 57.165 | 14.783 | Sweden | N | 14 |
| BAW | Bavarian Forest | 48.960 | 13.395 | Germany | S | 11 |
| BIL | Bílkovice | 49.761 | 14.848 | Czechia | S | 14 |
| BOR | Borki | 54.090 | 21.912 | Poland | P | 12 |
| EFI | Eastern Finland | 62.492 | 30.010 | Finland | N | 13 |
| GOS | Gościnno | 54.047 | 15.657 | Poland | P | 14 |
| LAN | Länsi | 61.723 | 23.633 | Finland | N | 13 |
| LIN | Linz | 48.092 | 13.874 | Austria | S | 14 |
| LUB | Lubaszki | 54.057 | 17.556 | Poland | P | 14 |
| MEL | Mellakoski | 66.399 | 24.440 | Finland | N | 14 |
| ROZ | Roztocze | 50.508 | 22.786 | Poland | P | 14 |
| SIL | Siljanfors | 60.757 | 14.066 | Sweden | N | 14 |
| STE | Steigerwald | 49.622 | 10.263 | Germany | S | 14 |
| STJ | Stjørdal | 63.469 | 10.918 | Norway | N | 13 |
| SVA | Svartberget | 64.236 | 19.570 | Sweden | N | 13 |
| TON | Tönnersjö | 56.643 | 13.070 | Sweden | N | 13 |
| TRE | Třebíč | 49.212 | 15.879 | Czechia | S | 13 |

**Supplementary Figures**

Figure S1 Comparison of *Ips typographus* and its close relative - *I. nitidus* – genomes, based on alignments of two assemblies. For clarity, only the 36 longest contigs of focal species are contrasted with 16 *I. nitidus* chromosomes in the dotplot (D-GENIES version 1.5.0; (Cabanettes and Klopp 2018), denoted as T1 – T36 and N1 – N16, respectively. (A) Alignment with *I. nitidius* as target vs *I. typographus* as mapping query. (B) Alignment with *I. typographus* as target vs *I. nitidus* as mapping query.

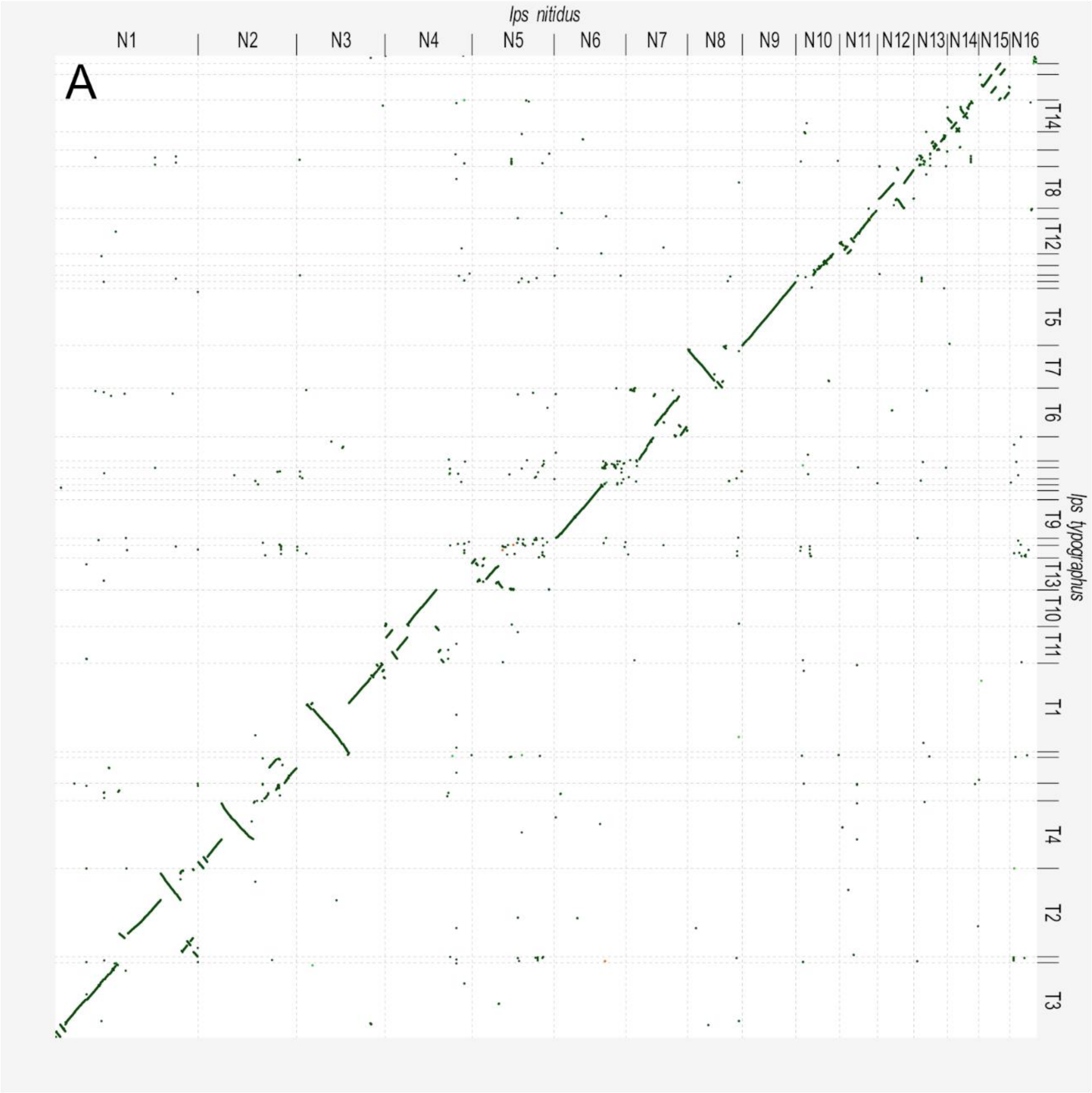

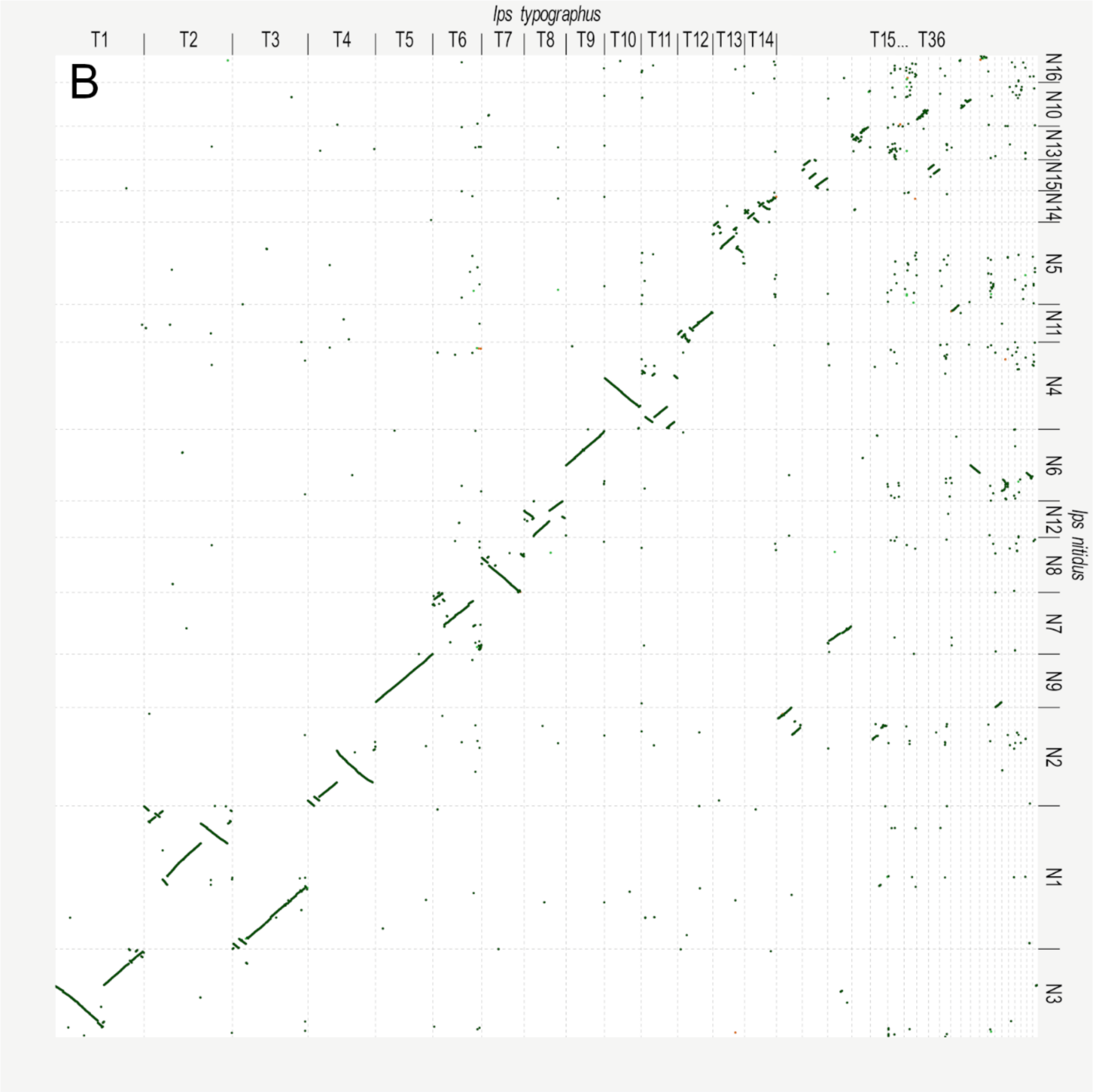
B
Ips typographus
T1
T2
T3
T4
T5
T6
T7
T8
T9
T10
T11
T12
T13
T14
T15... T36
Ips nitidus
N16
N10
N13
N15
N14
N5
N11
N4
N6
N12
N8
N7
N9
N2
N1
N3

**Figure S2** Inversion patterns in the *Ips typographus* genome. Each subplot shows multiple lines of evidence for inversion polymorphism: localPCA (A), per contig PCA (B), PCA for putative inversion region only (color coded by population (C) and heterozygosity (D)), $F_{ST}$ between inversion haplotypes (E), LD patterns for homozygous (upper triangle) and all individuals (bottom triangle) (F); population recombination rate for three inversion genotypes (G). (A) Identification of inversion coordinates with MDS, where MDS coordinates are plotted against the length of the contig in Mb. Y-axis title shows on which dimension a given inversion was identified as an outlier.  Each dot represents a single window, red dots - outlier inversion windows, grey dots - rest of the contig. (B) PCA plot for the whole contig; each dot is an individual colored according to a population it belongs to, blue colors represent populations from northern group, yellow colors – populations from Polish group and red and green colors – populations from southern group. (C) PCA plot for the inverted region each dot is an individual colored the same way as on the panel B. (D) PCA plot for the inverted region, each dot is colored according to the level of the observed heterozygosity in the inverted region.  (E) Genetic differentiation ($F_{\mathrm{ST}}$) between inversion haplotypes (major – MJ and minor MN inversion haplotype, and recombinant haplotype R, if present) along the contig. (F) Contig LD patterns for homozygous (upper triangle) and all individuals (bottom triangle). (G) Population recombination rate (ρ) calculated along the contig between inversion genotypes: M J – homozygotes of the more frequent haplotype. MN – homozygotes of the less frequent haplotype and HET - heterozygotes.

**Inv 2**

Inv2 could be genotyped only in southern group and C and D PCAs are shown only for southern populations for clearer visualization.

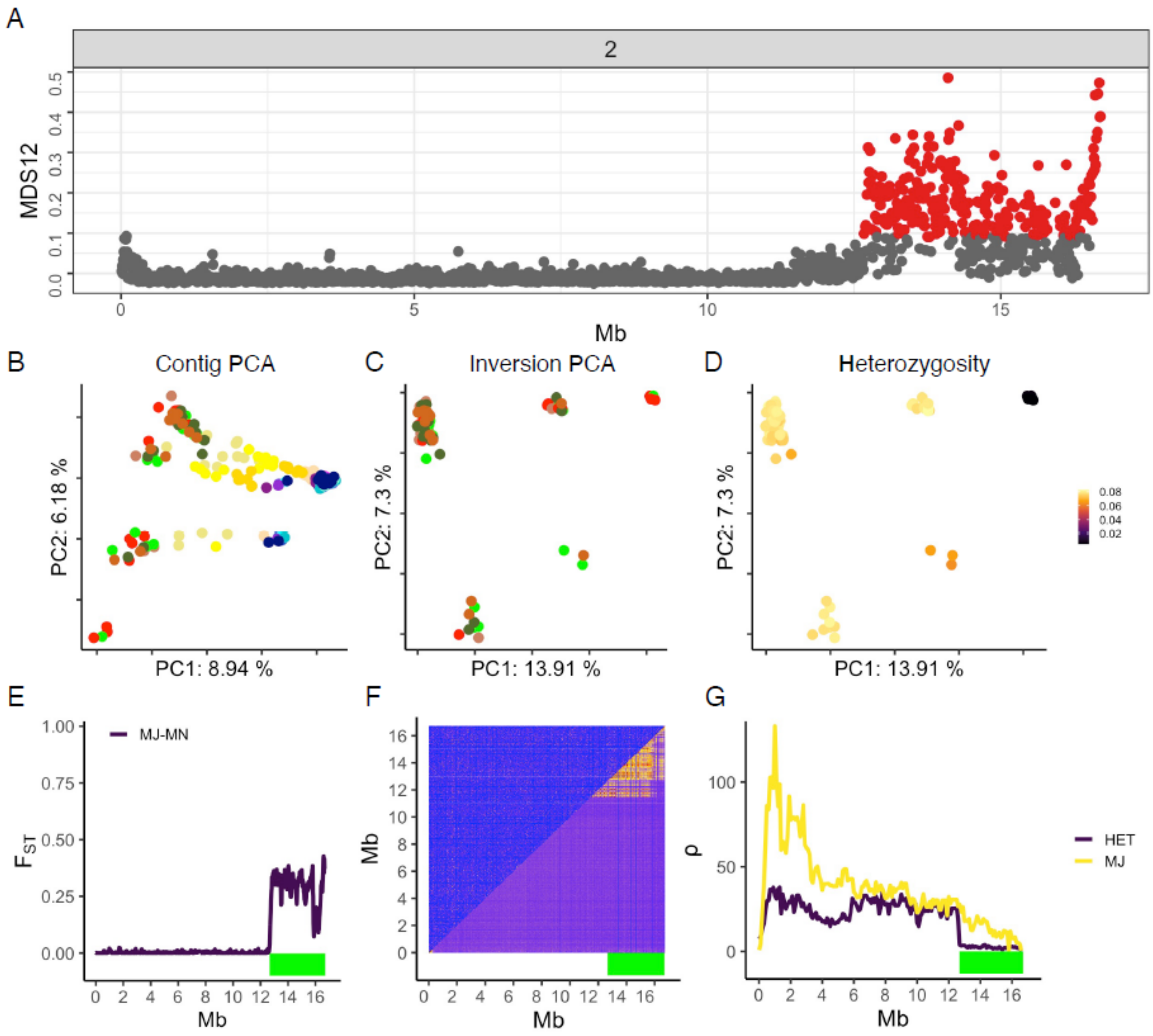

**Inv3**

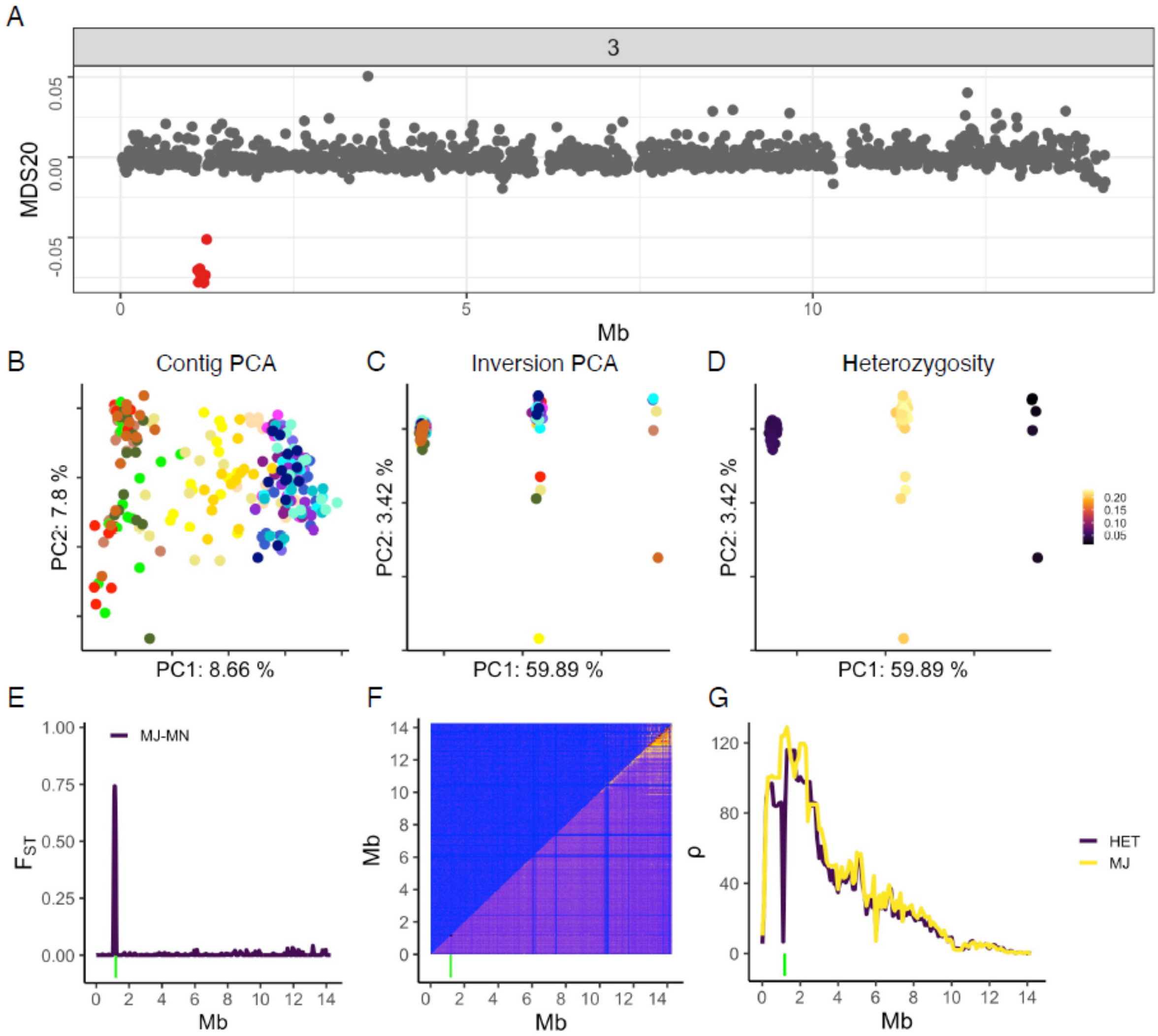
A
3
MDS20
0.05
0.00
-0.05
0
5
10
Mb
B
Contig PCA
PC2: 7.8 %
PC1: 8.66 %
C
Inversion PCA
PC2: 3.42 %
PC1: 59.89 %
D
Heterozygosity
PC2: 3.42 %
PC1: 59.89 %
0.20
0.15
0.10
0.05
E
MJ-MN
$F_{ST}$
1.00
0.75
0.50
0.25
0.00
0 2 4 6 8 10 12 14
Mb
F
Mb
14 12 10 8 6 4 2 0
0 2 4 6 8 10 12 14
Mb
G
ρ
120
80
40
0
0 2 4 6 8 10 12 14
Mb
HET
MJ

**Inv5**

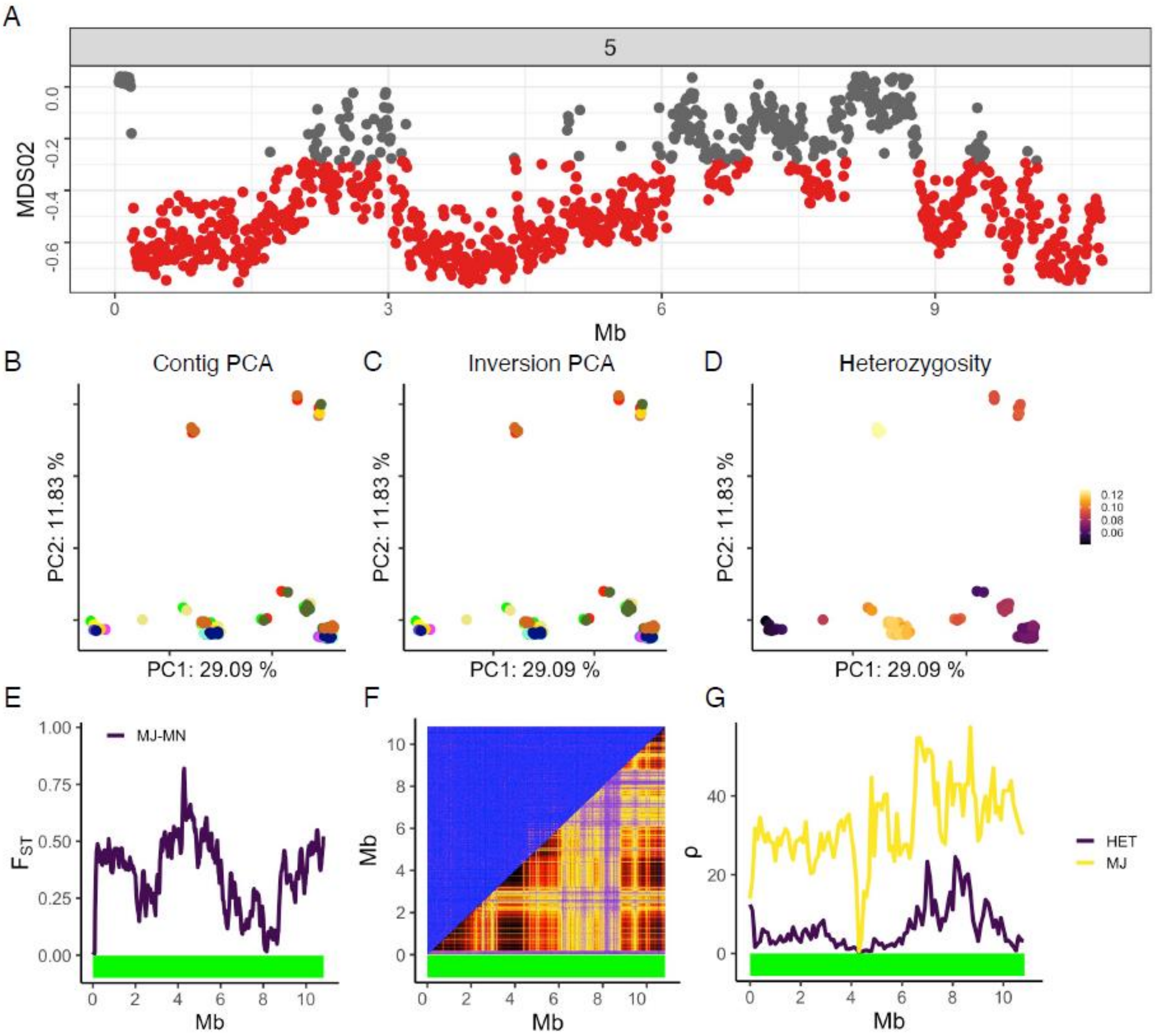

A
5
MDS02
0.0
-0.2
-0.4
-0.6
0
3
6
9
Mb
B
Contig PCA
PC2: 11.83 %
PC1: 29.09 %
C
Inversion PCA
PC2: 11.83 %
PC1: 29.09 %
D
Heterozygosity
PC2: 11.83 %
PC1: 29.09 %
0.12
0.10
0.08
0.06
E
MJ-MN
1.00
0.75
0.50
0.25
0.00
$F_{ST}$
0
2
4
6
8
10
Mb
F
10
8
6
4
2
0
Mb
0
2
4
6
8
10
Mb
G
40
20
0
ρ
HET
MJ
0
2
4
6
8
10
Mb

**Inv6**

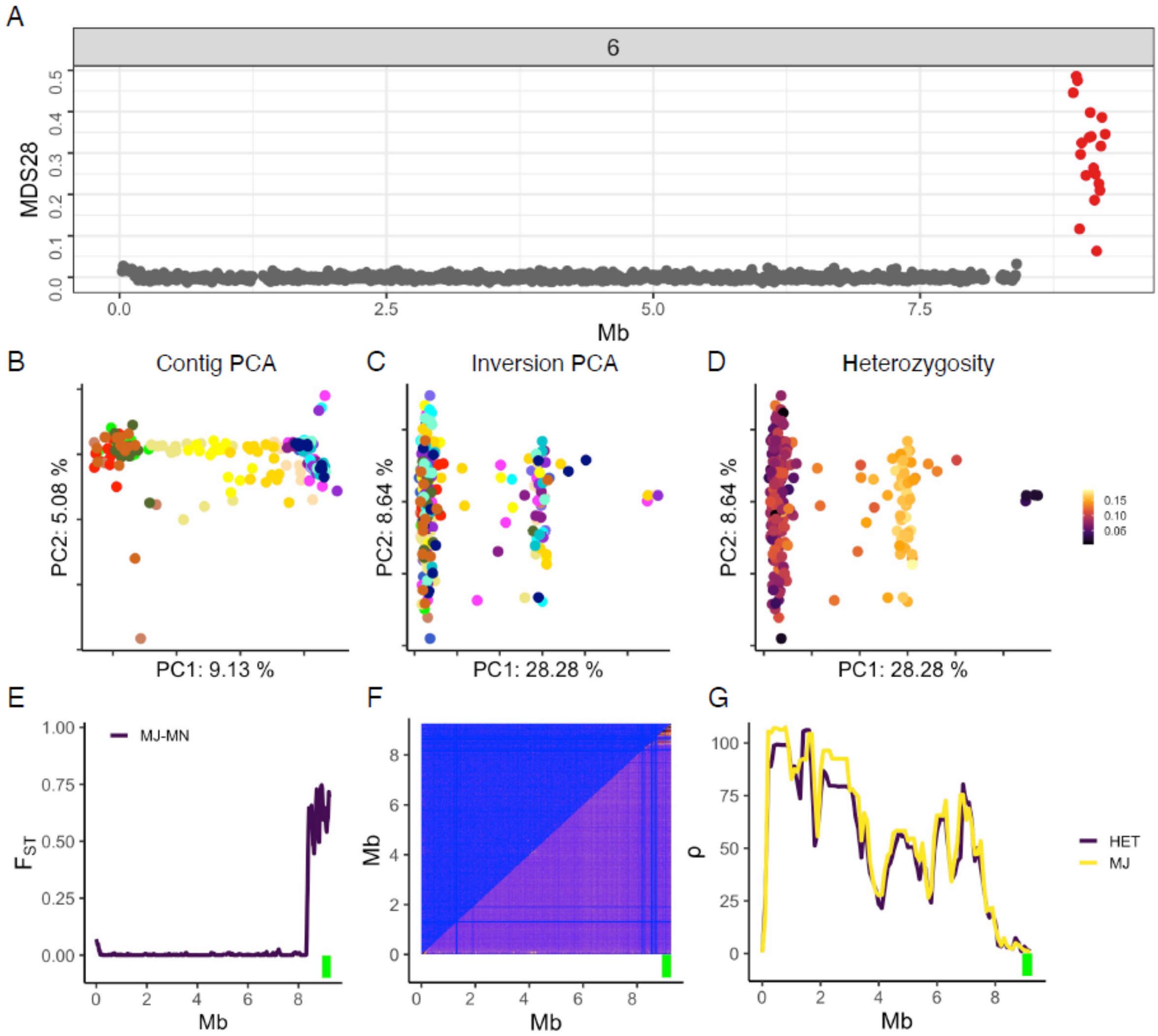

A
6
MDS28
0.0
0.1
0.2
0.3
0.4
0.5
0.0
2.5
5.0
7.5
Mb
B
Contig PCA
PC2: 5.08 %
PC1: 9.13 %
C
Inversion PCA
PC2: 8.64 %
PC1: 28.28 %
D
Heterozygosity
PC2: 8.64 %
PC1: 28.28 %
0.15
0.10
0.05
E
MJ-MN
$F_{ST}$
1.00
0.75
0.50
0.25
0.00
0
2
4
6
8
Mb
F
Mb
8
6
4
2
0
0
2
4
6
8
Mb
G
ρ
100
75
50
25
0
0
2
4
6
8
Mb
HET
MJ

**Inv7.1**

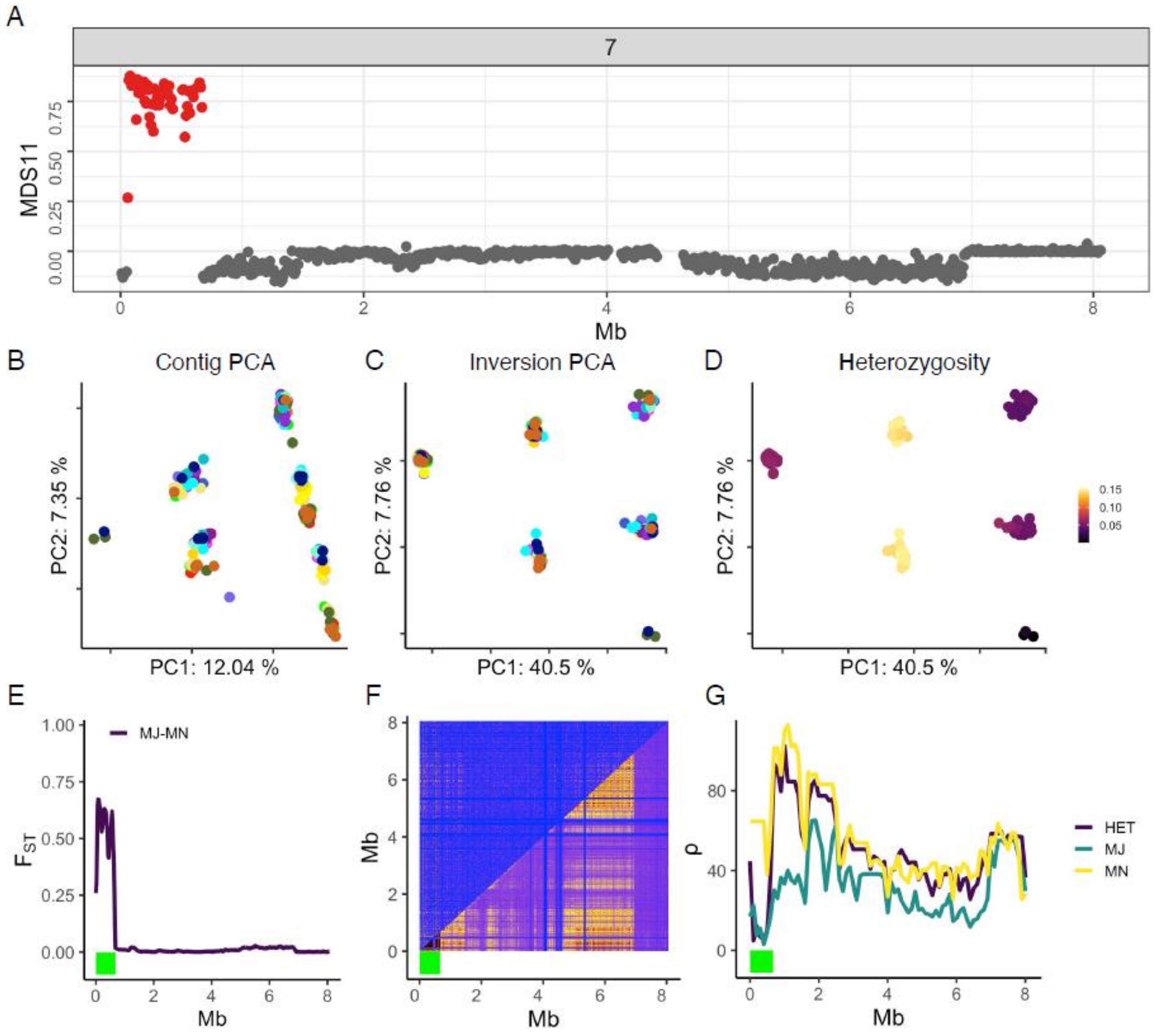

A
7
MDS11
0.00
0.25
0.50
0.75
0
2
4
6
8
Mb
B
Contig PCA
PC2: 7.35 %
PC1: 12.04 %
C
Inversion PCA
PC2: 7.76 %
PC1: 40.5 %
D
Heterozygosity
PC2: 7.76 %
PC1: 40.5 %
0.15
0.10
0.05
E
MJ-MN
1.00
0.75
0.50
0.25
0.00
Mb
F
Mb
Mb
G
ρ
80
40
0
HET
MJ
MN
Mb

**Inv7.2**

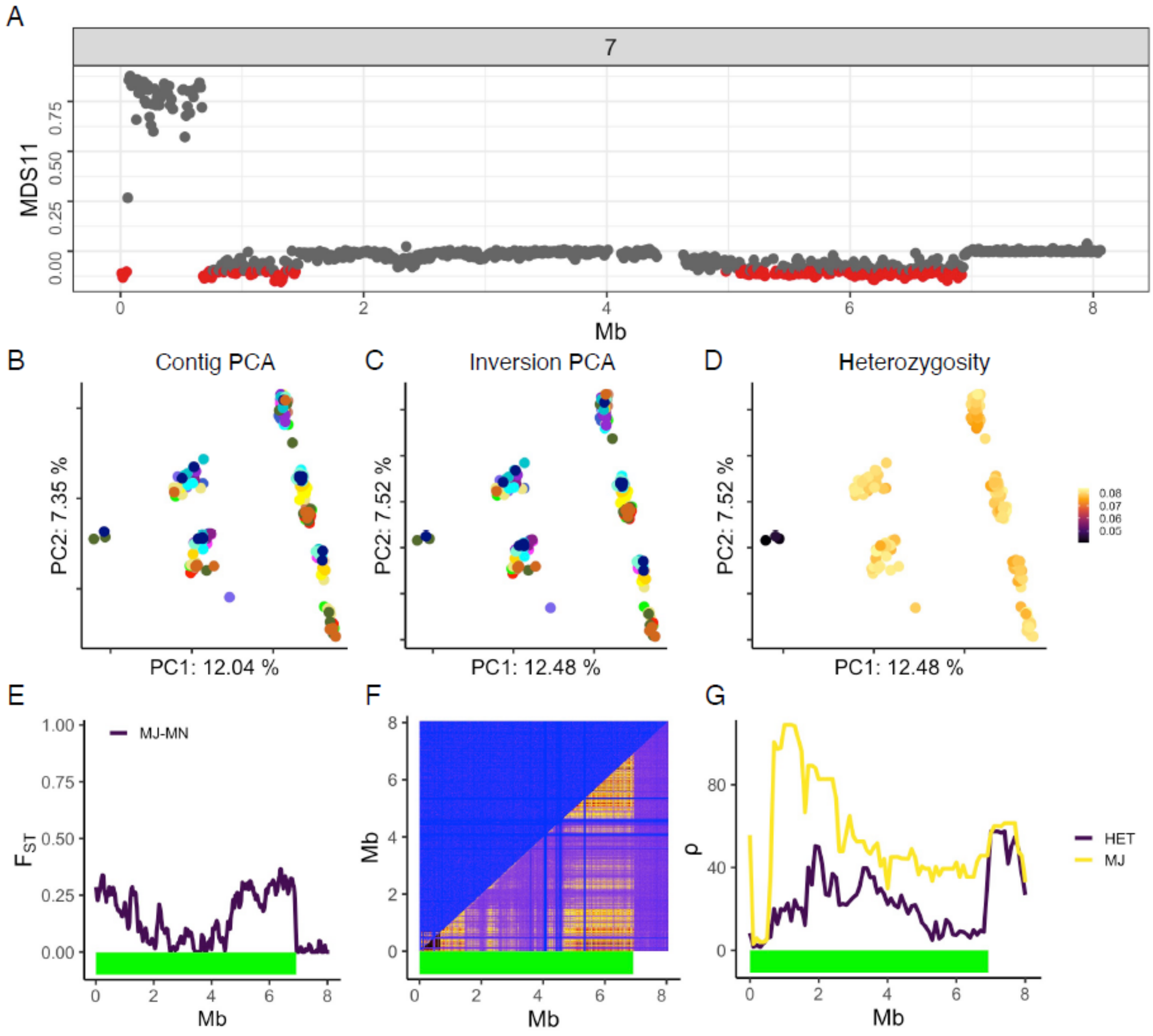

A
7
MDS11
0.00
0.25
0.50
0.75
0
2
4
6
8
Mb
B
Contig PCA
PC2: 7.35 %
PC1: 12.04 %
C
Inversion PCA
PC2: 7.52 %
PC1: 12.48 %
D
Heterozygosity
PC2: 7.52 %
PC1: 12.48 %
0.08
0.07
0.06
0.05
E
MJ-MN
1.00
0.75
0.50
0.25
0.00
$F_{ST}$
0
2
4
6
8
Mb
F
Mb
8
6
4
2
0
0
2
4
6
8
Mb
G
ρ
80
40
0
0
2
4
6
8
Mb
HET
MJ

**Inv9**

Inv9 is only polymorphic in northern group and PCAs are shown only for northern populations for clearer visualization. Population recombination rate was calculated using females only, as this is a putative X chromosome.

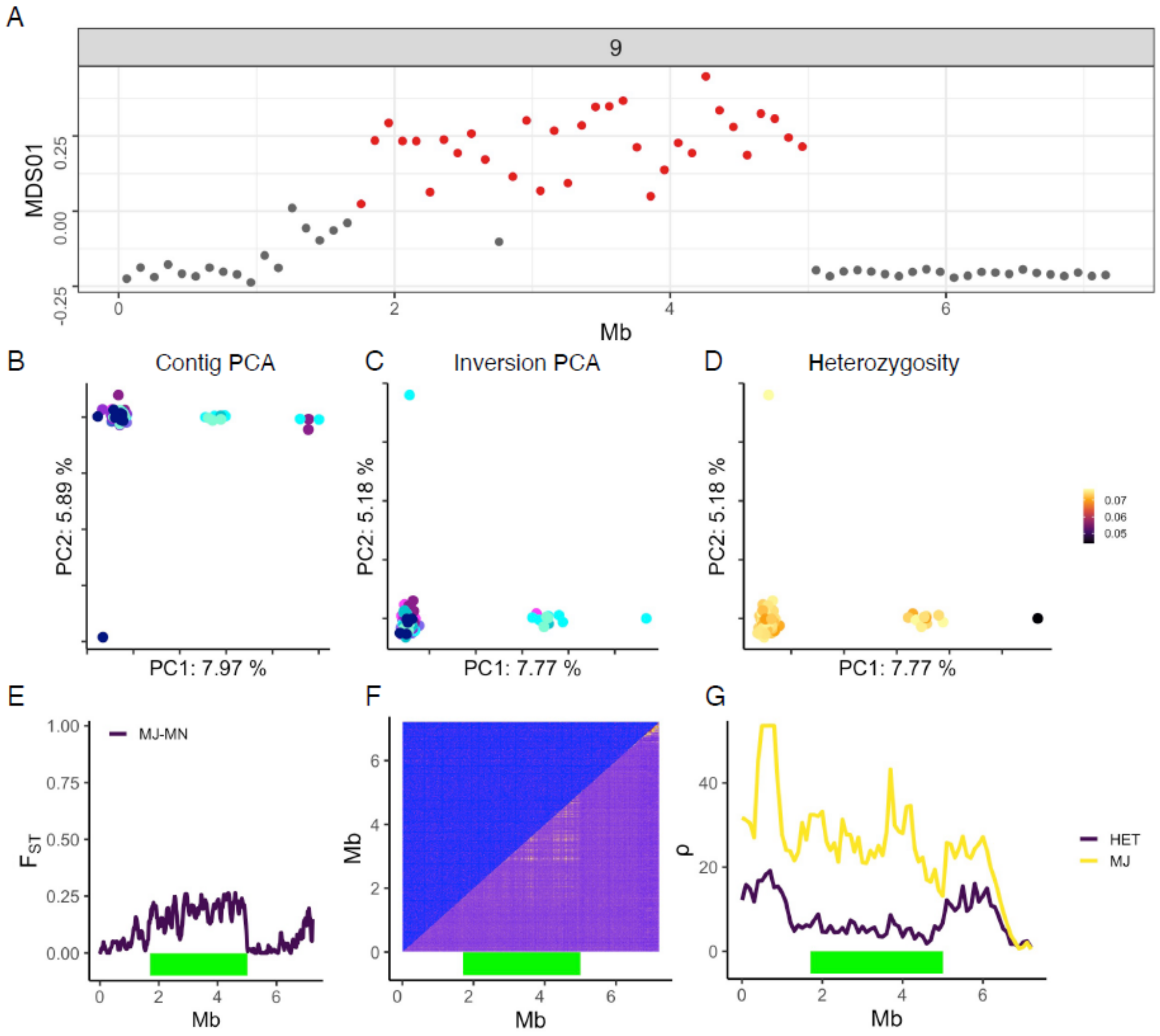

### Inv10

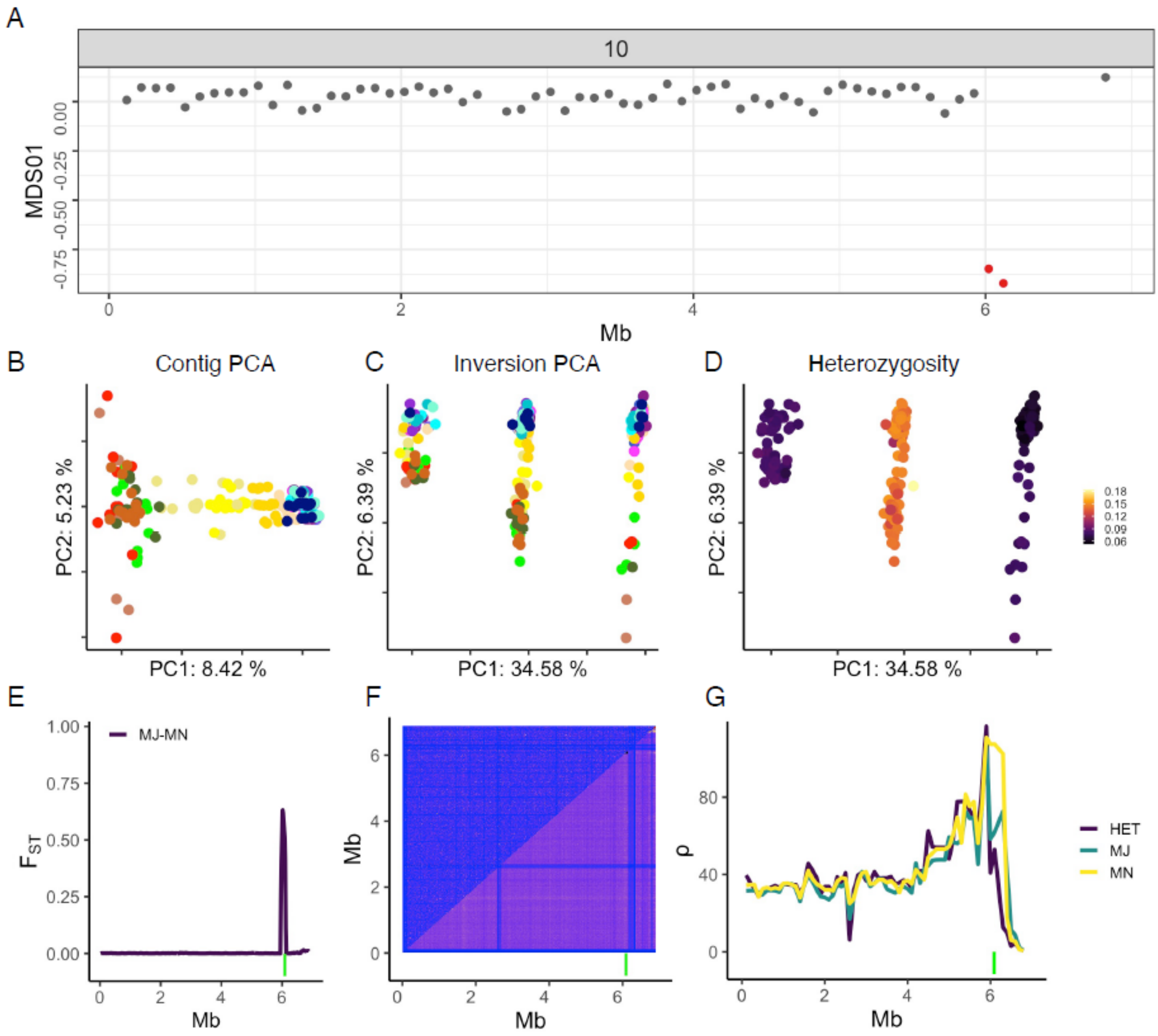

A
10
MDS01
0.00
-0.25
-0.50
-0.75
0
2
4
6
Mb
B
Contig PCA
PC2: 5.23 %
PC1: 8.42 %
C
Inversion PCA
PC2: 6.39 %
PC1: 34.58 %
D
Heterozygosity
PC2: 6.39 %
PC1: 34.58 %
0.18
0.15
0.12
0.09
0.06
E
MJ-MN
1.00
0.75
0.50
0.25
0.00
Mb
F
Mb
Mb
G
80
40
0
HET
MJ
MN
Mb

**Inv12**

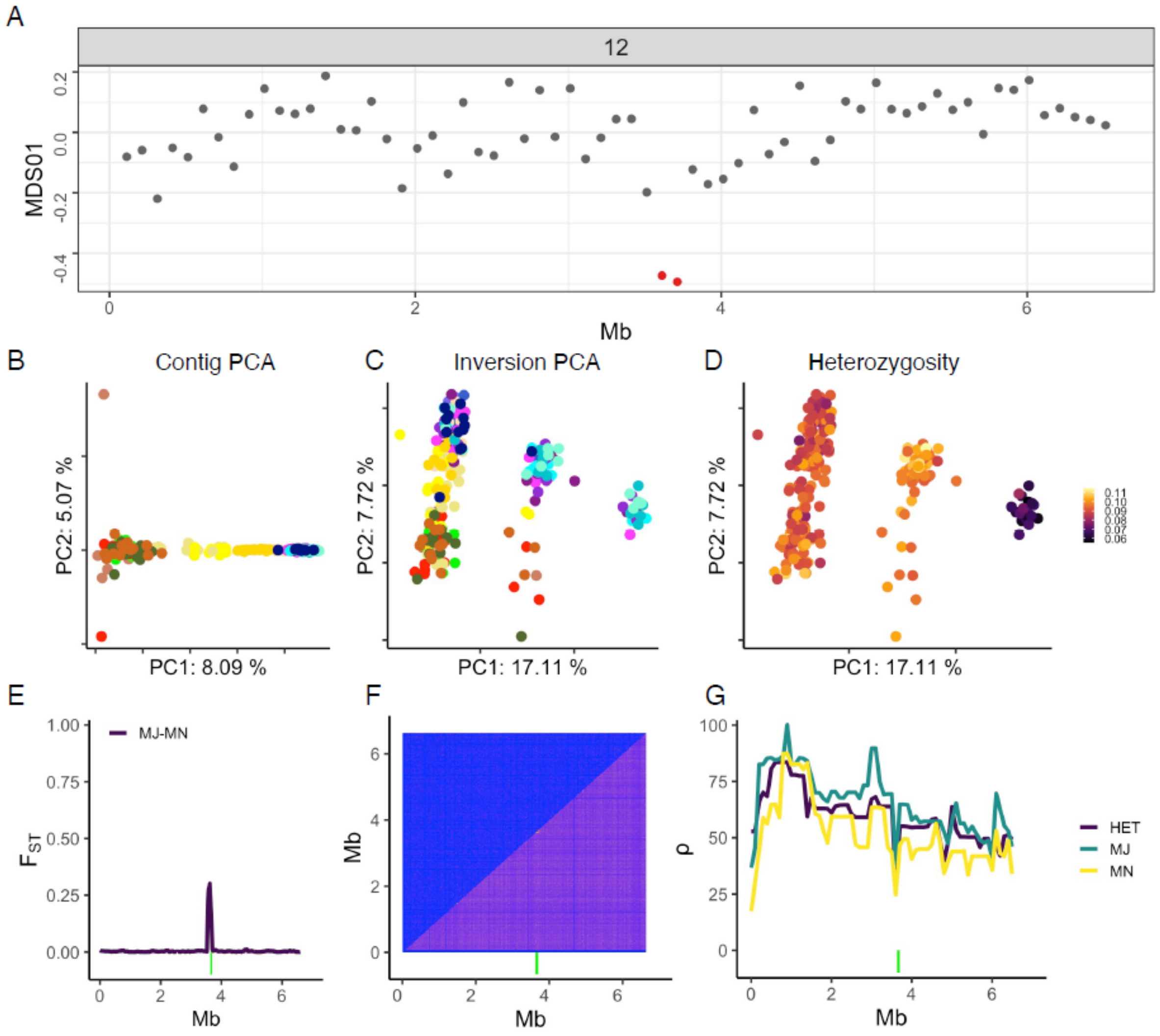

A
12
MDS01
0.2
0.0
-0.2
-0.4
0
2
4
6
Mb
B
Contig PCA
PC2: 5.07 %
PC1: 8.09 %
C
Inversion PCA
PC2: 7.72 %
PC1: 17.11 %
D
Heterozygosity
PC2: 7.72 %
PC1: 17.11 %
0.11
0.10
0.09
0.08
0.07
0.06
E
MJ-MN
$F_{ST}$
1.00
0.75
0.50
0.25
0.00
Mb
F
Mb
6
4
2
0
Mb
G
ρ
100
75
50
25
0
HET
MJ
MN
Mb

**Inv13**

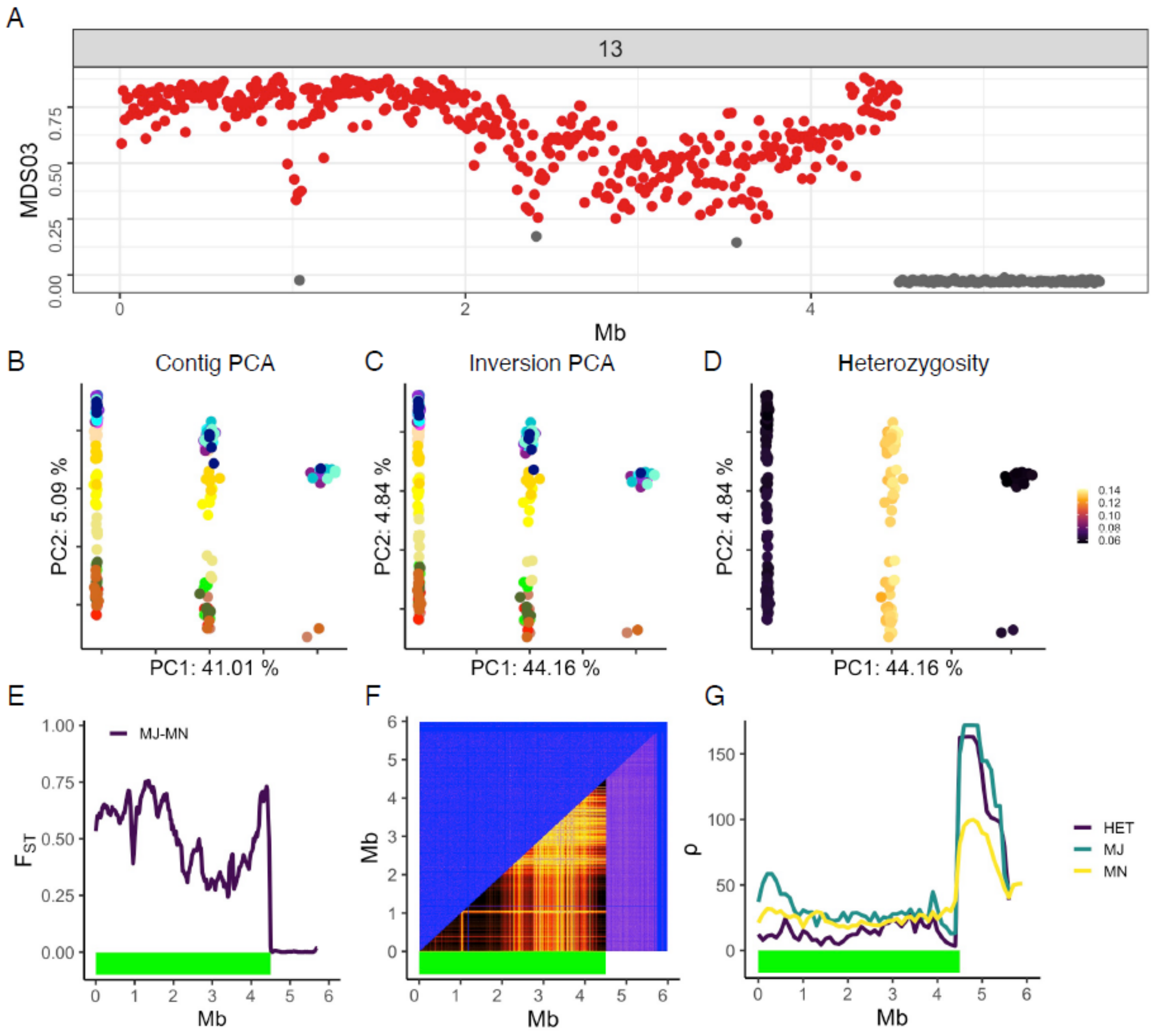

A
13
MDS03
0.00
0.25
0.50
0.75
0
2
4
Mb
B
Contig PCA
PC2: 5.09 %
PC1: 41.01 %
C
Inversion PCA
PC2: 4.84 %
PC1: 44.16 %
D
Heterozygosity
PC2: 4.84 %
PC1: 44.16 %
0.14
0.12
0.10
0.08
0.06
E
MJ-MN
$F_{ST}$
1.00
0.75
0.50
0.25
0.00
Mb
F
Mb
G
ρ
150
100
50
0
HET
MJ
MN
Mb

**Inv14.1**

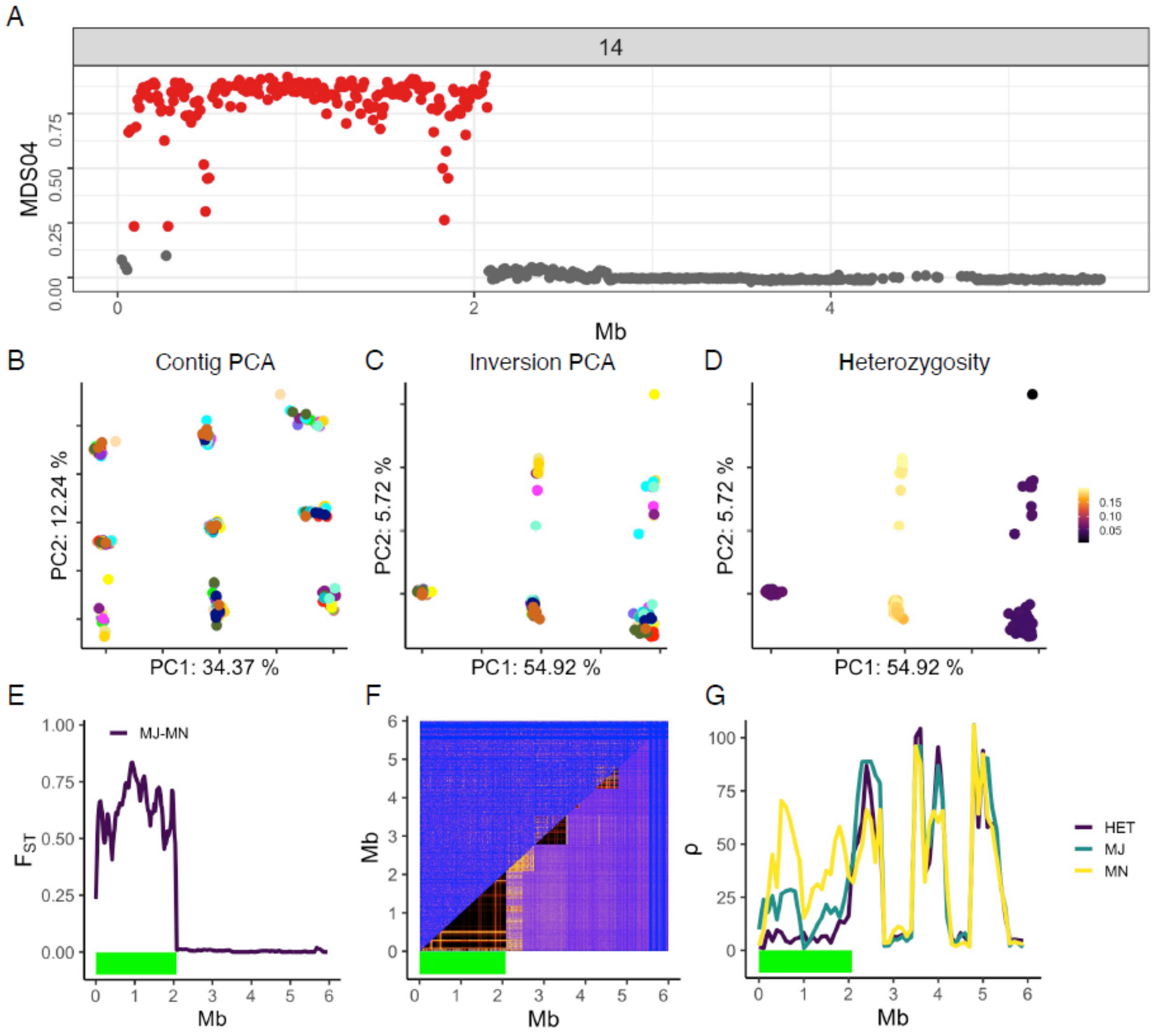

A
14
MDS04
0.00
0.25
0.50
0.75
0
2
4
Mb
B
Contig PCA
PC2: 12.24 %
PC1: 34.37 %
C
Inversion PCA
PC2: 5.72 %
PC1: 54.92 %
D
Heterozygosity
PC2: 5.72 %
PC1: 54.92 %
0.15
0.10
0.05
E
MJ-MN
$F_{ST}$
1.00
0.75
0.50
0.25
0.00
Mb
F
Mb
Mb
G
ρ
100
75
50
25
0
Mb
HET
MJ
MN

**Inv14.2**

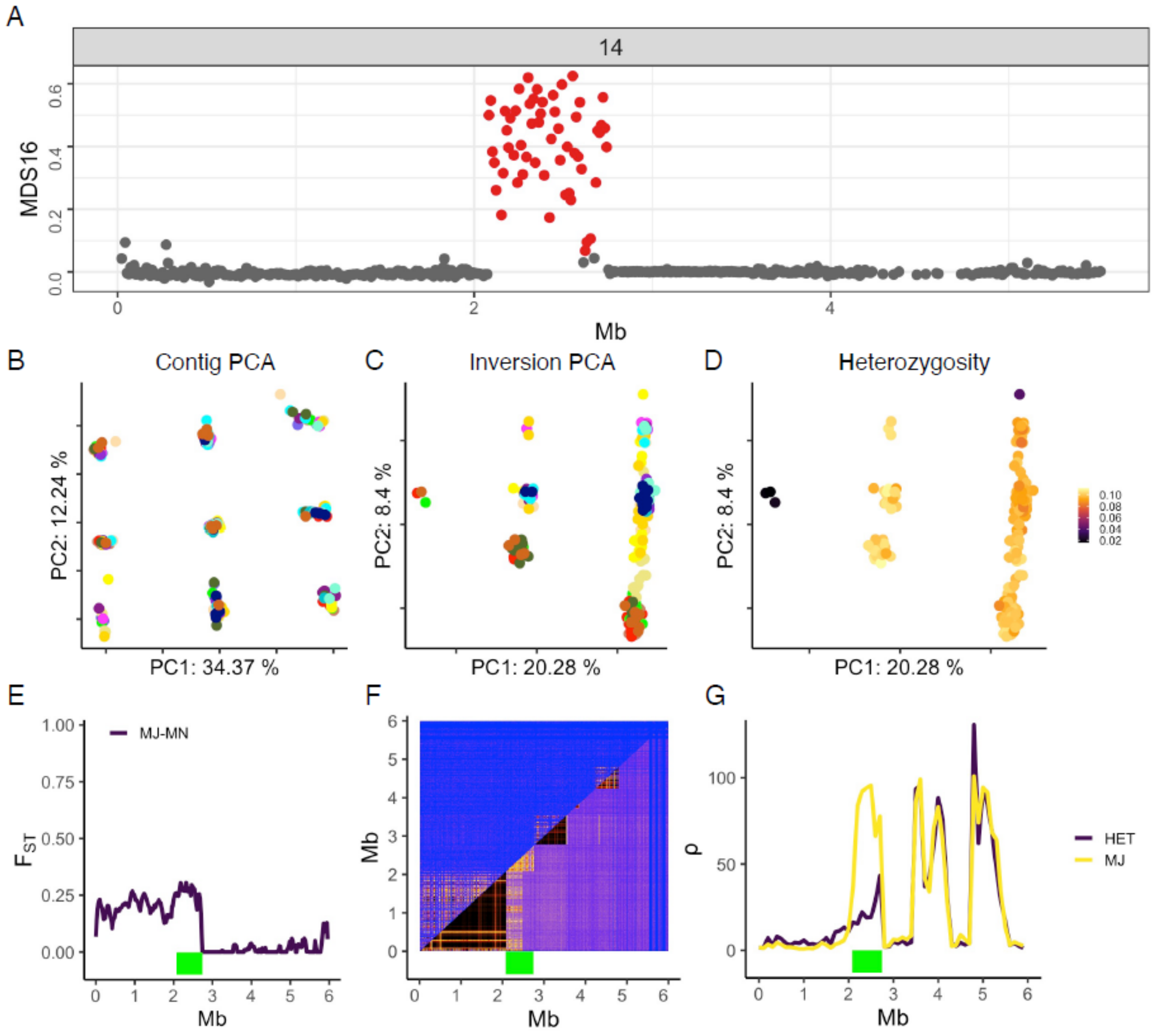

A
14
MDS16
0.0
0.2
0.4
0.6
0
2
4
Mb
B
Contig PCA
PC2: 12.24 %
PC1: 34.37 %
C
Inversion PCA
PC2: 8.4 %
PC1: 20.28 %
D
Heterozygosity
PC2: 8.4 %
PC1: 20.28 %
0.10
0.08
0.06
0.04
0.02
E
MJ-MN
1.00
0.75
0.50
0.25
0.00
$F_{ST}$
0 1 2 3 4 5 6
Mb
F
Mb
0 1 2 3 4 5 6
Mb
G
HET
MJ
ρ
100
50
0
0 1 2 3 4 5 6
Mb

**Inv14.3**

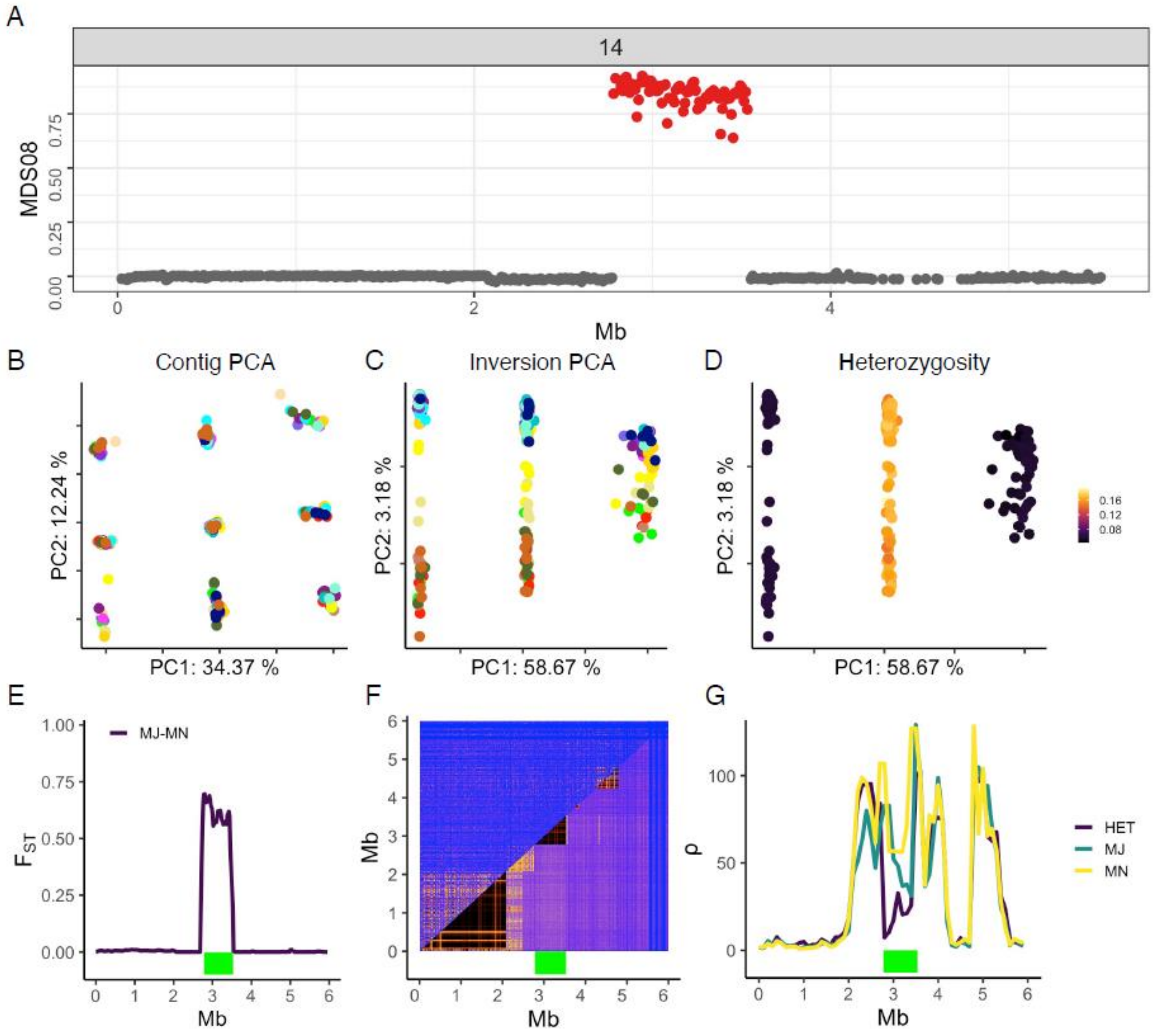
A
14
MDS08
0.00
0.25
0.50
0.75
0
2
4
Mb
B
Contig PCA
PC2: 12.24 %
PC1: 34.37 %
C
Inversion PCA
PC2: 3.18 %
PC1: 58.67 %
D
Heterozygosity
PC2: 3.18 %
PC1: 58.67 %
0.16
0.12
0.08
E
MJ-MN
FST
1.00
0.75
0.50
0.25
0.00
Mb
F
Mb
Mb
G
ρ
100
50
0
Mb
HET
MJ
MN

**Inv14.4**

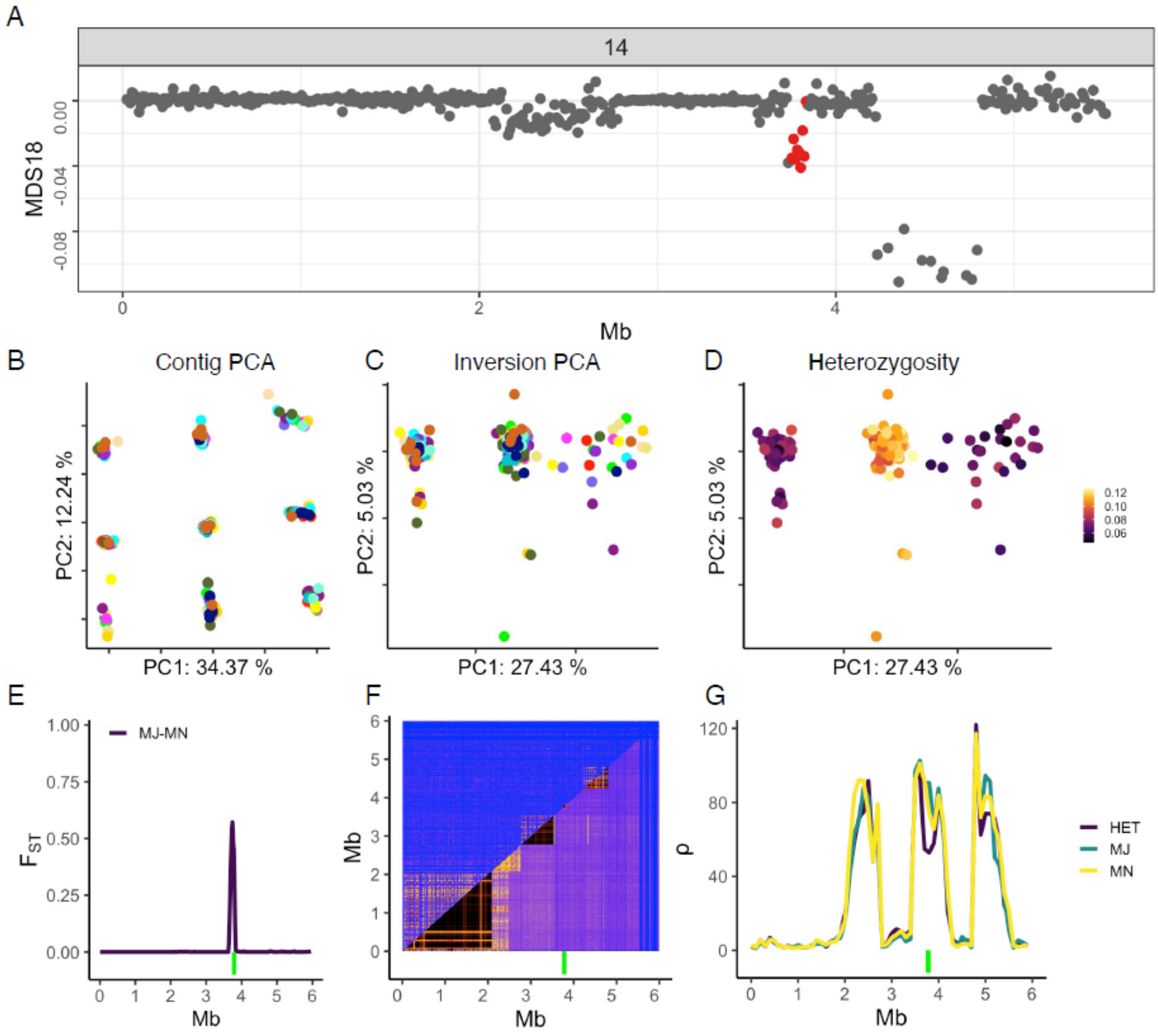

A
14
MDS18
0.00
-0.04
-0.08
0
2
4
Mb
B
Contig PCA
PC2: 12.24 %
PC1: 34.37 %
C
Inversion PCA
PC2: 5.03 %
PC1: 27.43 %
D
Heterozygosity
PC2: 5.03 %
PC1: 27.43 %
0.12
0.10
0.08
0.06
E
MJ-MN
1.00
0.75
0.50
0.25
0.00
$F_{ST}$
0 1 2 3 4 5 6
Mb
F
6 5 4 3 2 1 0
Mb
0 1 2 3 4 5 6
Mb
G
120
80
40
0
ρ
0 1 2 3 4 5 6
Mb
HET
MJ
MN

**Inv14.5**

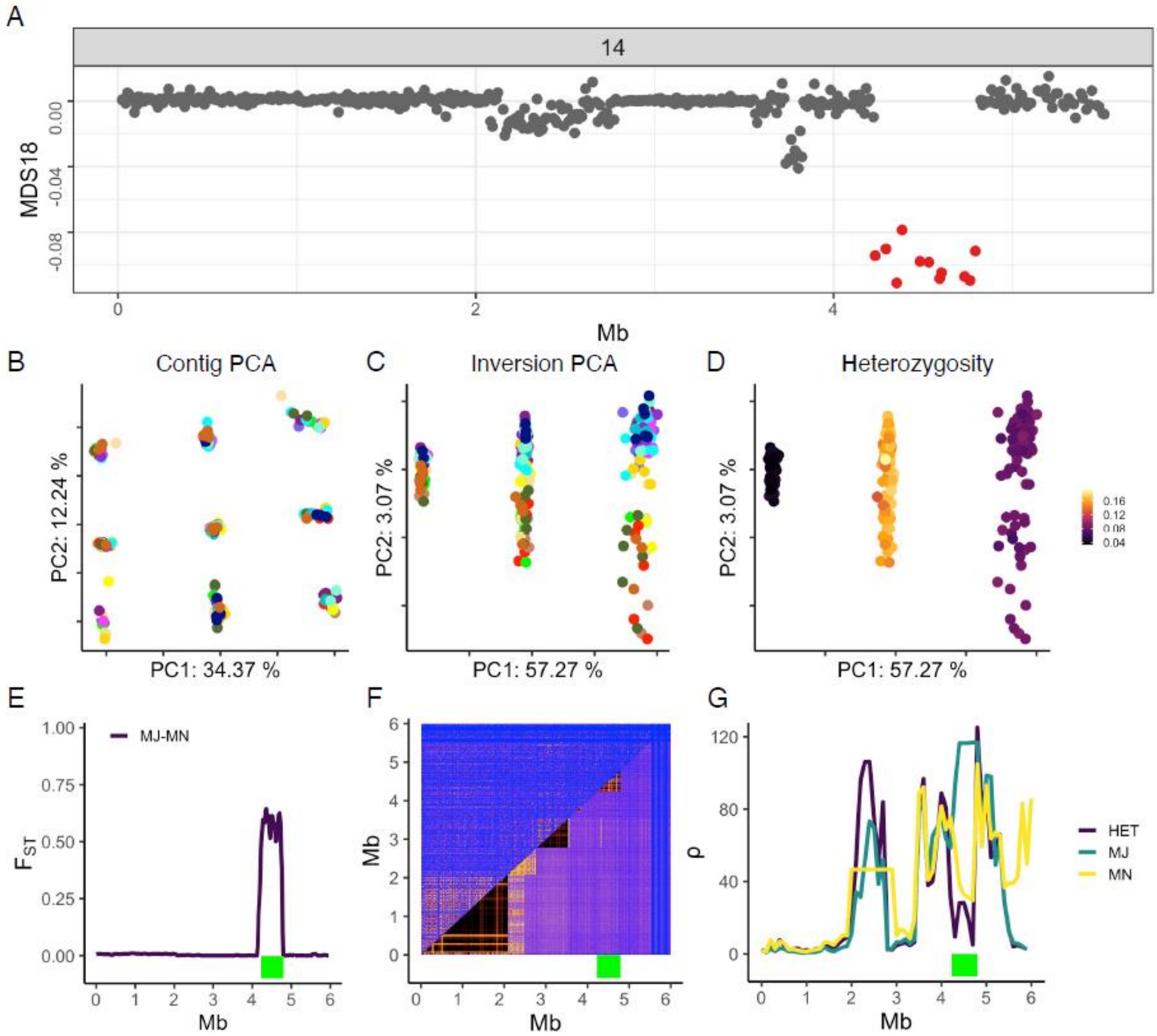

A
14
MDS18
0.00
-0.04
-0.08
0
2
4
Mb
B
Contig PCA
PC2: 12.24 %
PC1: 34.37 %
C
Inversion PCA
PC2: 3.07 %
PC1: 57.27 %
D
Heterozygosity
PC2: 3.07 %
PC1: 57.27 %
0.16
0.12
0.08
0.04
E
MJ-MN
1.00
0.75
0.50
0.25
0.00
$F_{ST}$
Mb
F
Mb
Mb
G
120
80
40
0
ρ
HET
MJ
MN
Mb

**Inv15**

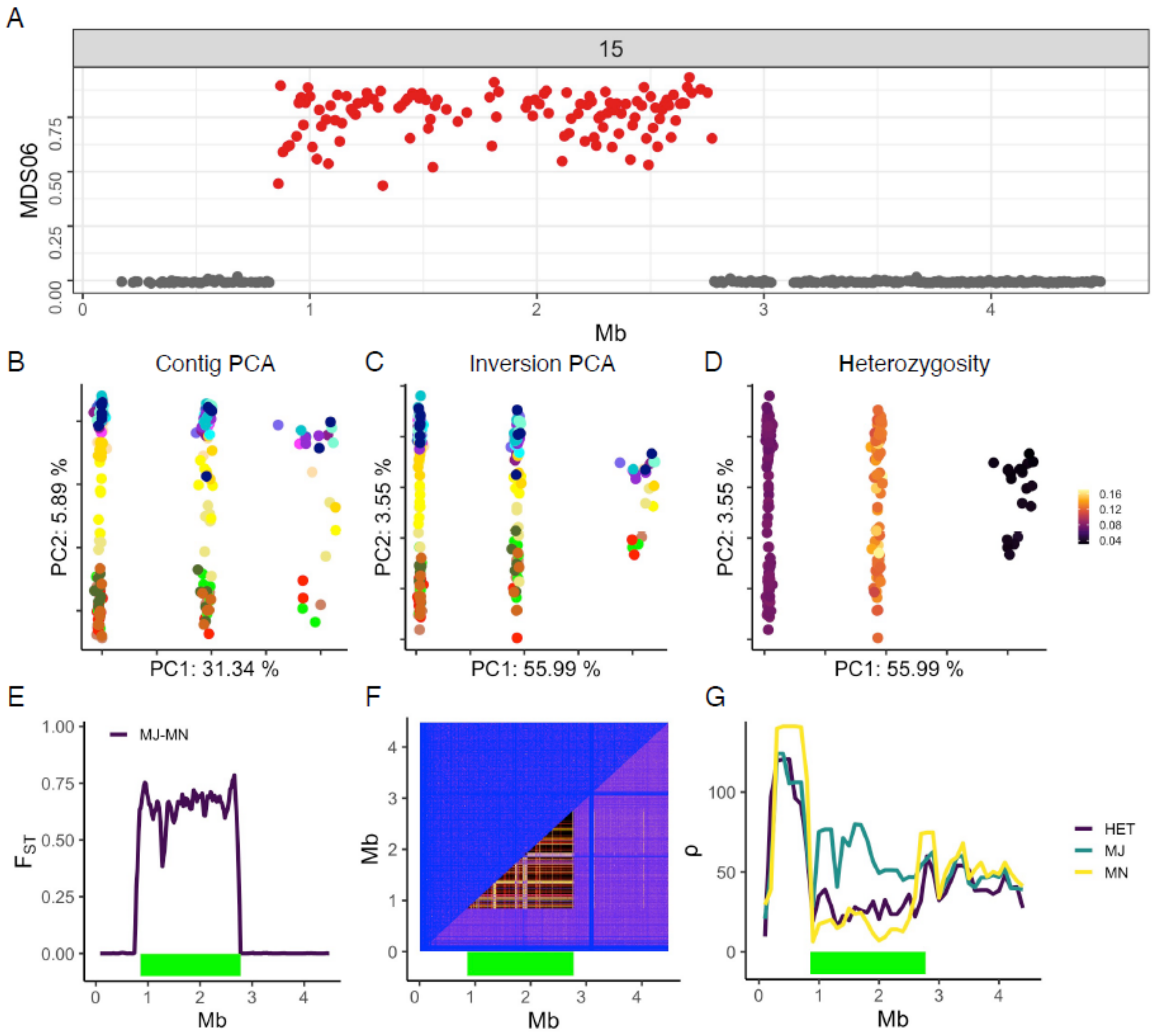
A
15
MDS06
0.00
0.25
0.50
0.75
0
1
2
3
4
Mb
B
Contig PCA
PC2: 5.89 %
PC1: 31.34 %
C
Inversion PCA
PC2: 3.55 %
PC1: 55.99 %
D
Heterozygosity
PC2: 3.55 %
PC1: 55.99 %
0.16
0.12
0.08
0.04
E
MJ-MN
1.00
0.75
0.50
0.25
0.00
$F_{ST}$
Mb
F
Mb
Mb
G
HET
MJ
MN
100
50
0
ρ
Mb

**Inv16.1**

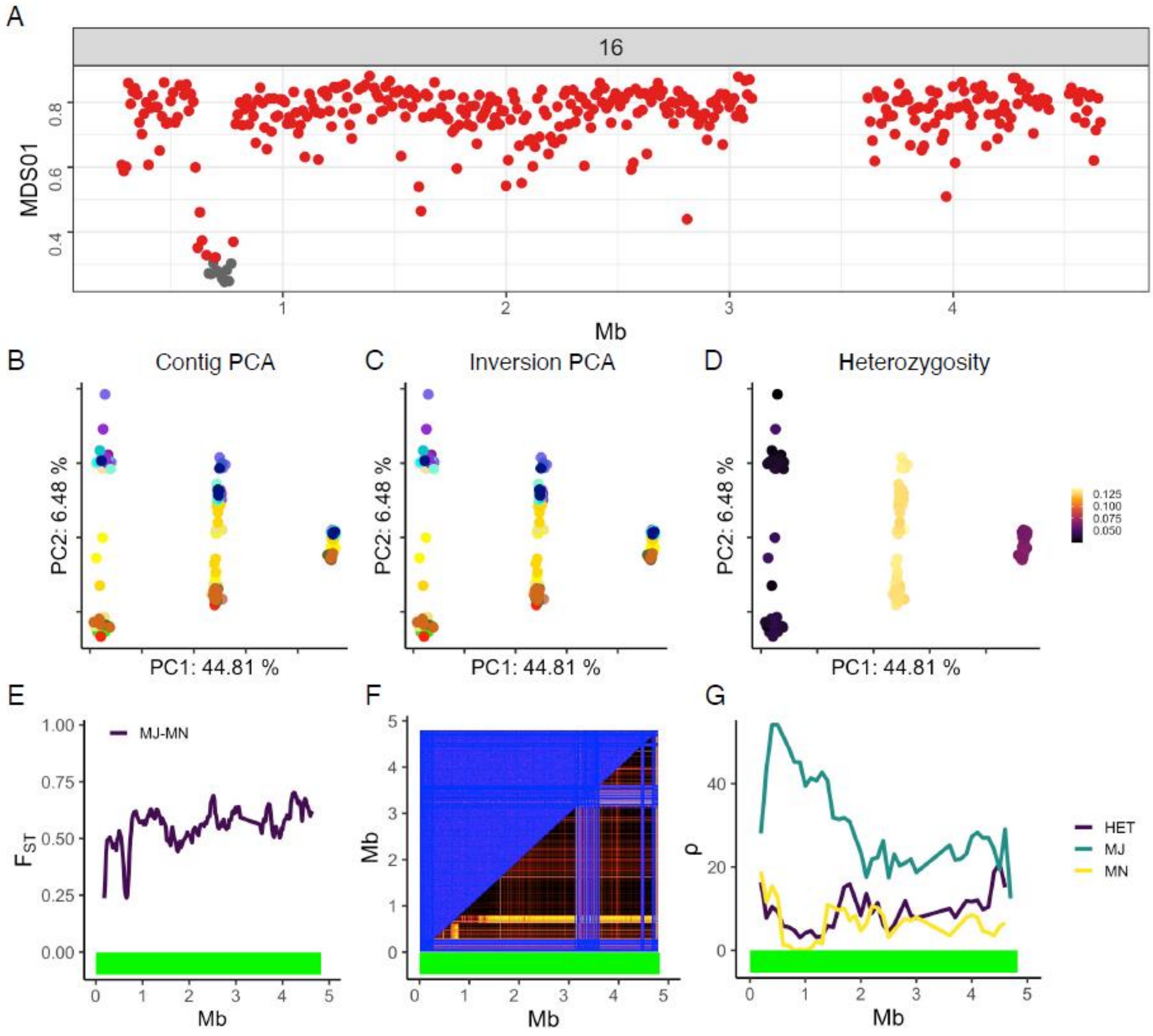

A
16
MDS01
Mb
B
Contig PCA
C
Inversion PCA
D
Heterozygosity
PC2: 6.48 %
PC1: 44.81 %
0.125
0.100
0.075
0.050
E
MJ-MN
$F_{ST}$
F
G
HET
MJ
MN
ρ

**Inv17**

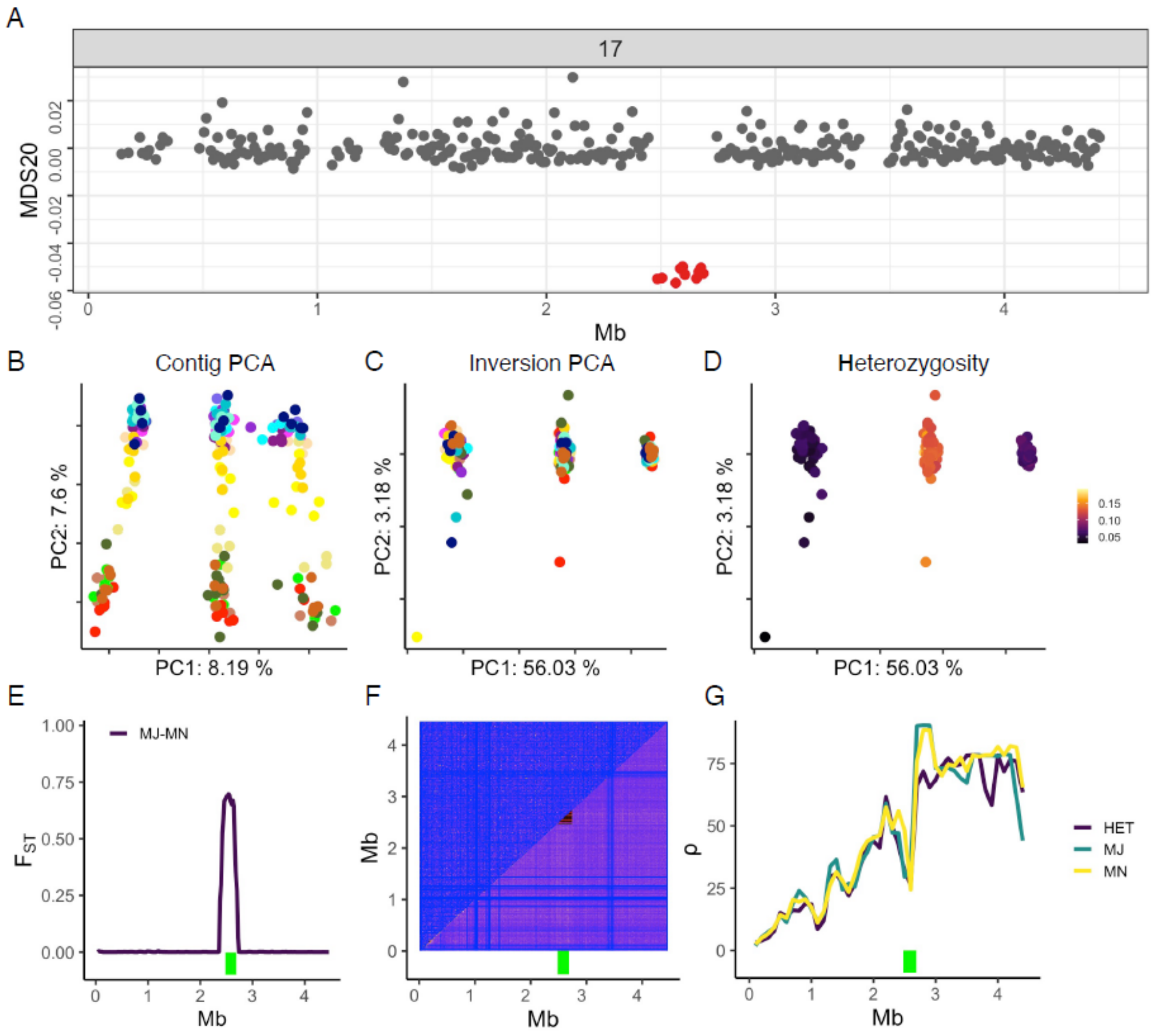
A
17
MDS20
0.02
0.00
-0.02
-0.04
-0.06
0
1
2
3
4
Mb
B
Contig PCA
PC2: 7.6 %
PC1: 8.19 %
C
Inversion PCA
PC2: 3.18 %
PC1: 56.03 %
D
Heterozygosity
PC2: 3.18 %
PC1: 56.03 %
0.15
0.10
0.05
E
MJ-MN
1.00
0.75
0.50
0.25
0.00
F
Mb
G
75
50
25
0
HET
MJ
MN

### Inv18

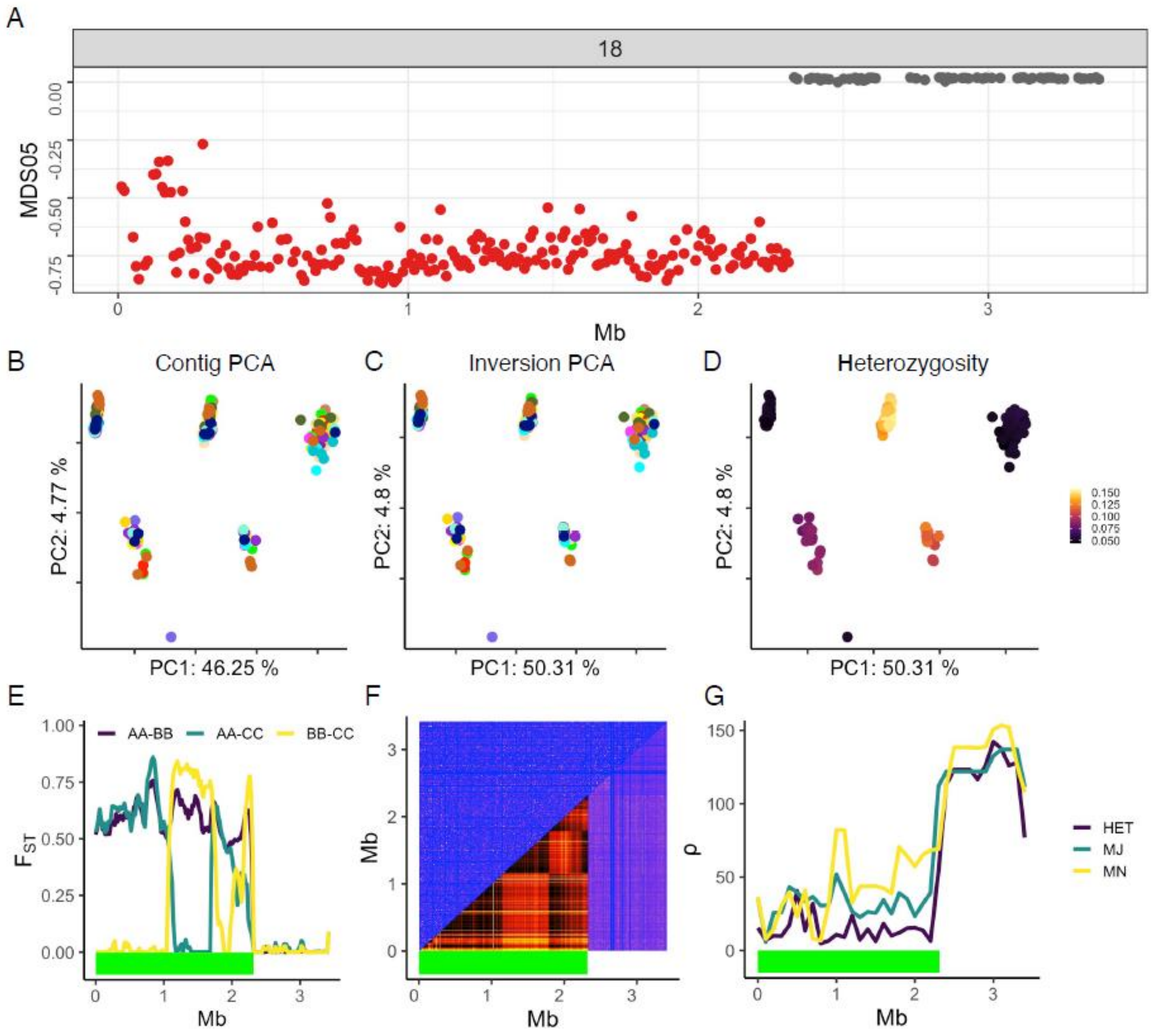

A
18
MDS05
0.00
-0.25
-0.50
-0.75
0
1
2
3
Mb
B
Contig PCA
PC2: 4.77 %
PC1: 46.25 %
C
Inversion PCA
PC2: 4.8 %
PC1: 50.31 %
D
Heterozygosity
PC2: 4.8 %
PC1: 50.31 %
0.150
0.125
0.100
0.075
0.050
E
AA-BB
AA-CC
BB-CC
1.00
0.75
0.50
0.25
0.00
F
G
150
100
50
HET
MJ
MN

### Inv22.1

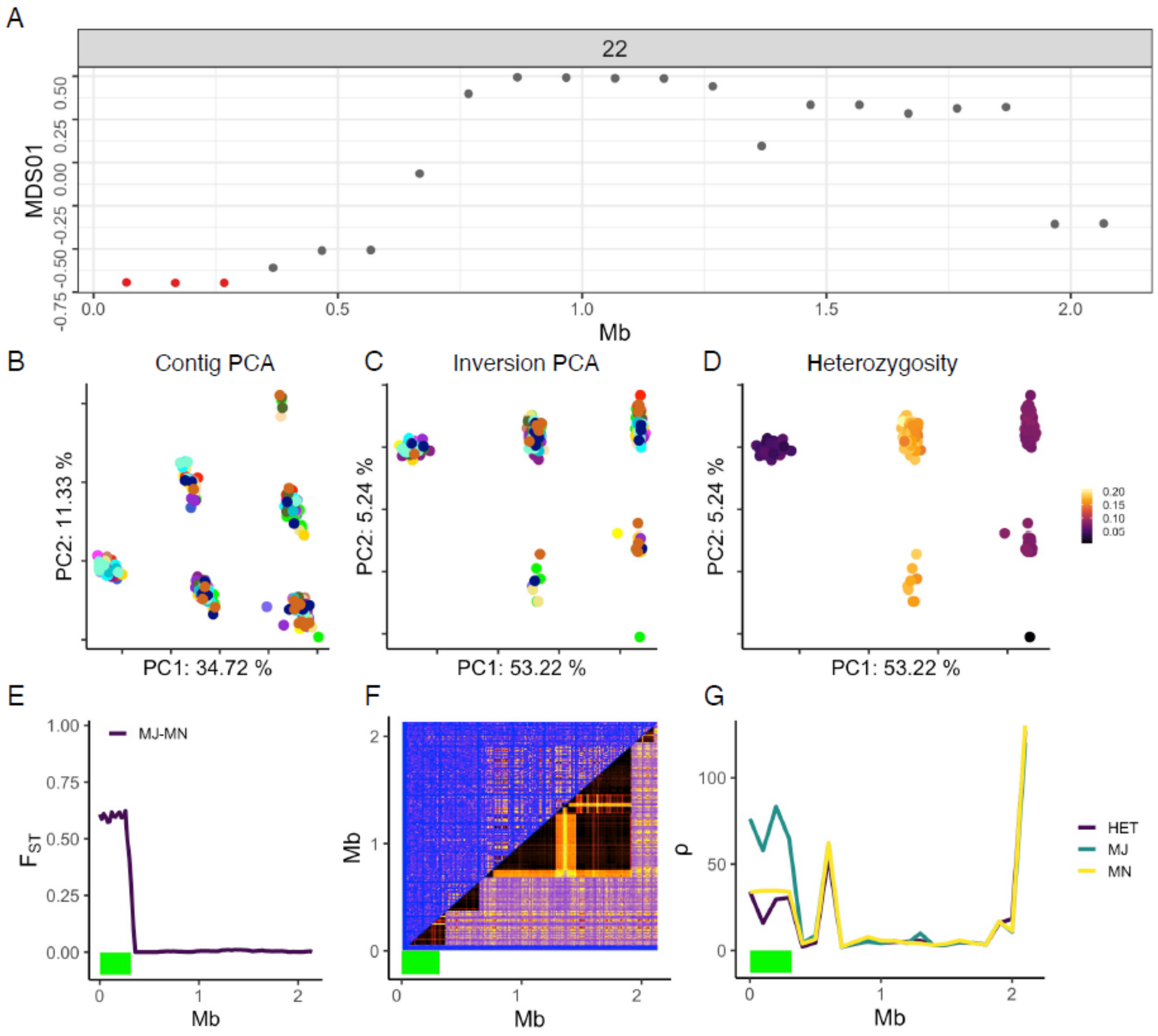

A
22
MDS01
Mb
B
Contig PCA
PC2: 11.33 %
PC1: 34.72 %
C
Inversion PCA
PC2: 5.24 %
PC1: 53.22 %
D
Heterozygosity
PC2: 5.24 %
PC1: 53.22 %
E
MJ-MN
$F_{ST}$
Mb
F
Mb
Mb
G
ρ
HET
MJ
MN
Mb

**Inv22.2**

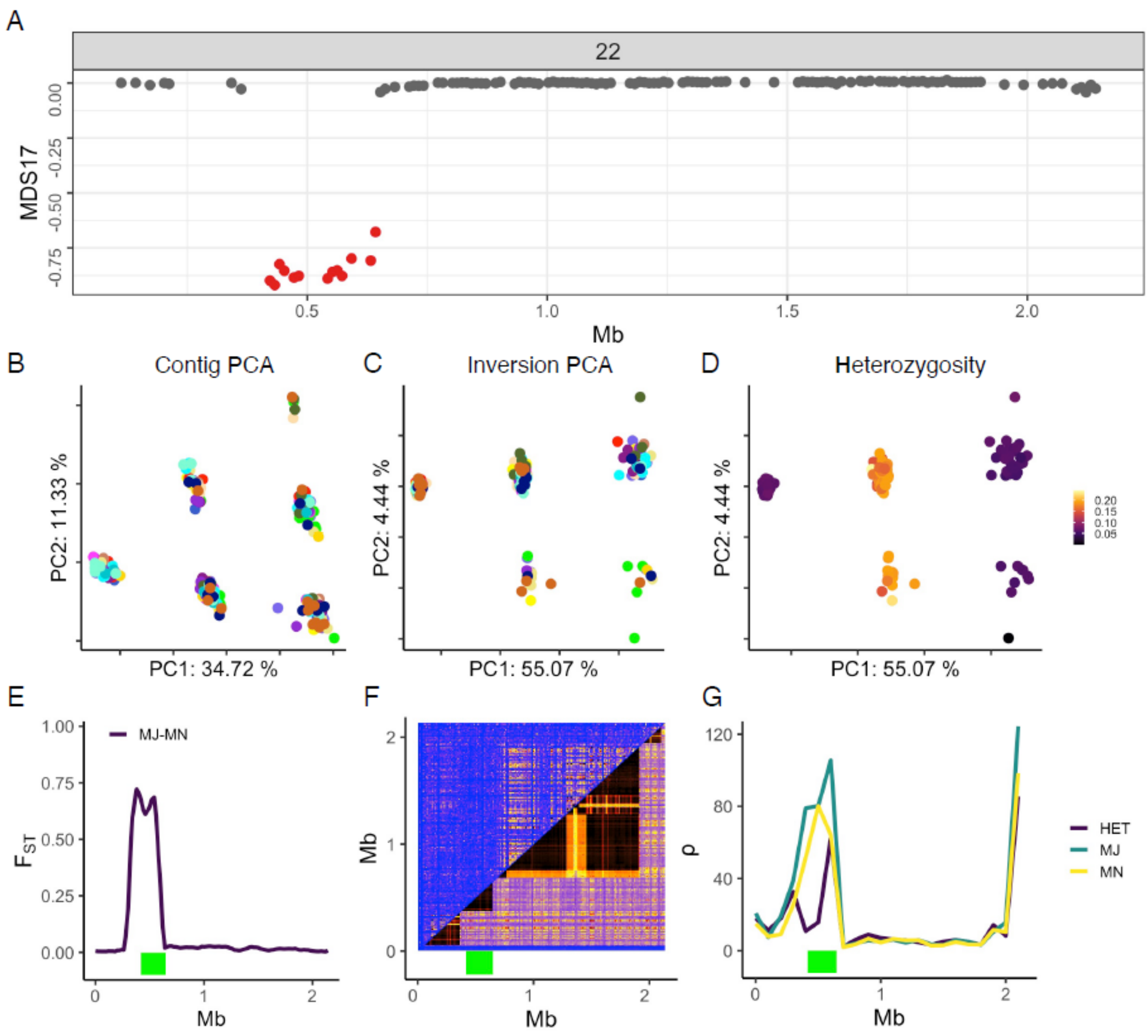
A
22
MDS17
0.00
-0.25
-0.50
-0.75
0.5
1.0
1.5
2.0
Mb
B
Contig PCA
PC2: 11.33 %
PC1: 34.72 %
C
Inversion PCA
PC2: 4.44 %
PC1: 55.07 %
D
Heterozygosity
PC2: 4.44 %
PC1: 55.07 %
0.20
0.15
0.10
0.05
E
MJ-MN
1.00
0.75
0.50
0.25
0.00
0
1
2
Mb
F
2
1
0
Mb
0
1
2
Mb
G
120
80
40
0
ρ
HET
MJ
MN
0
1
2
Mb

**Inv22.3**

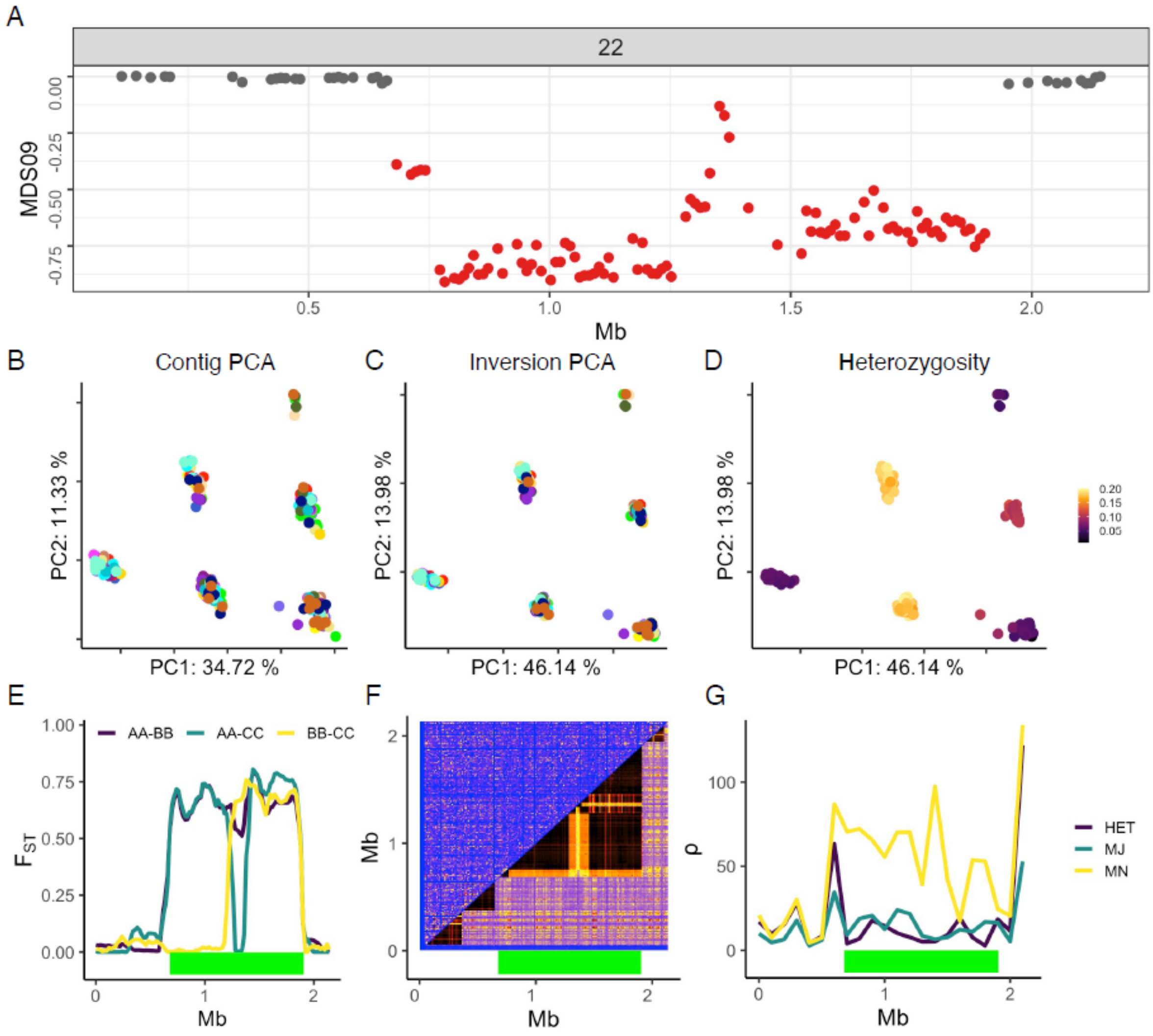

A
22
MDS09
0.00
-0.25
-0.50
-0.75
0.5
1.0
1.5
2.0
Mb
B
Contig PCA
PC2: 11.33 %
PC1: 34.72 %
C
Inversion PCA
PC2: 13.98 %
PC1: 46.14 %
D
Heterozygosity
PC2: 13.98 %
PC1: 46.14 %
0.20
0.15
0.10
0.05
E
AA-BB
AA-CC
BB-CC
1.00
0.75
0.50
0.25
0.00
F
ST
0
1
2
Mb
F
2
1
0
Mb
0
1
2
Mb
G
100
50
0
ρ
0
1
2
Mb
HET
MJ
MN

**Inv22.4**

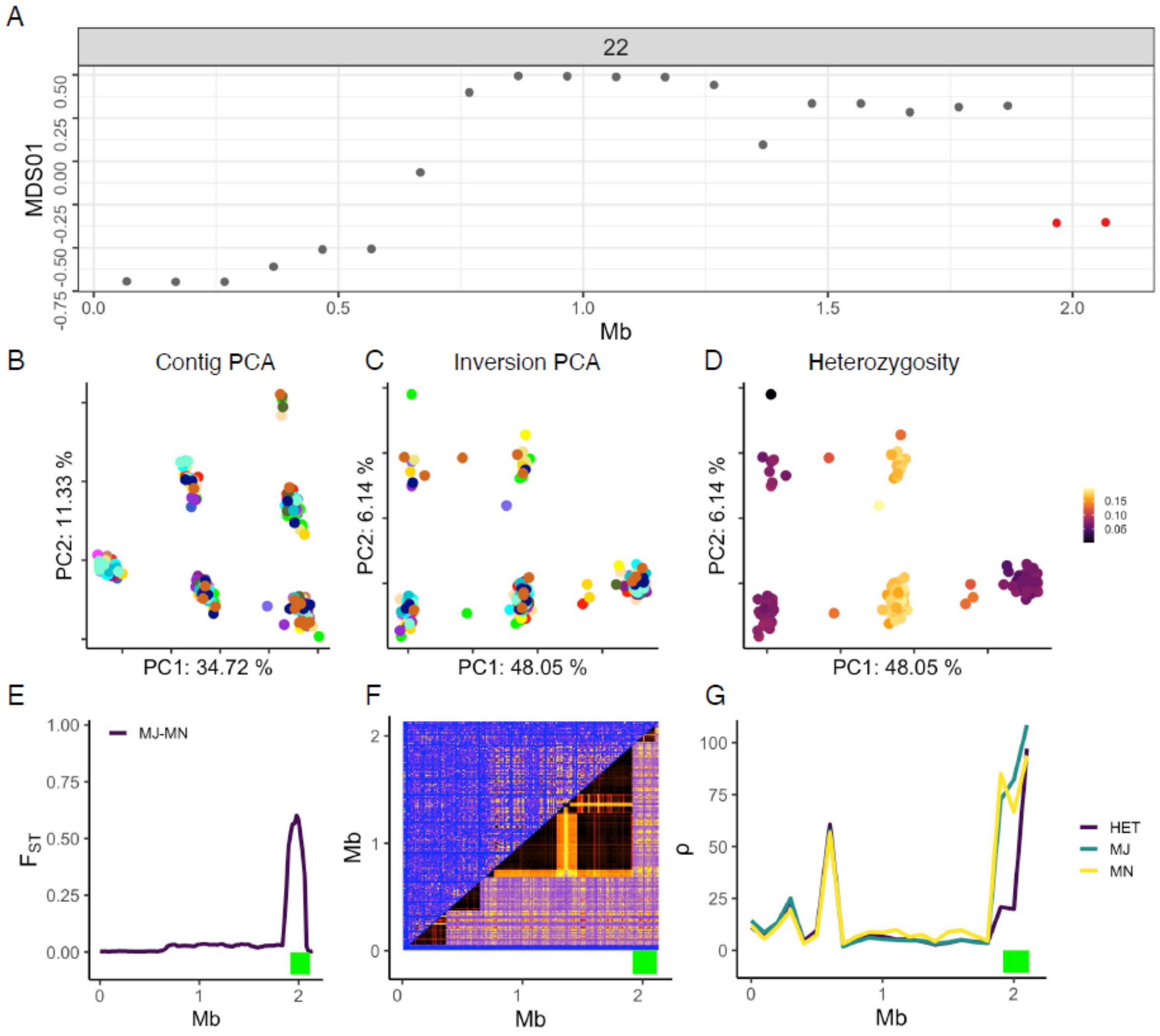

A
22
MDS01
-0.75 -0.50 -0.25 0.00 0.25 0.50
0.0 0.5 1.0 1.5 2.0
Mb
B Contig PCA
PC2: 11.33 %
PC1: 34.72 %
C Inversion PCA
PC2: 6.14 %
PC1: 48.05 %
D Heterozygosity
PC2: 6.14 %
PC1: 48.05 %
0.15
0.10
0.05
E
1.00 0.75 0.50 0.25 0.00
MJ-MN
$F_{ST}$
0 1 2
Mb
F
Mb
0 1 2
Mb
G
100 75 50 25 0
ρ
0 1 2
Mb
HET
MJ
MN

**Inv23.1**

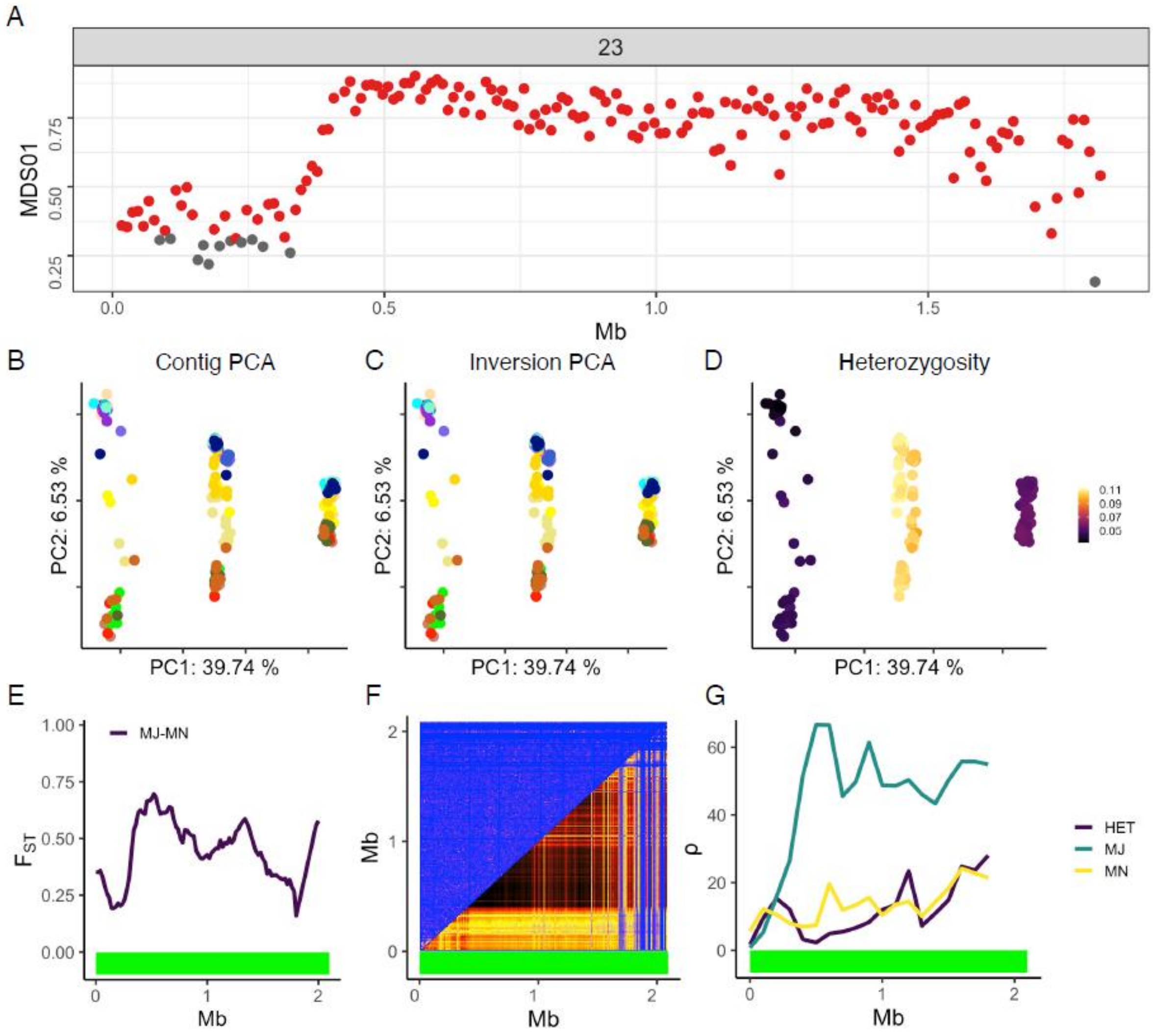

A
23
MDS01
0.25
0.50
0.75
0.0
0.5
1.0
1.5
Mb
B
Contig PCA
PC2: 6.53 %
PC1: 39.74 %
C
Inversion PCA
PC2: 6.53 %
PC1: 39.74 %
D
Heterozygosity
PC2: 6.53 %
PC1: 39.74 %
0.11
0.09
0.07
0.05
E
MJ-MN
1.00
0.75
0.50
0.25
0.00
$F_{ST}$
0
1
2
Mb
F
Mb
2
1
0
0
1
2
Mb
G
60
40
20
0
ρ
0
1
2
Mb
HET
MJ
MN

**Inv26**

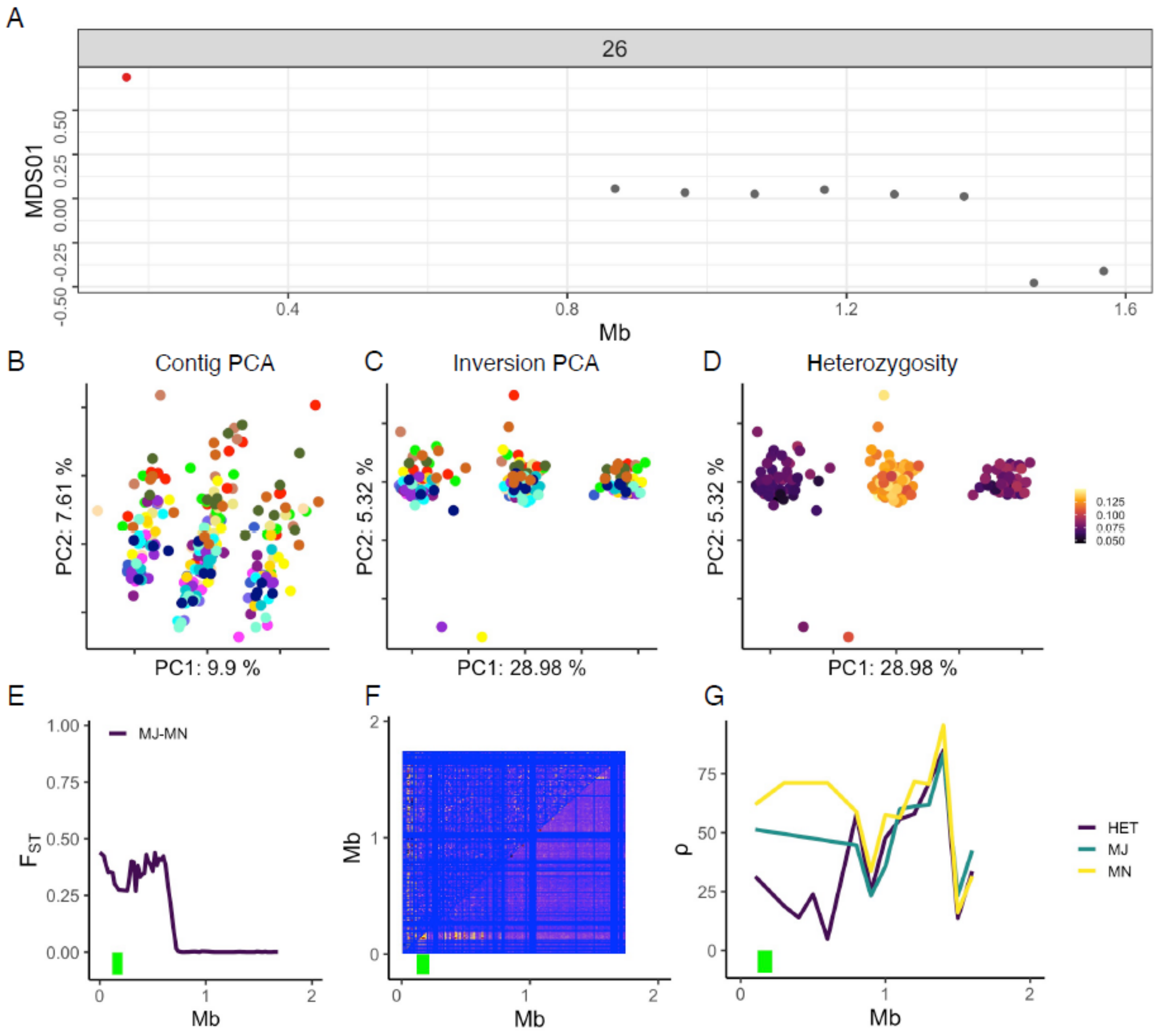

A
26
MDS01
-0.50
-0.25
0.00
0.25
0.50
0.4
0.8
1.2
1.6
Mb
B
Contig PCA
PC2: 7.61 %
PC1: 9.9 %
C
Inversion PCA
PC2: 5.32 %
PC1: 28.98 %
D
Heterozygosity
PC2: 5.32 %
PC1: 28.98 %
0.125
0.100
0.075
0.050
E
MJ-MN
1.00
0.75
0.50
0.25
0.00
$F_{ST}$
0
1
2
Mb
F
Mb
2
1
0
0
1
2
Mb
G
ρ
75
50
25
0
0
1
2
Mb
HET
MJ
MN

**Figure S3.** Identification of putative double-crossover events and overlapping inversions.

#### Double-crossover patterns

Plots show linkage disequilibrium (LD) patterns in a contig of interest with a green bar indicating inversion position (A); PCA for inversion region with clusters colored by genotype (B) and heterozygosity (C). Inversion haplotypes are labeled A, B, and C (haplotype created by a putative double crossover event). Double-crossover patterns are characterized by a cross in lighter (lower LD) in the middle of the high LD region and six groups in PCA that correspond to three haplotypes and create six genotypes with heterozygosity patterns that match expectations: homozygotes with lower heterozygosity than heterozygotes.

#### Inv18

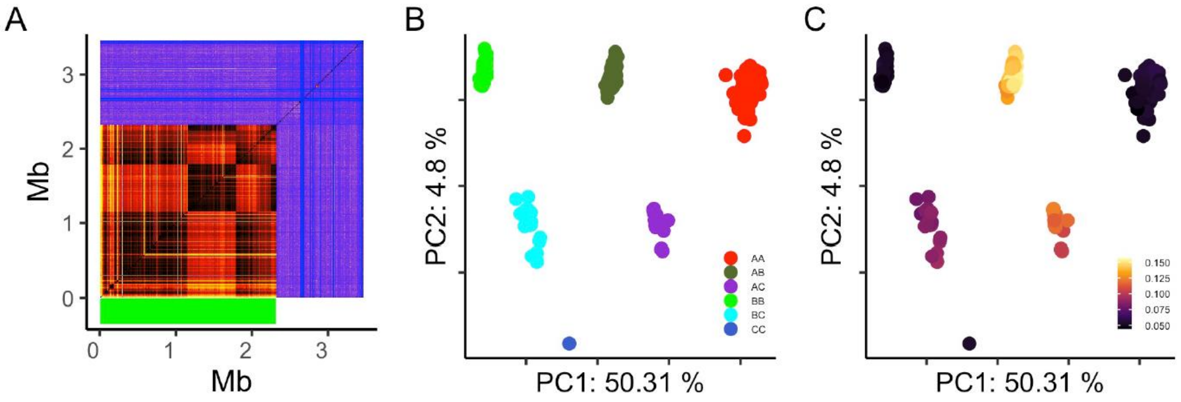


#### Inv22.3

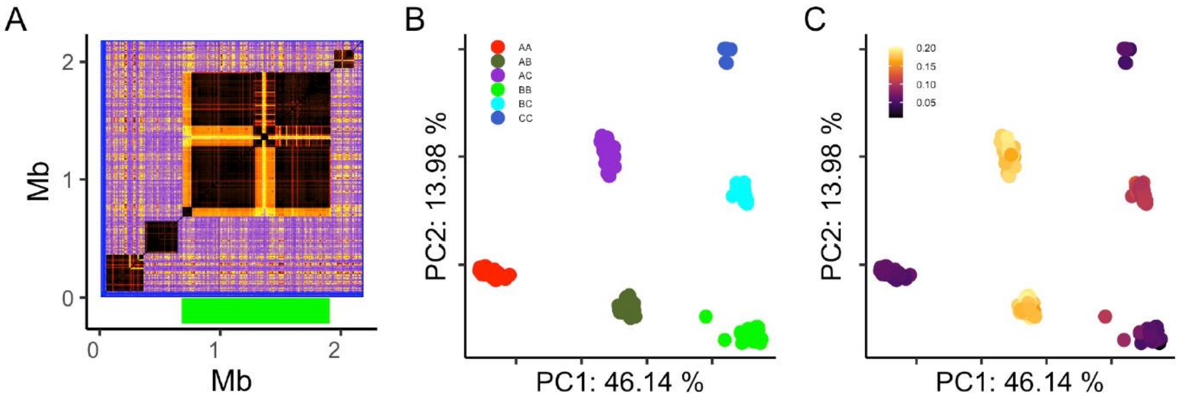

**Inv22.4**

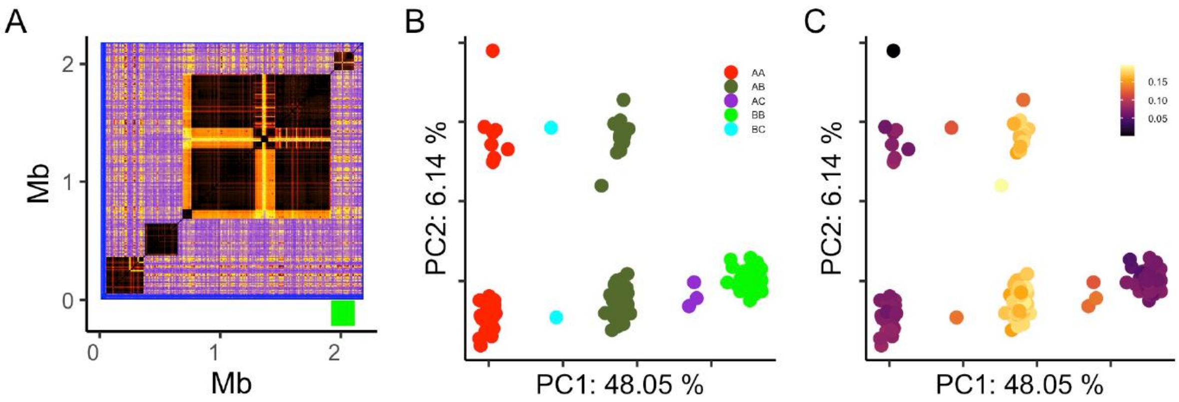


**Overlapping inversions**

**Inv7.2**

Inv7.2 overlaps with Inv7.1 (at the beginning of the contig). Plots show linkage disequilibrium (LD) patterns in a contig of interest with a green bar indicating Inv7.2 position (A), PCA for the part of Inv7.2 that overlaps with Inv7.1 (B) and the PCA for the nonoverlapping part of Inv7.2. Both PCAs are colored according to inversion genotype. We genotyped Inv7.2 based on PC2 from (B) and PC1 clustering from (C). Both genotyping methods provided the same results. See FigureS1 for more information on both inversions. AA, AB, BB – different inversion genotypes.

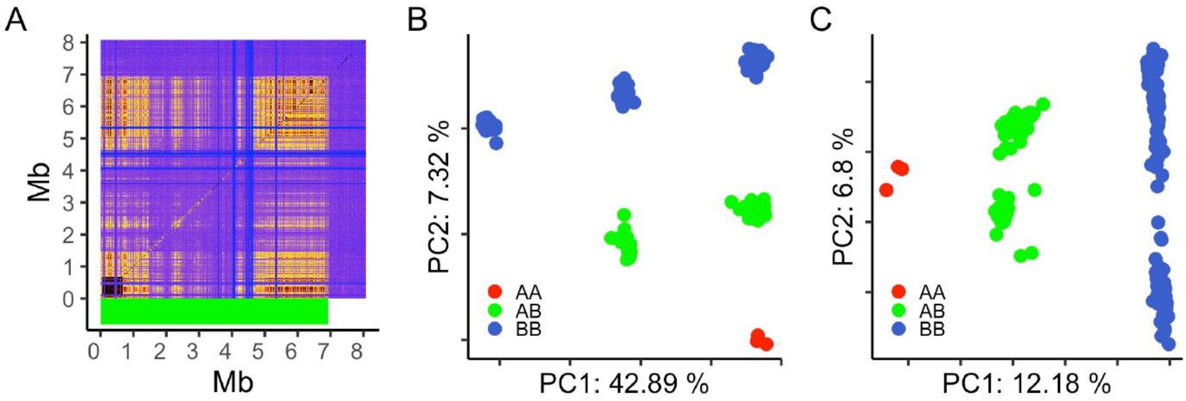

**Inv14.6**

Inv14.6 overlaps with Inv14.1 and Inv14.2 (at the beginning of the contig). Plots show PCA of the region of Inv14.6 overlapping with Inv14.1 (A); PCA of the region of Inv14.6 overlapping with 14.2 (B), $F_{ST}$ between both Inv14.6 haplotypes (C), linkage disequilibrium patterns for IpsContig14 with Inv14.6 position indicated as green bar (D) and population recombination rate for individuals with different genotypes (E). AA, AB, BB - different inversion genotypes; MJ - more frequent haplotype; MN - less frequent haplotype; HET heterozygotes. Genotyping was performed based on PC2 from (A) and PC2-PCA from (B). Both genotyping methods give the same results.

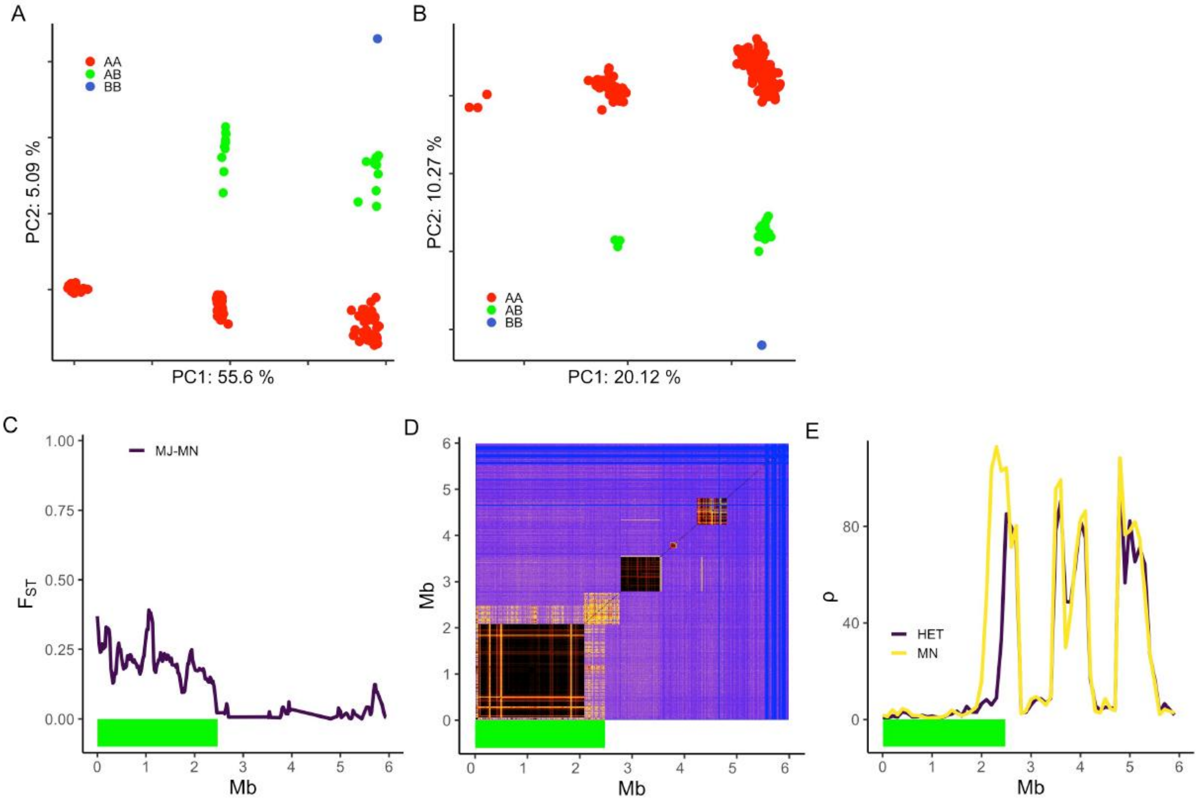

**Inv.16.2**

Inv16.2 overlaps with Inv16.1. It is not visible at the LD plots (C) but additional clustering to three groups is visible at PC2 from PCA analysis of the whole 16.1 inversion (A). Genotyping was done based on PC2 from PCA from the region of 16.1 (A). B – $F_{ST}$ between both inversion haplotypes; C – linkage disequilibrium patterns; D – population recombination rate for individuals with different genotypes. AA, AB, BB – different inversion genotypes; MJ – more frequent haplotype; MN – less frequent haplotype; HET – heterozygotes. Inversion position indicated with green bar.

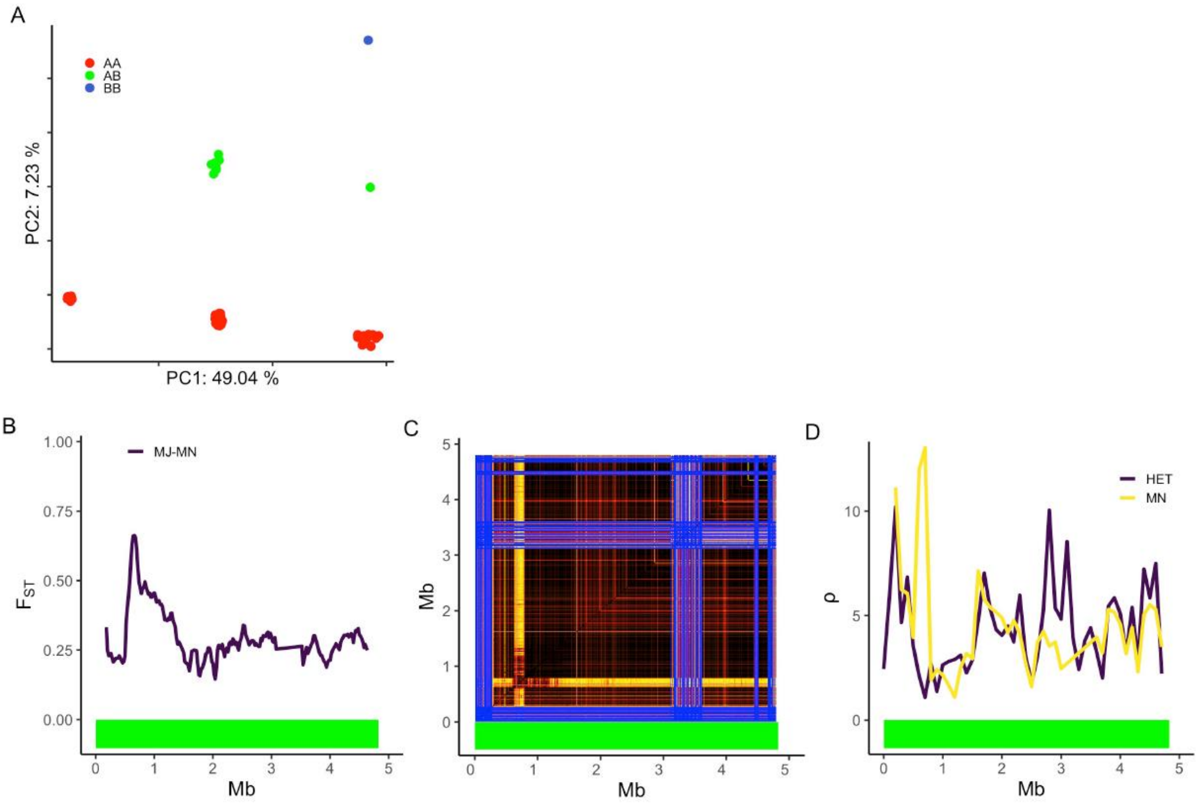

**Inv22.5**

Inv22.5 overlaps with Inv22.1, Inv22.2, Inv22.3 and 22.4. Genotyping was performed based on PC2 from PCAs of the region of 22.5 overlapping with 22.1 (A); 22.2 (B); 22.4 (D) and PC3 from PCA of the region of 22.5 overlapping with 14.3 (C; in this case PC2 mainly showed patterns of double crossover, see above). All genotyping methods give the same results. E - $F_{ST}$ between the two inversion haplotypes; F - Linkage disequilibrium patterns; G - Population recombination rate for individuals with different genotypes. AA, AB, BB - different inversion genotypes; MJ - more frequent haplotype; MN - less frequent haplotype; HET heterozygotes. The position of the inversion is indicated by a green bar.

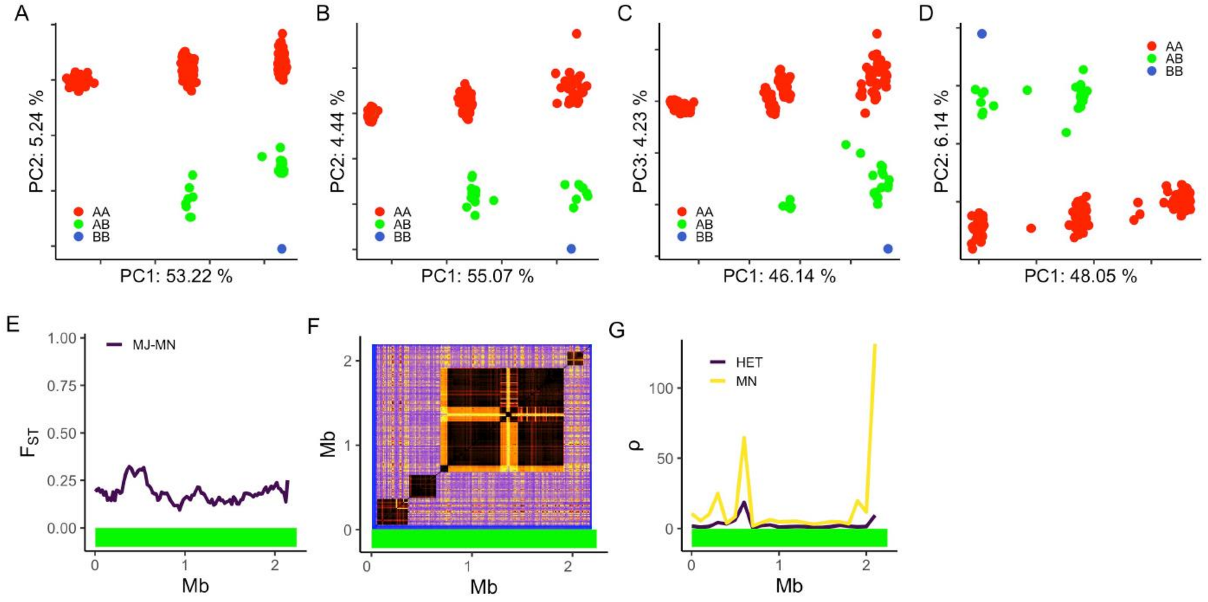

**Inv 23.2**

Inv23.2 overlaps with Inv23.1. It is not visible in the LD plots (C), but additional clustering into three groups is visible in PC2 from PCA analysis of the entire 23.1 inversion (A). Genotyping was performed based on PC2 from PCA of the 23.1 region (A). B - $F_{ST}$ between both inversion haplotypes; C - Linkage disequilibrium patterns; D - Population recombination rate for individuals with different genotypes. AA, AB, BB - different inversion genotypes; MJ - more frequent haplotype; MN - less frequent haplotype; HET heterozygotes. The position of the inversion is indicated by a green bar.

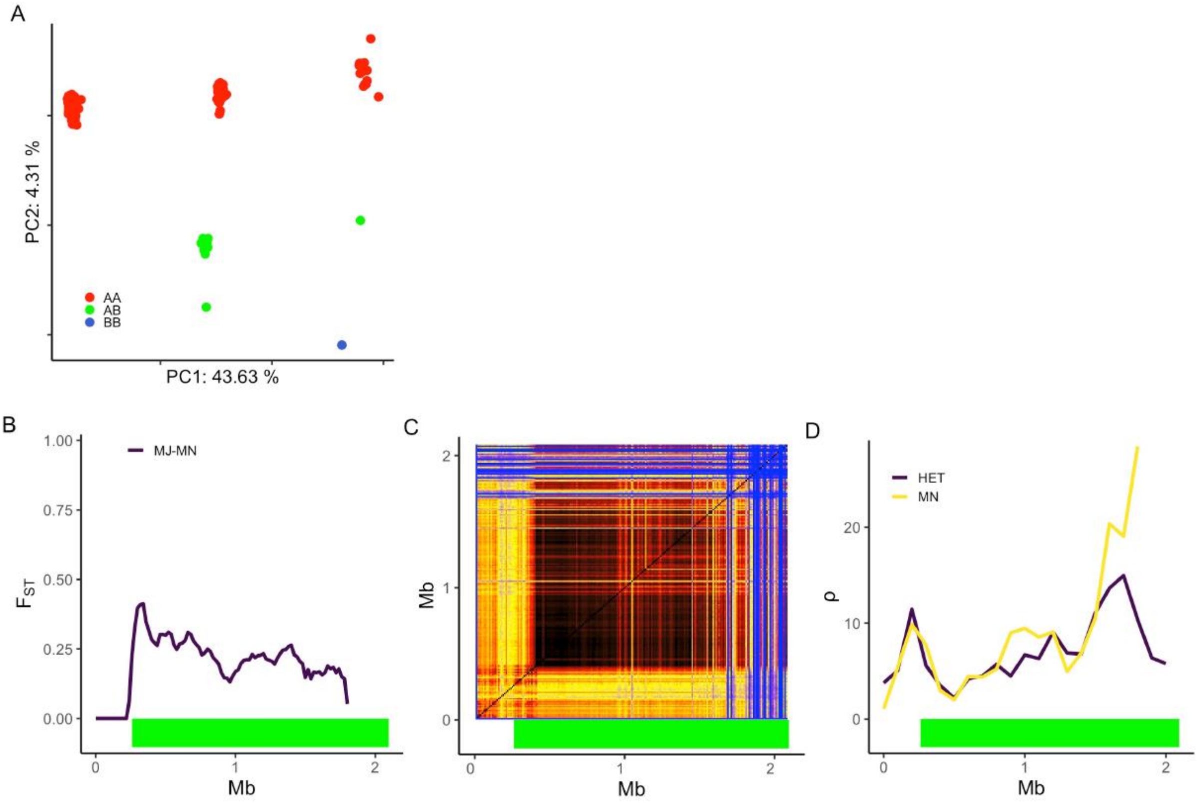

**Figure S4** Linkage disequilibrium (LD) between inversions in the *Ips typographus* genome. Non-blue off diagonal squares for Inv16.1 and Inv23.1 indicate parts of the same inversion in different contigs. LD was calculated between inversions that could be genotyped in all studied populations. The color bar represents LD from 0 to 1.

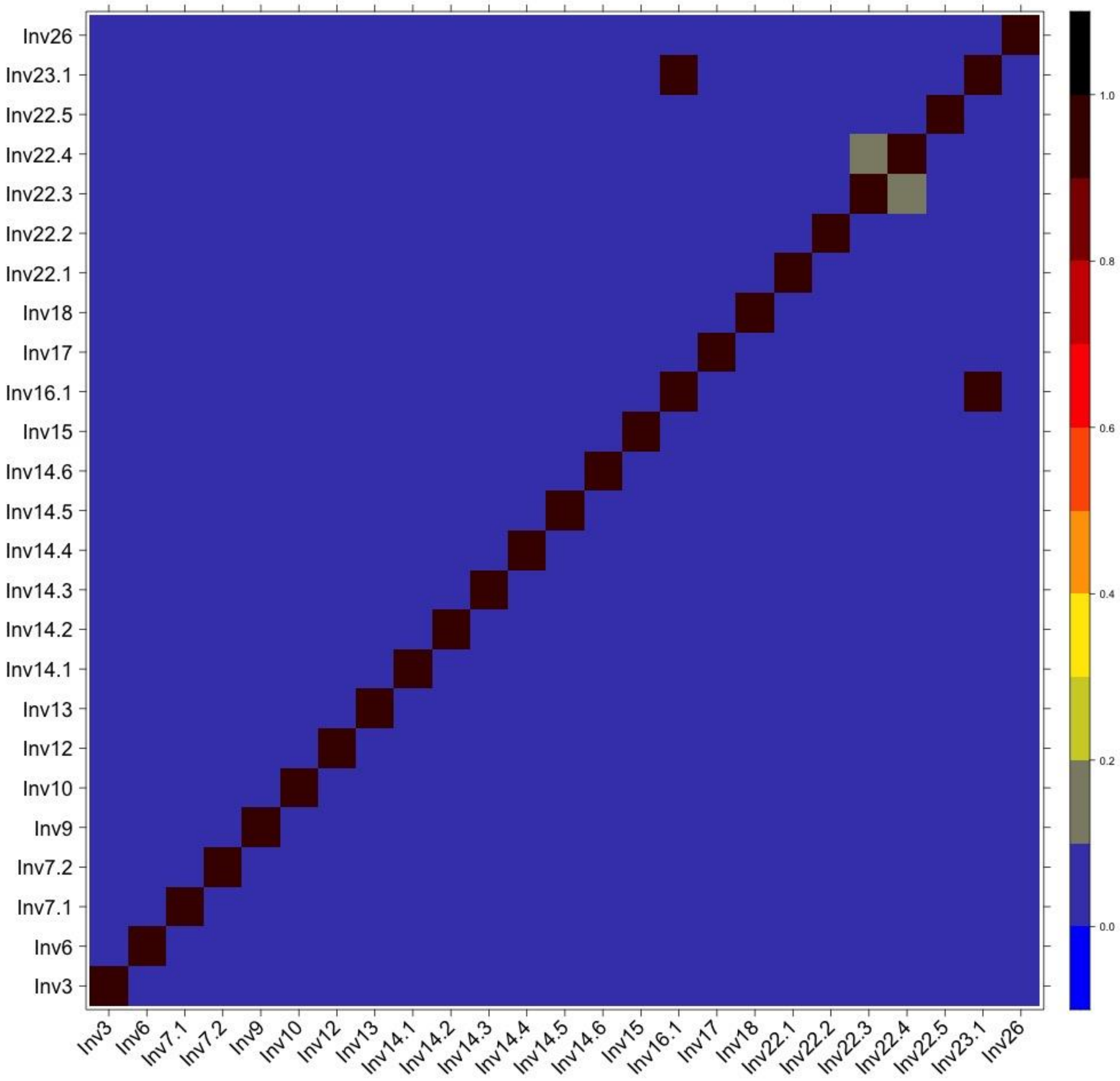

**Figure S5** Correlation between inversion age and the major (MJ) inversion haplotype frequency.

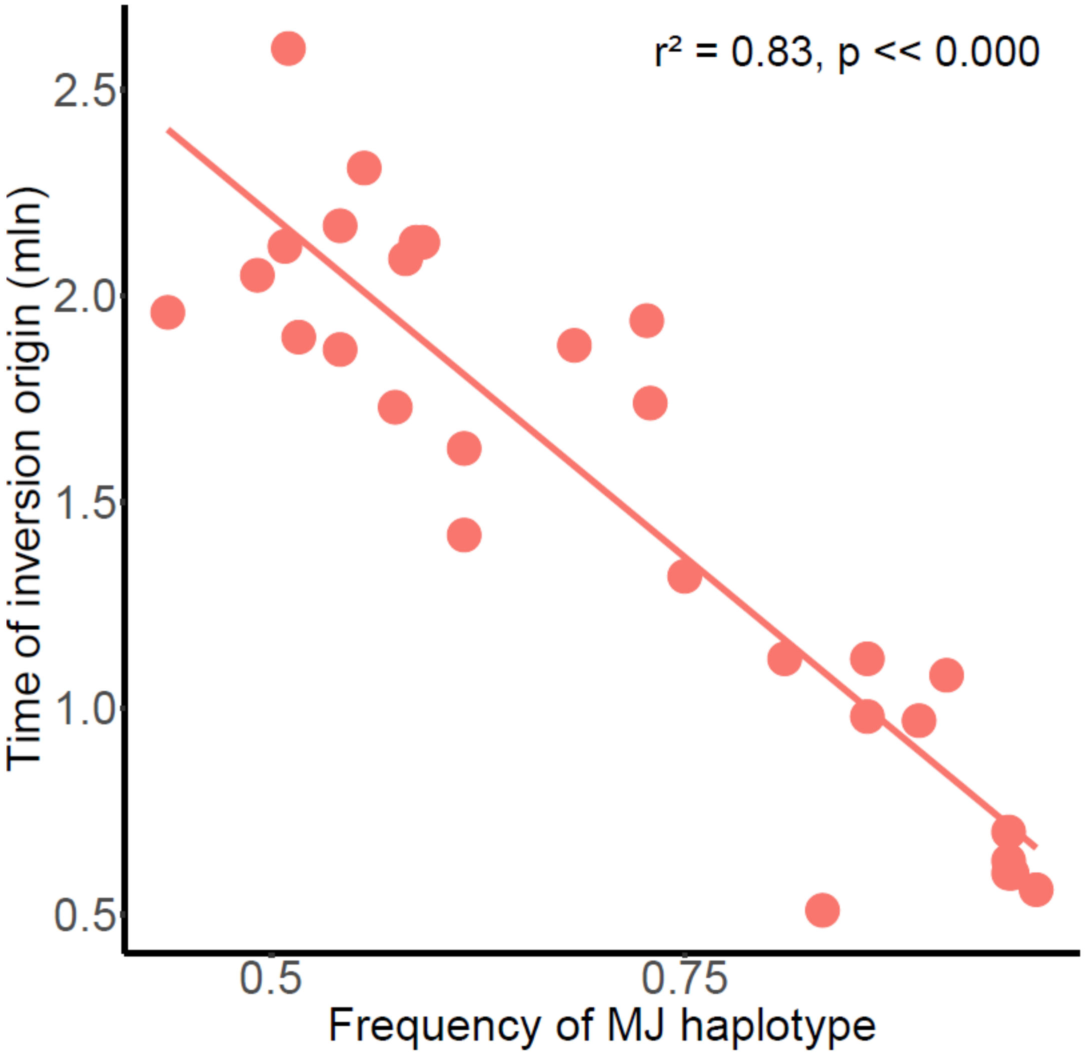

**Figure S6** Inversion PCA analysis for northern (A) and southern (B) groups separately. Individuals colored by population.

#### Inv2

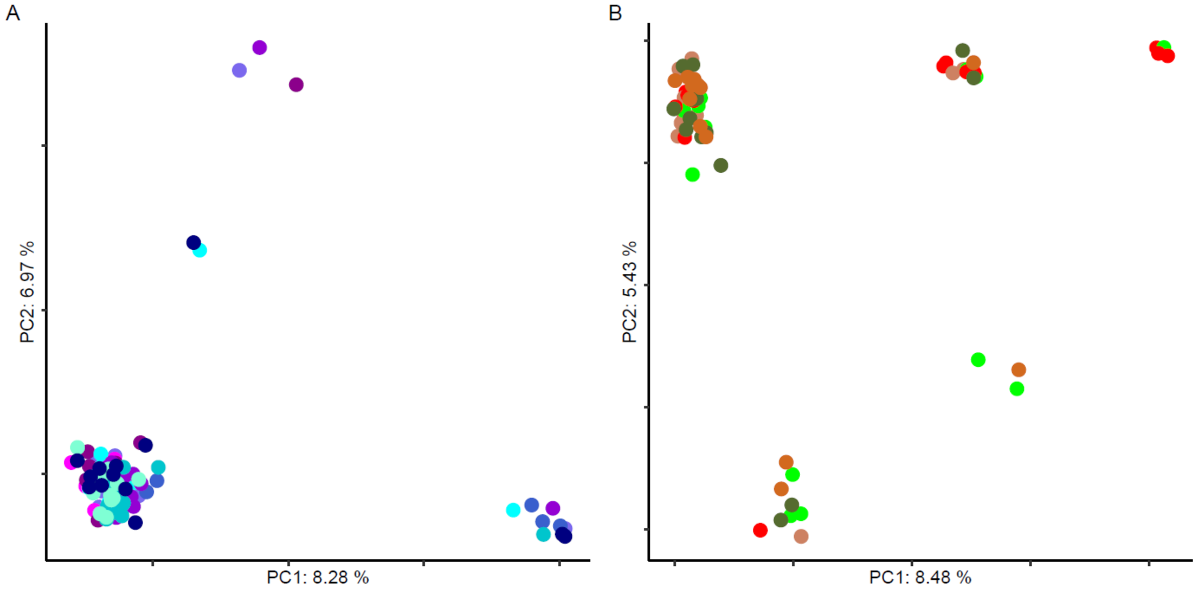


#### Inv3

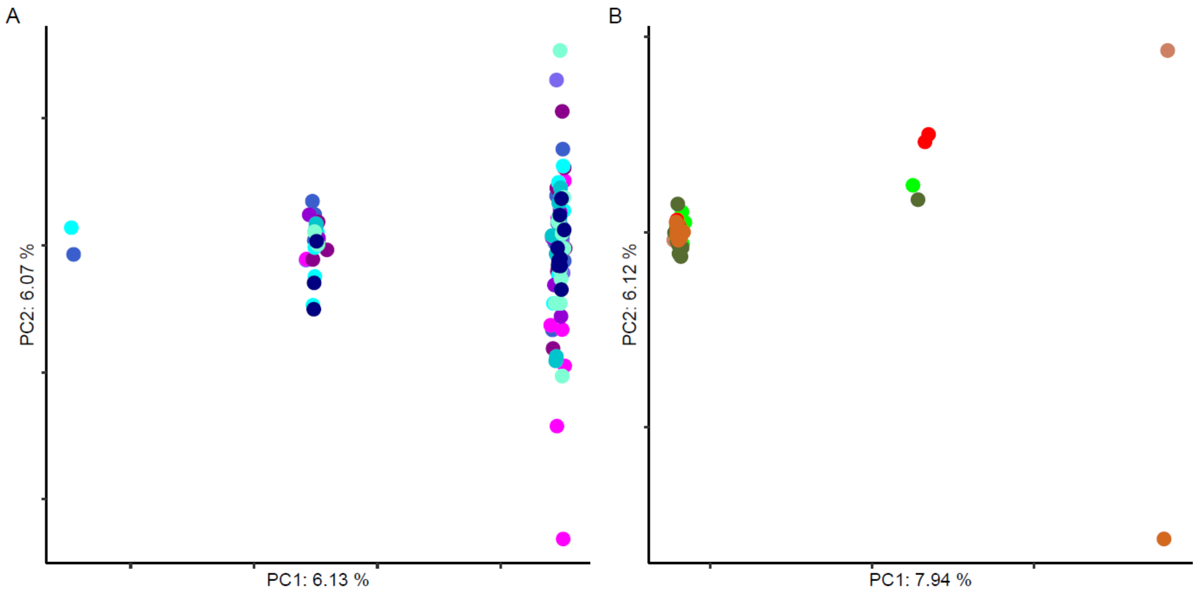

### Inv5

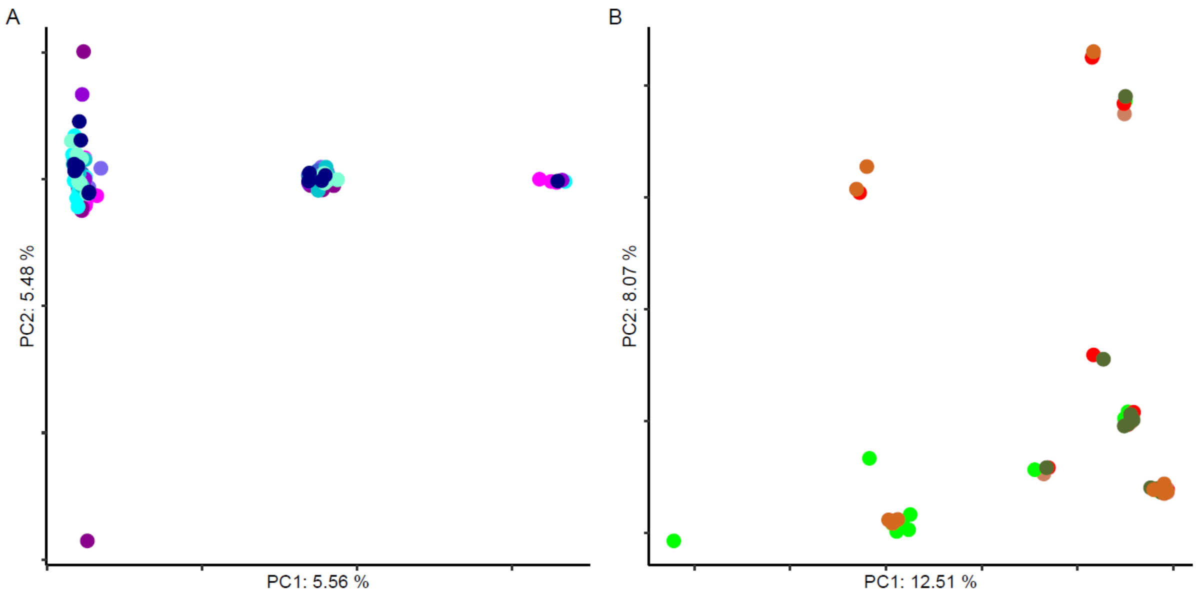
A
PC2: 5.48 %
PC1: 5.56 %
B
PC2: 8.07 %
PC1: 12.51 %

### Inv6

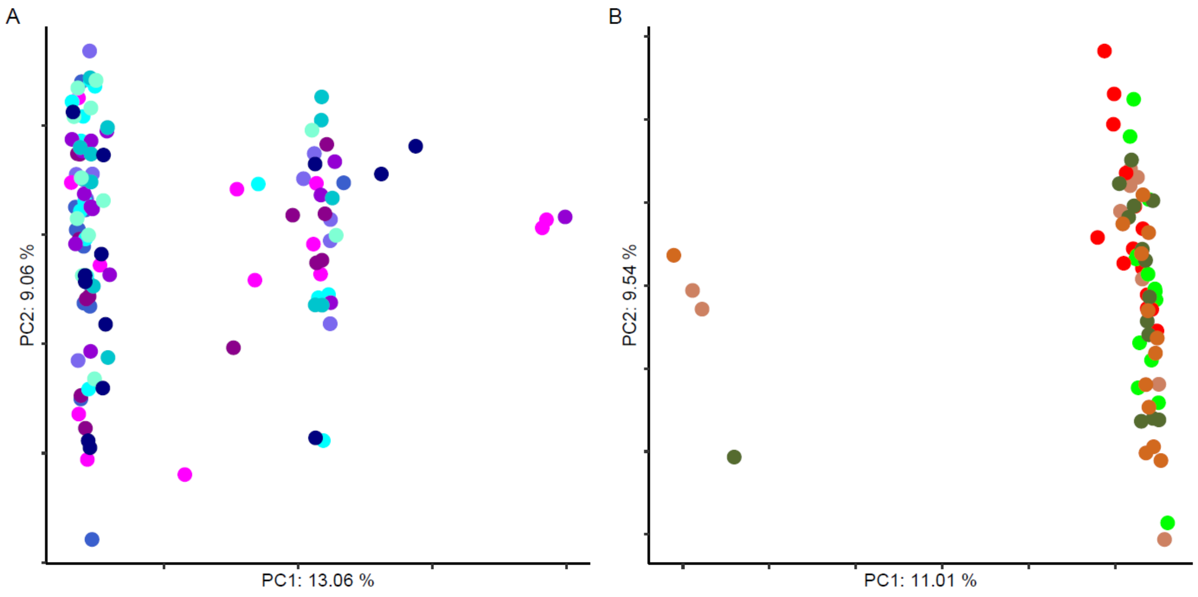
A
PC2: 9.06 %
PC1: 13.06 %
B
PC2: 9.54 %
PC1: 11.01 %

### Inv7.1

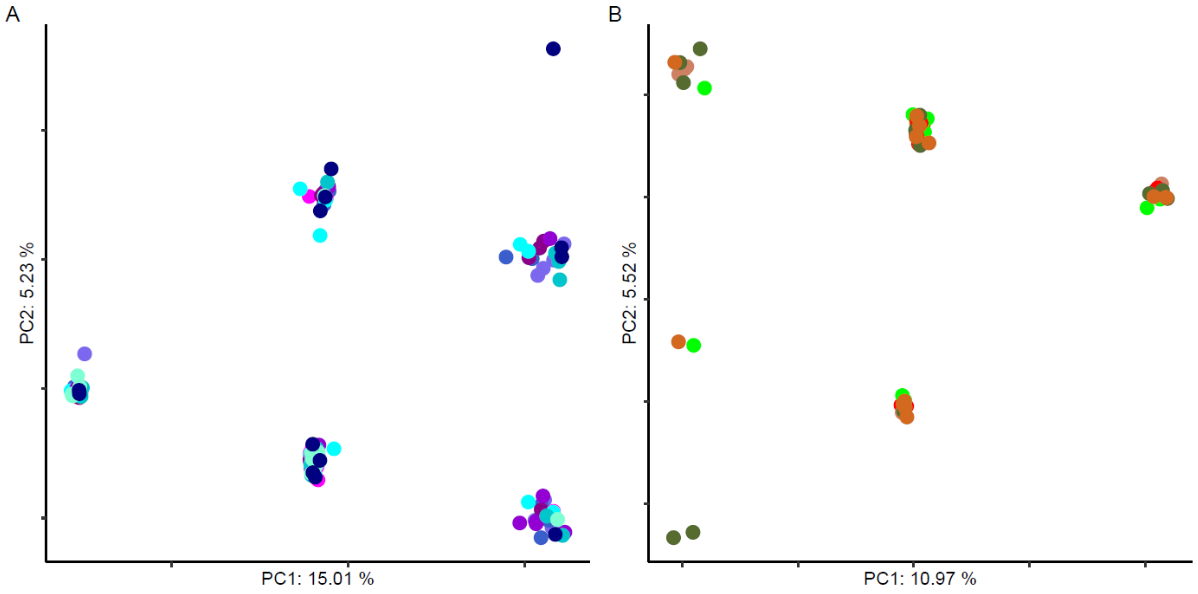
A
PC2: 5.23 %
PC1: 15.01 %
B
PC2: 5.52 %
PC1: 10.97 %

### Inv7.2

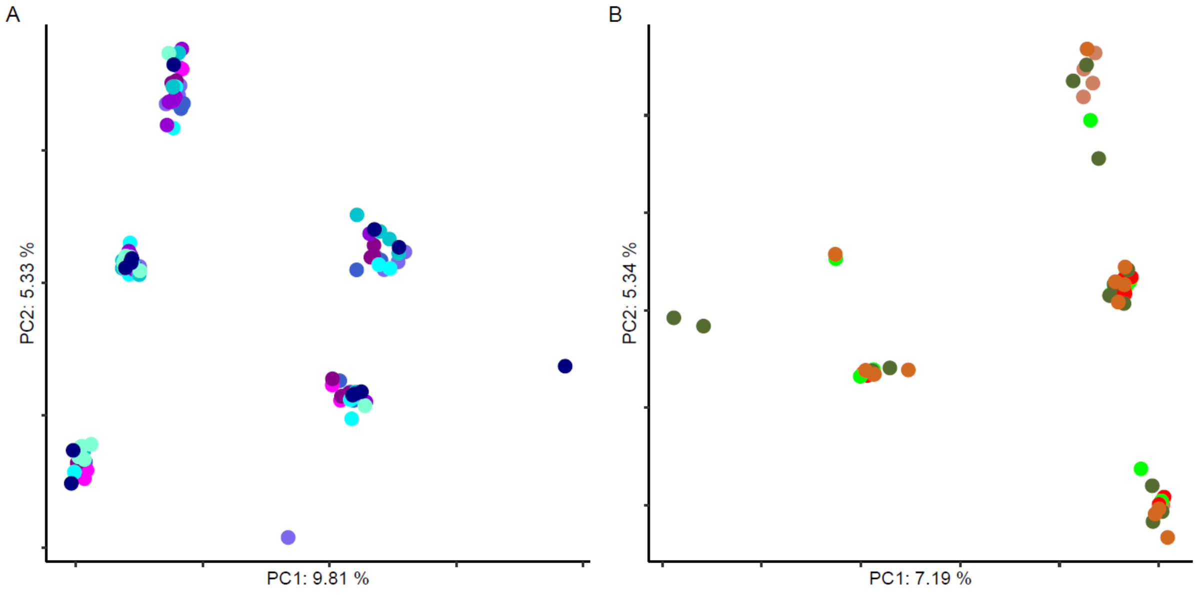
A
PC2: 5.33 %
PC1: 9.81 %
B
PC2: 5.34 %
PC1: 7.19 %

### Inv9

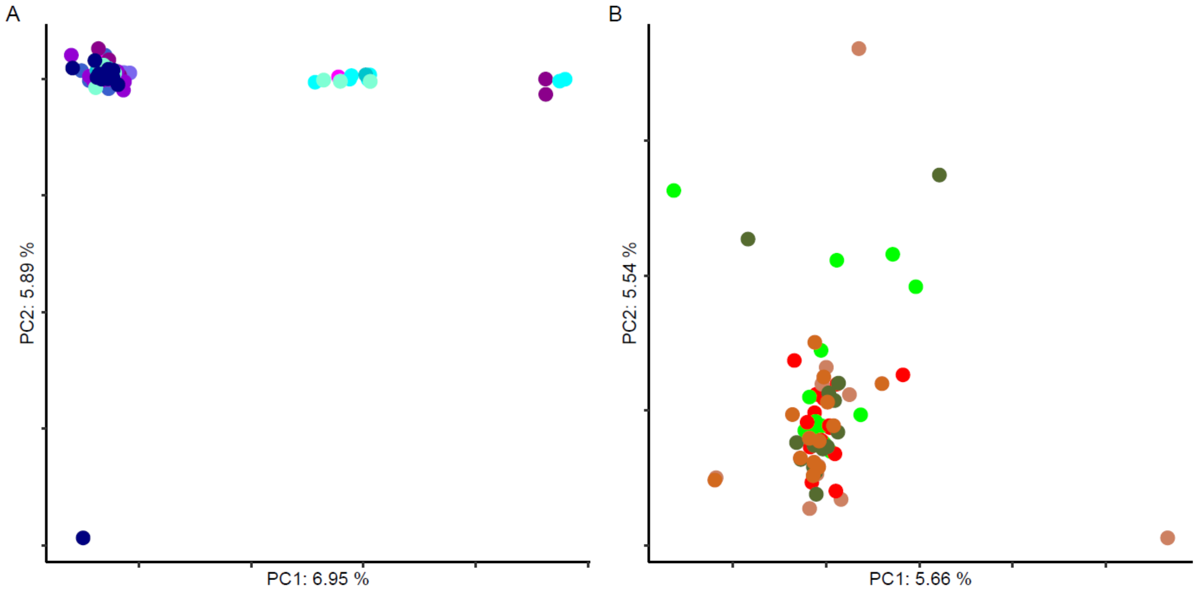
A
PC2: 5.89 %
PC1: 6.95 %
B
PC2: 5.54 %
PC1: 5.66 %

### Inv10

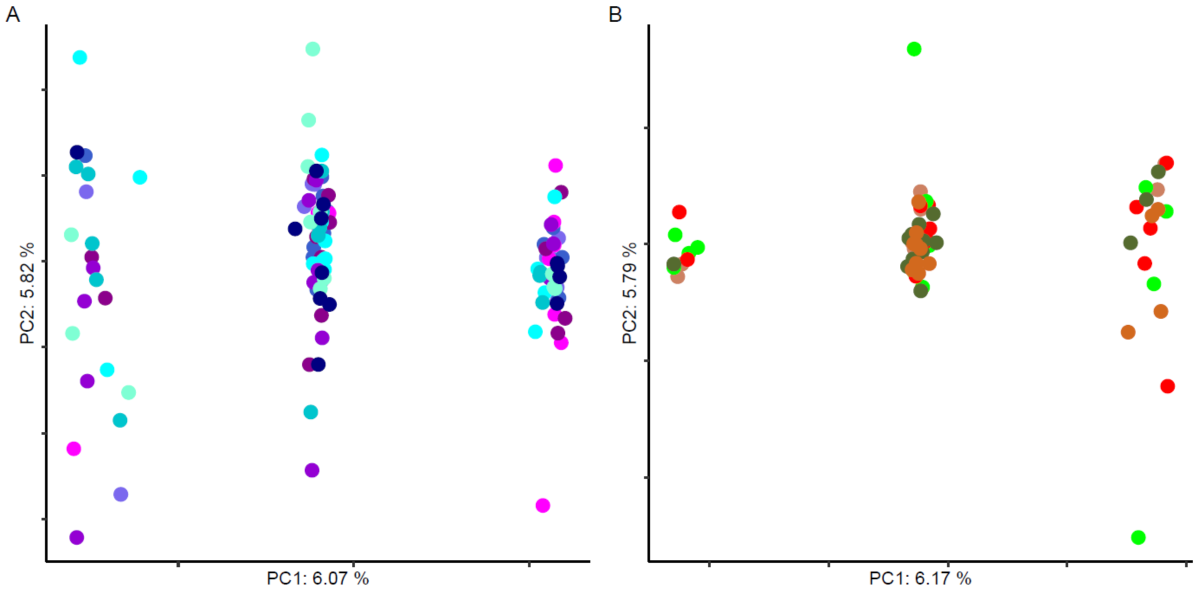
A
PC2: 5.82 %
PC1: 6.07 %
B
PC2: 5.79 %
PC1: 6.17 %

### Inv12

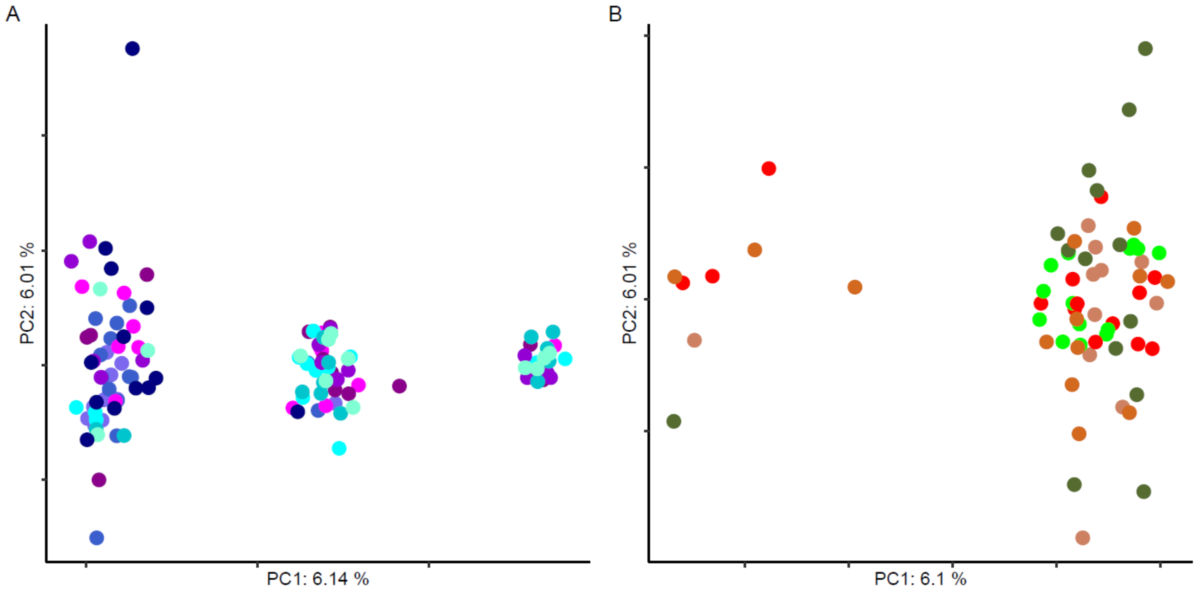
A
PC2: 6.01 %
PC1: 6.14 %
B
PC2: 6.01 %
PC1: 6.1 %

### Inv13

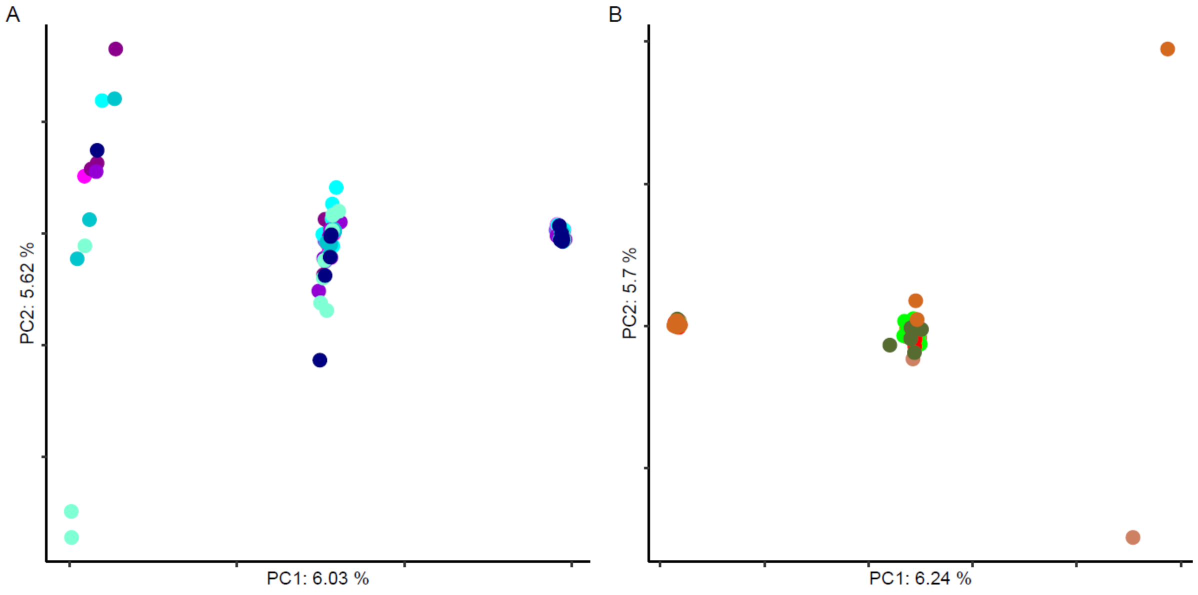
A
PC2: 5.62 %
PC1: 6.03 %
B
PC2: 5.7 %
PC1: 6.24 %

### Inv14.1

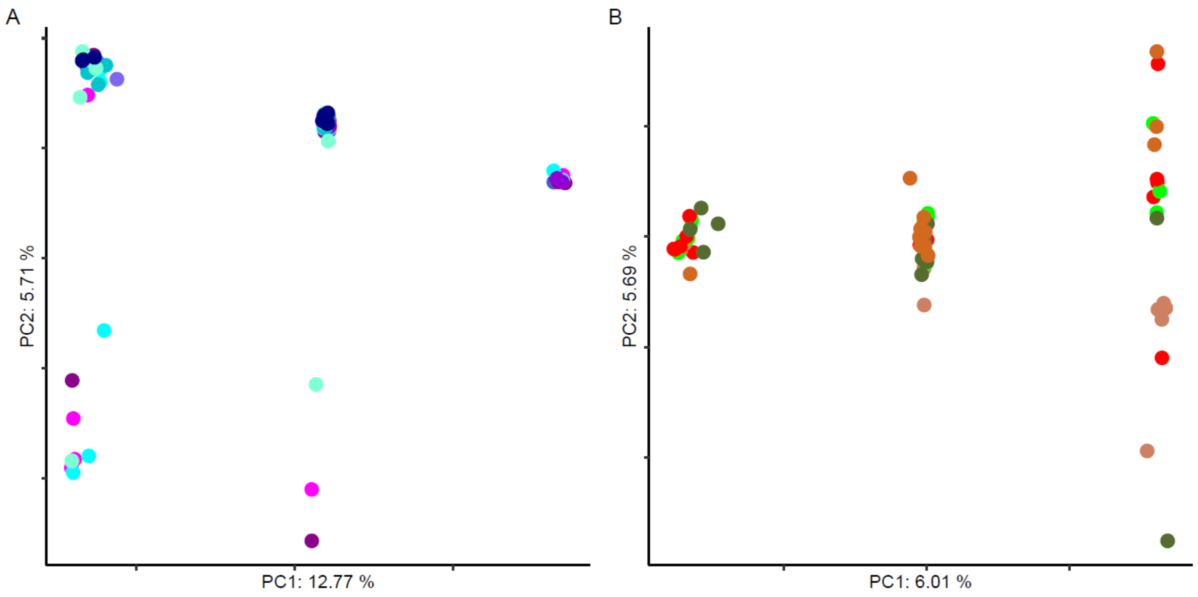


### Inv14.2

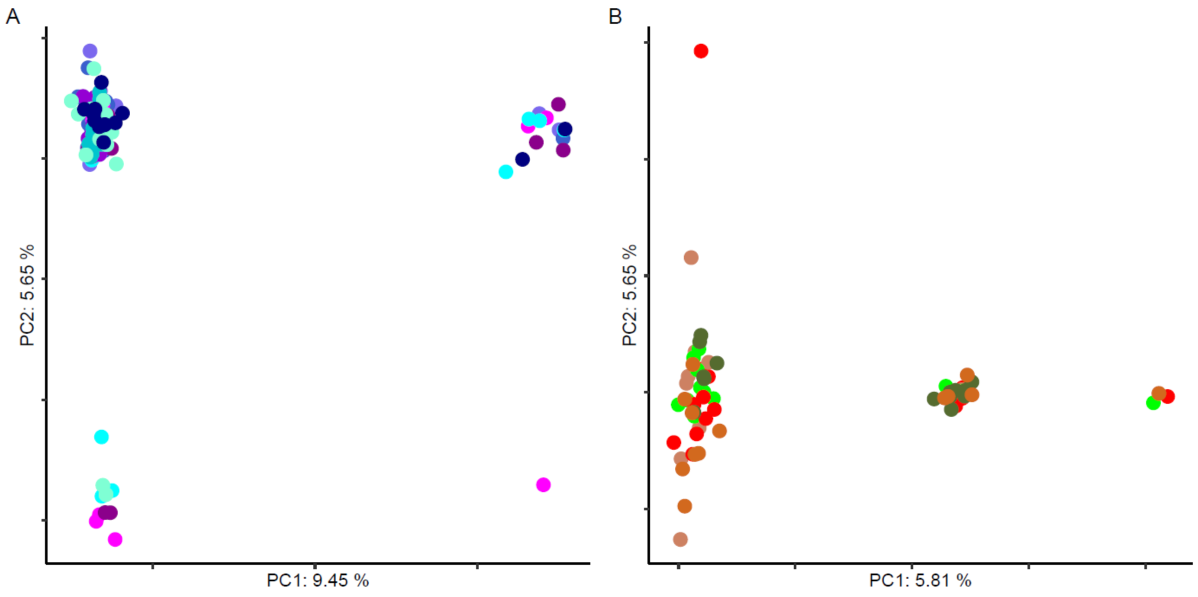

**Inv14.3**

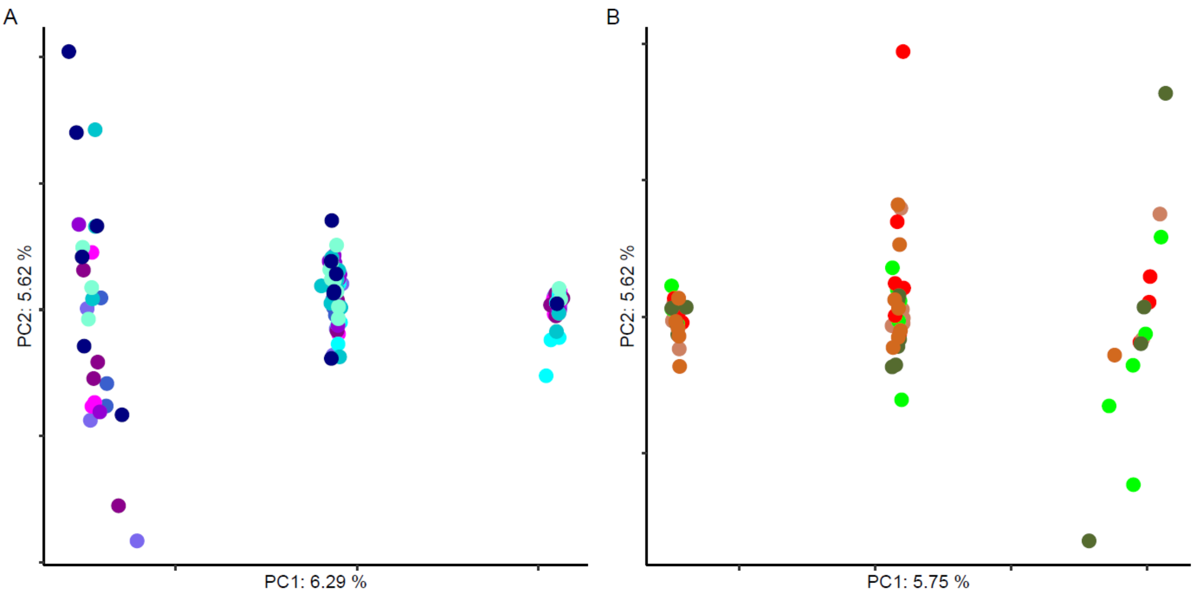


**Inv14.4**

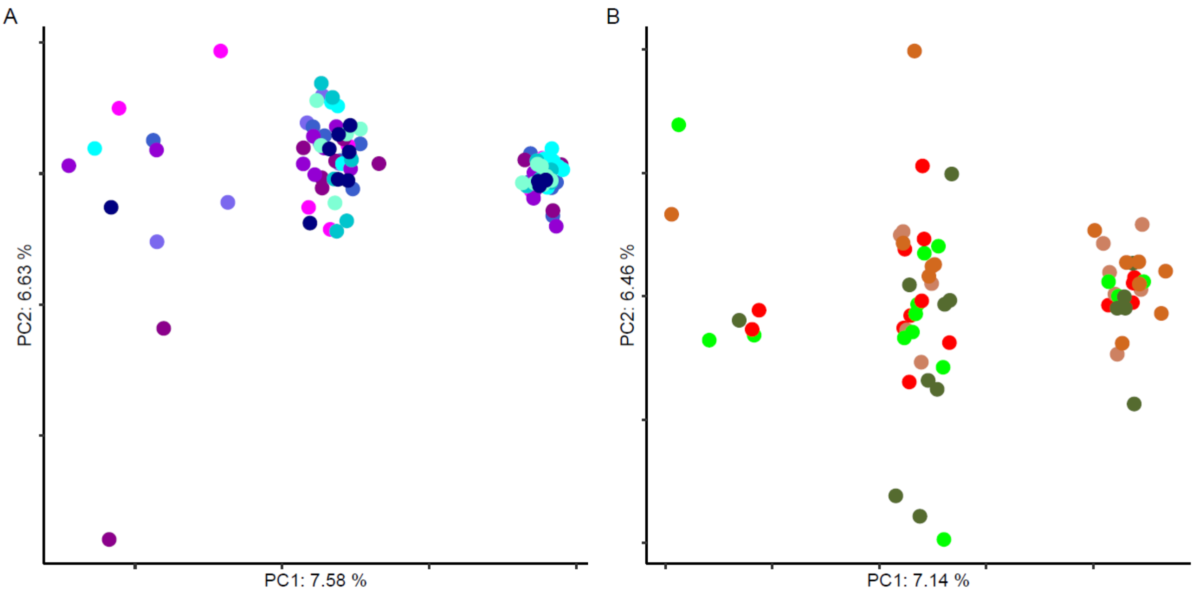

**Inv14.5**

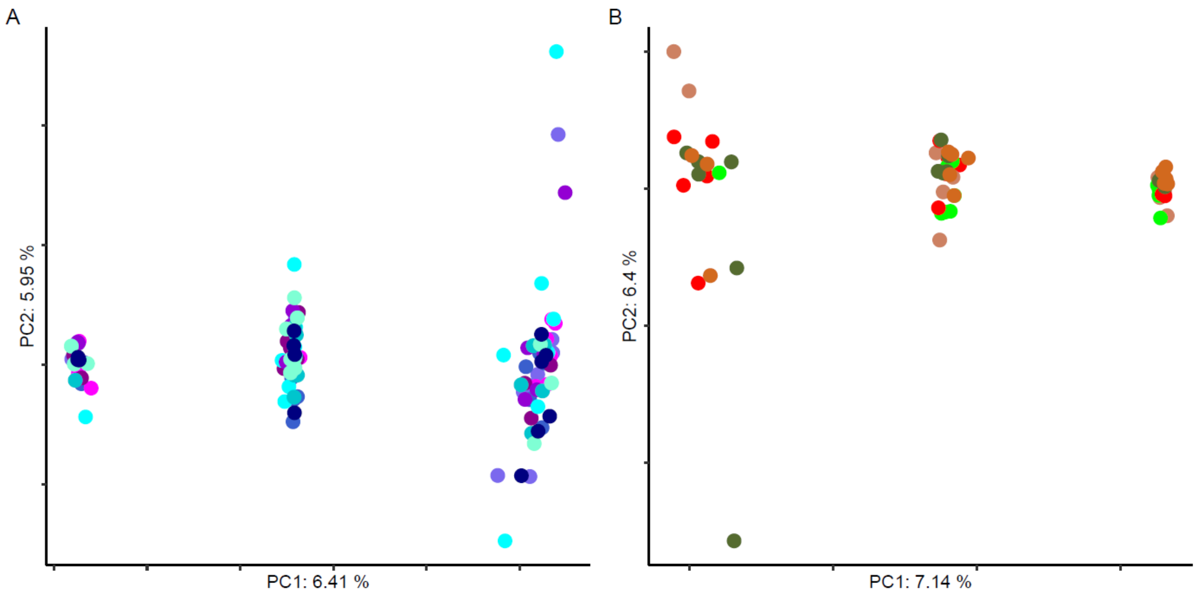


**Inv15**

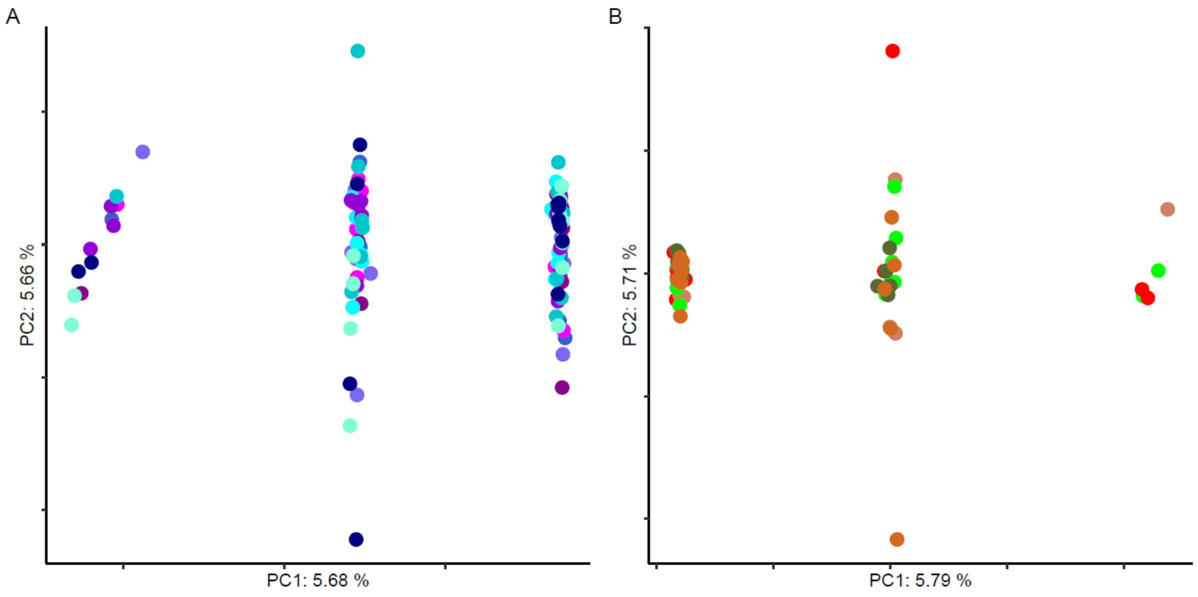

**Inv16.1**

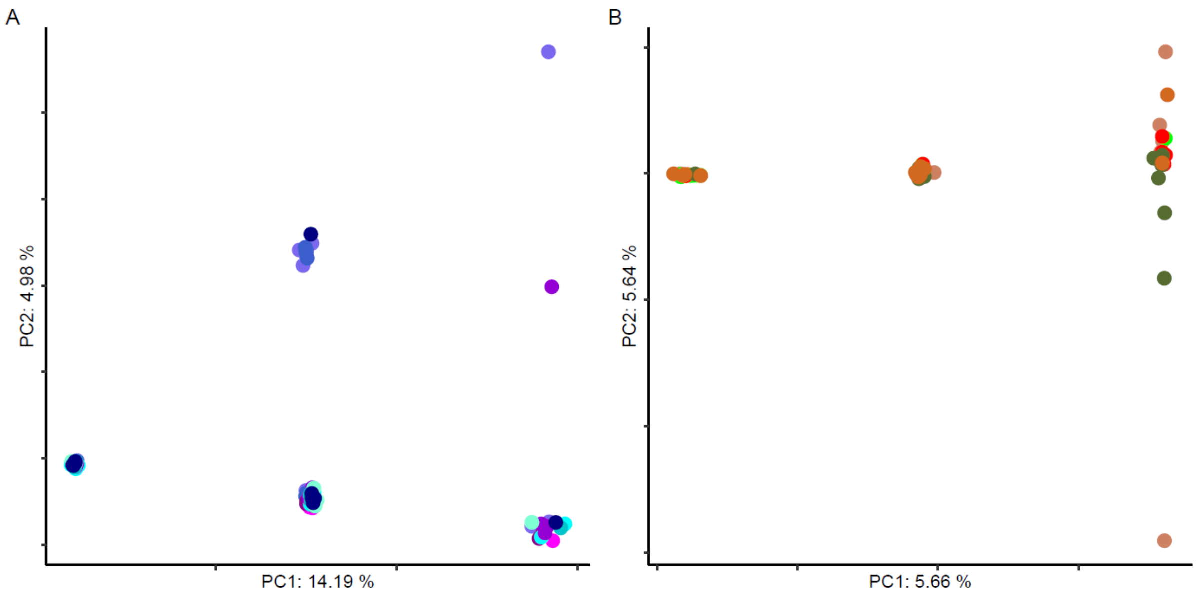


**Inv17**

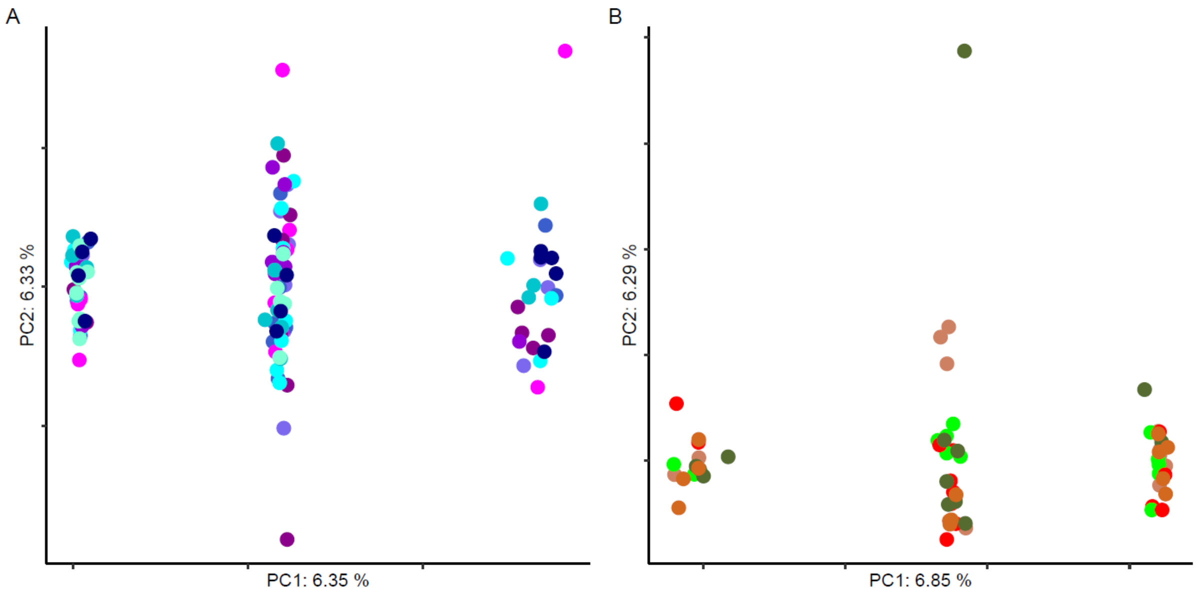

### Inv18

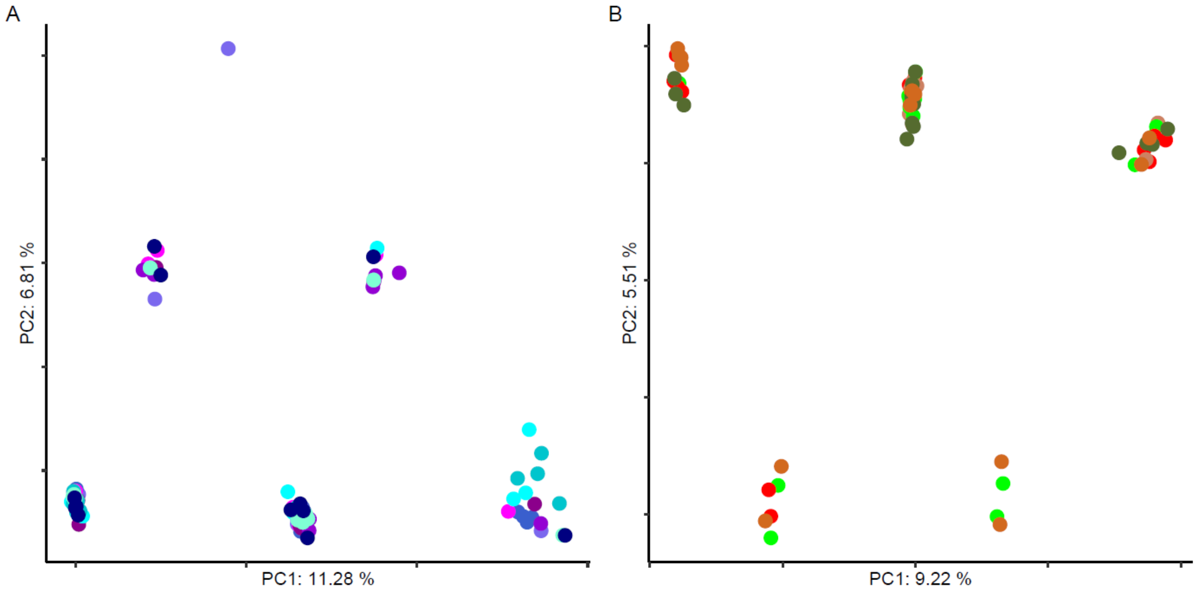


### Inv22.1

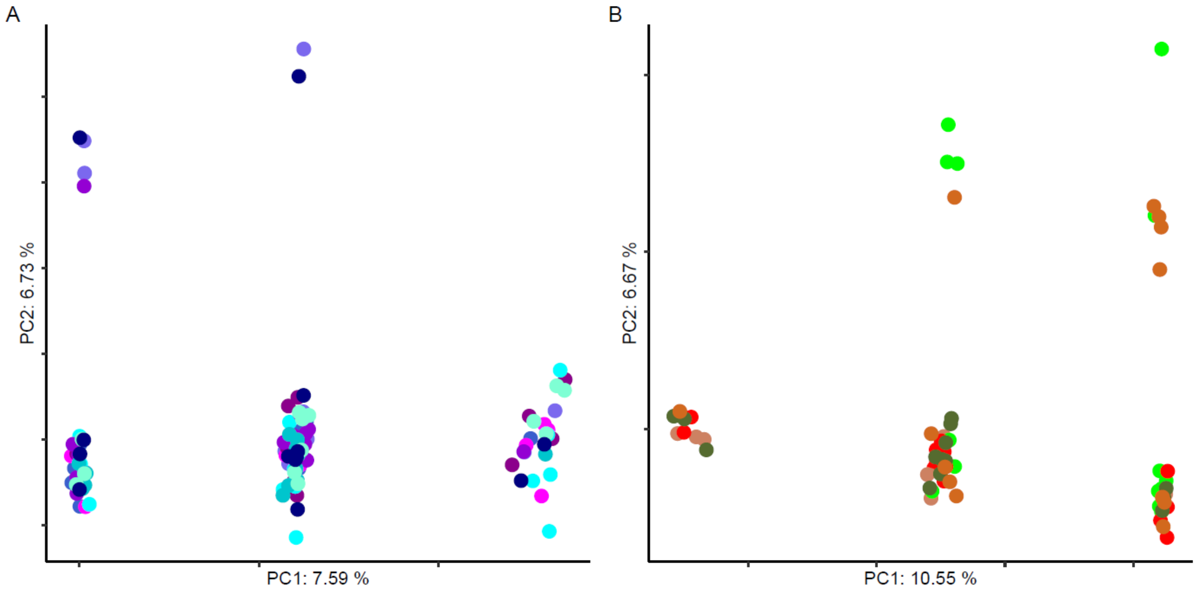

**Inv22.2**

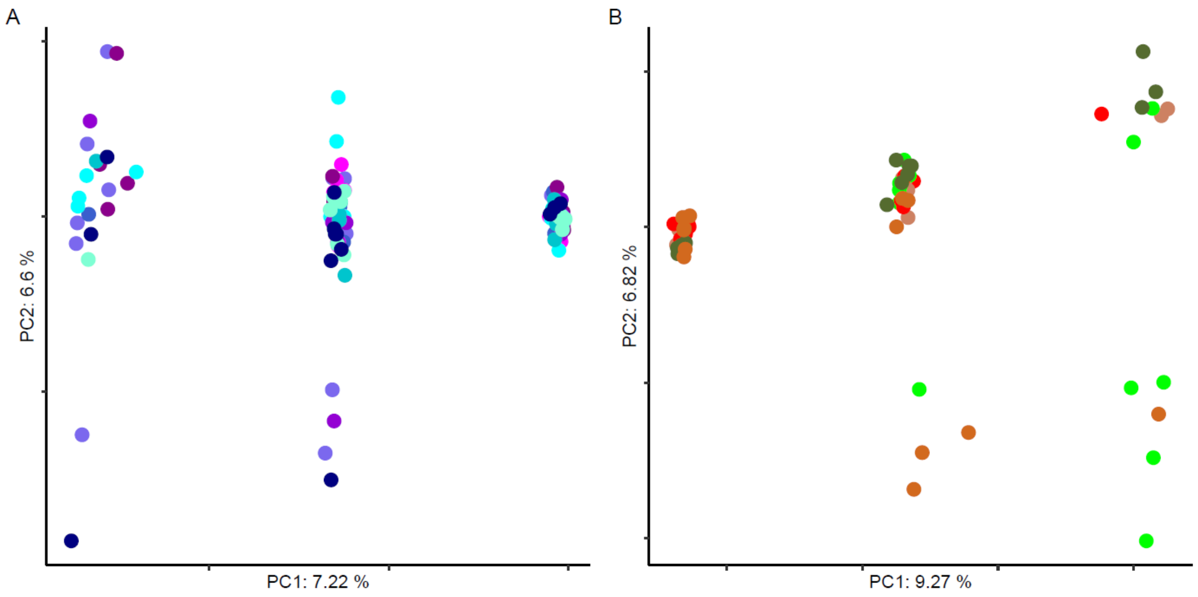


**Inv22.3**

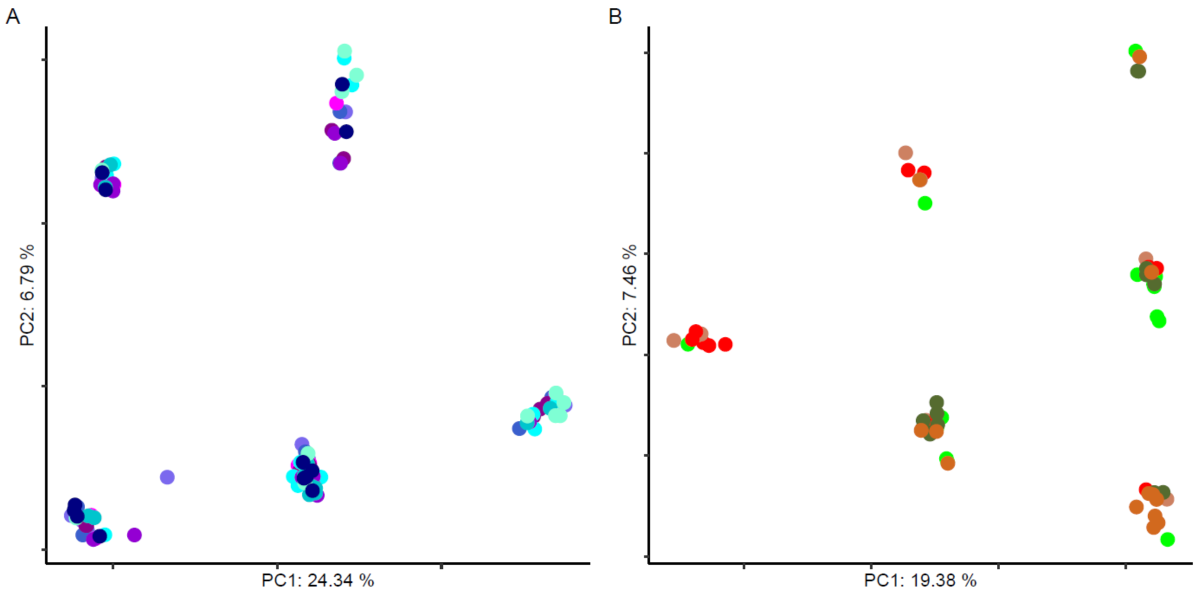

**Inv22.4**

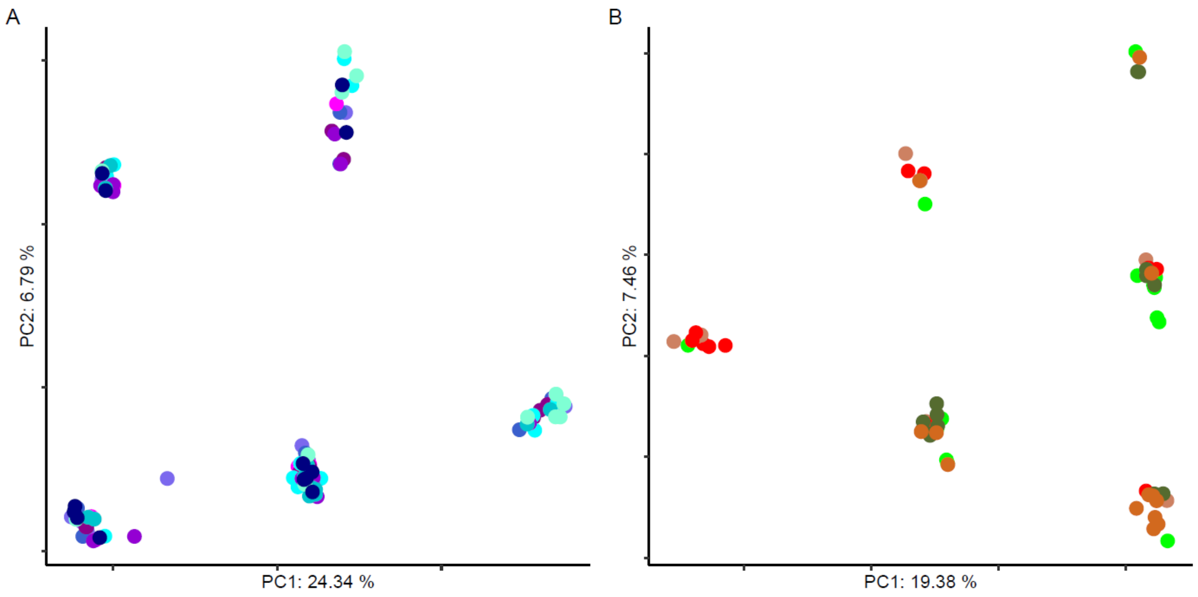


**Inv23.1**

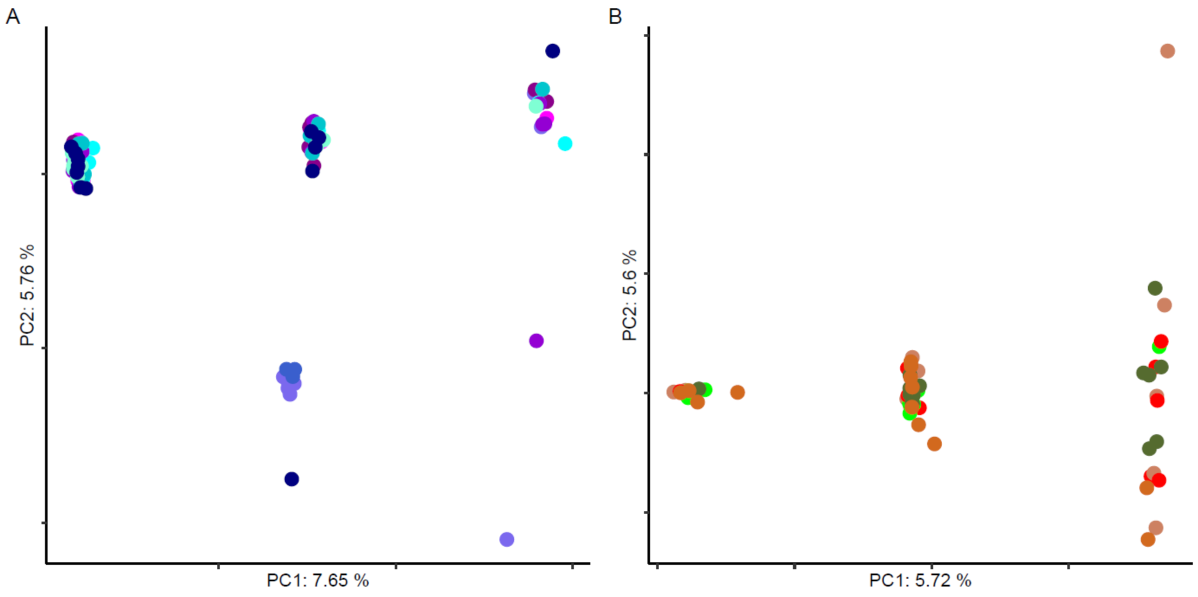

**Inv26**

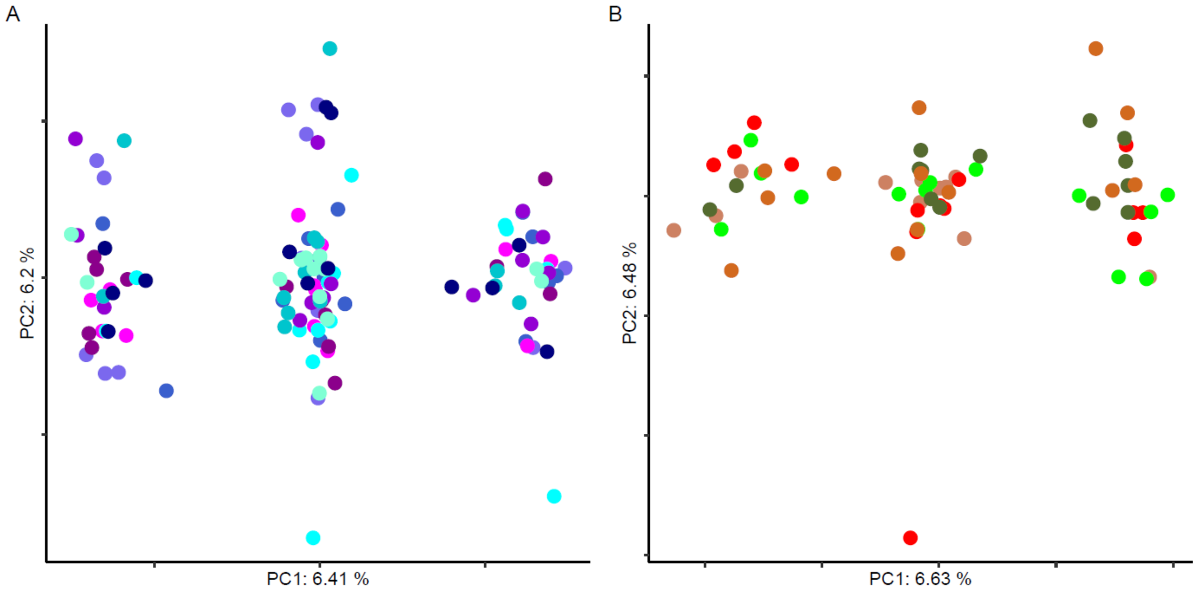

**Figure S7** Nucleotide diversity, Tajima's D and $d_{xy}$ calculated in 100kb windows along 36 contigs from the *Ips typographus* genome. Nucleotide diversity and Tajima's D were calculated per populations: northern, southern and Polish populations are shown in different blue, red/green and yellow colors, respectively. $D_{xy}$ was calculated between southern and northern groups.

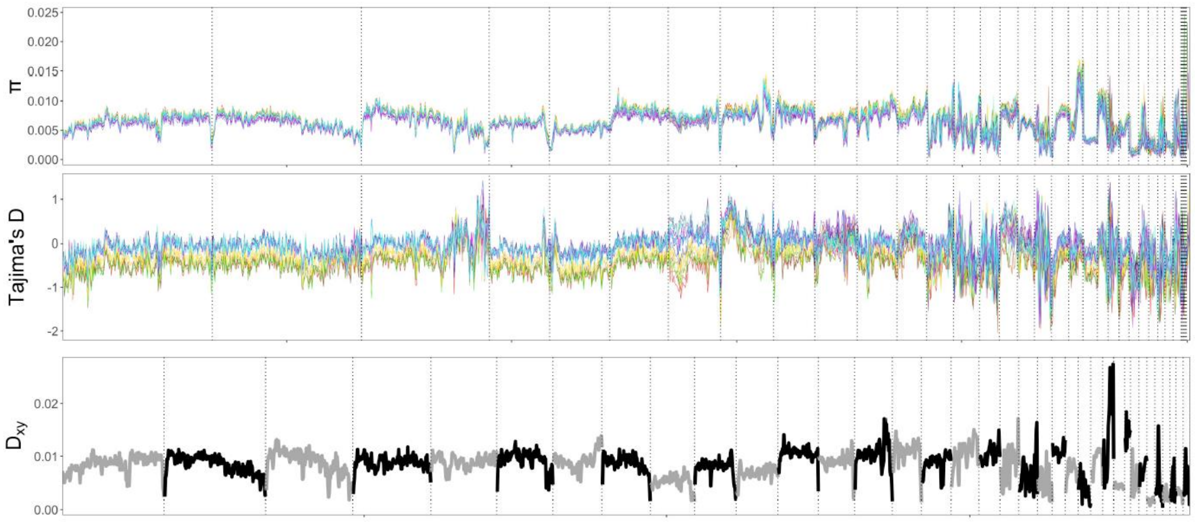

**Figure S8** Correlation between nucleotide diversity (per *Ips typographus* population) and the latitude where the population was collected.

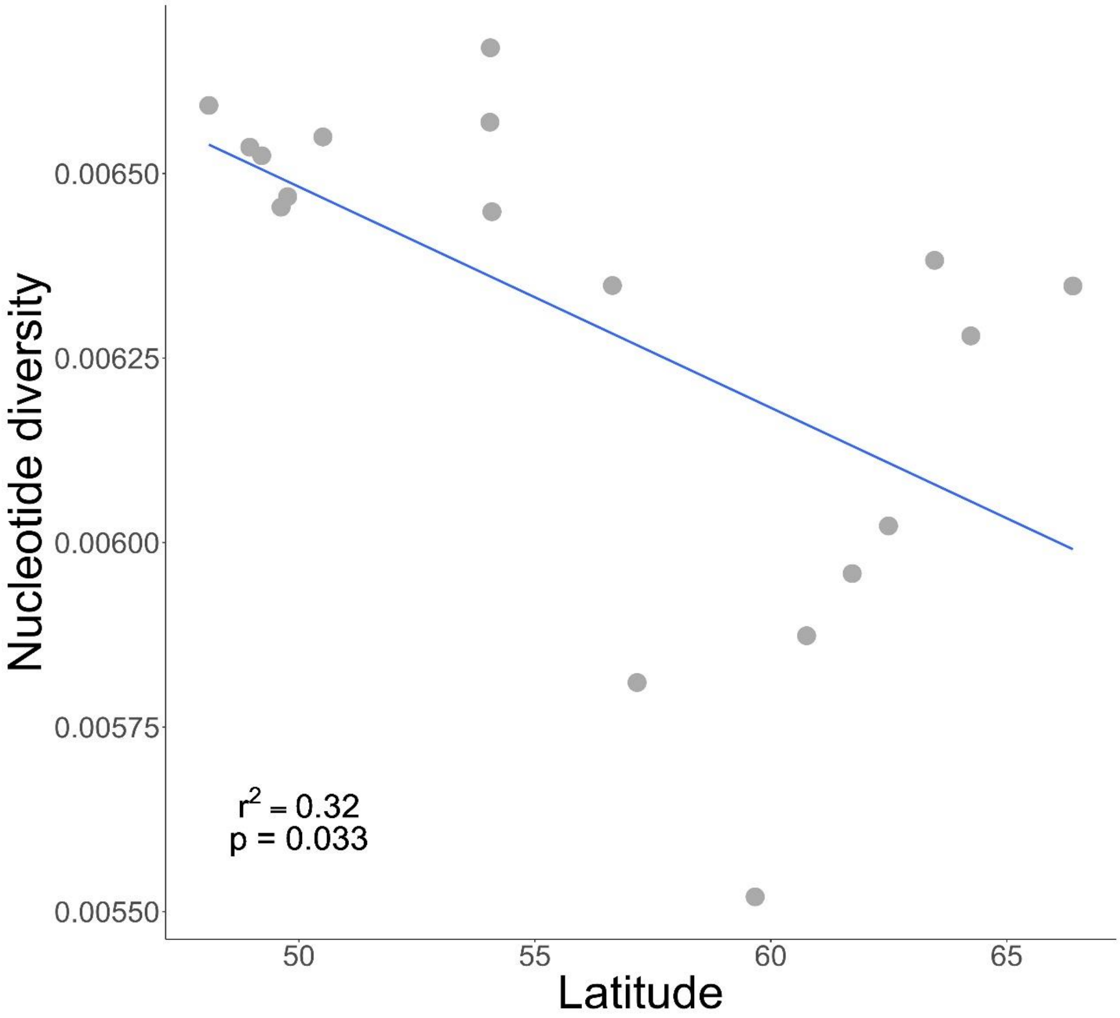

**Figure S9** PC1 versus PC2 for inversion regions based on the dataset that included *Ips typographus* individuals with known diapause phenotype. Only inversions that could be genotyped in all studied populations were included. Purple dots represent individuals with obligate diapause and orange dots individuals with facultative diapause. Grey dots represent *I. typographus* individuals of unknown diapause phenotype.

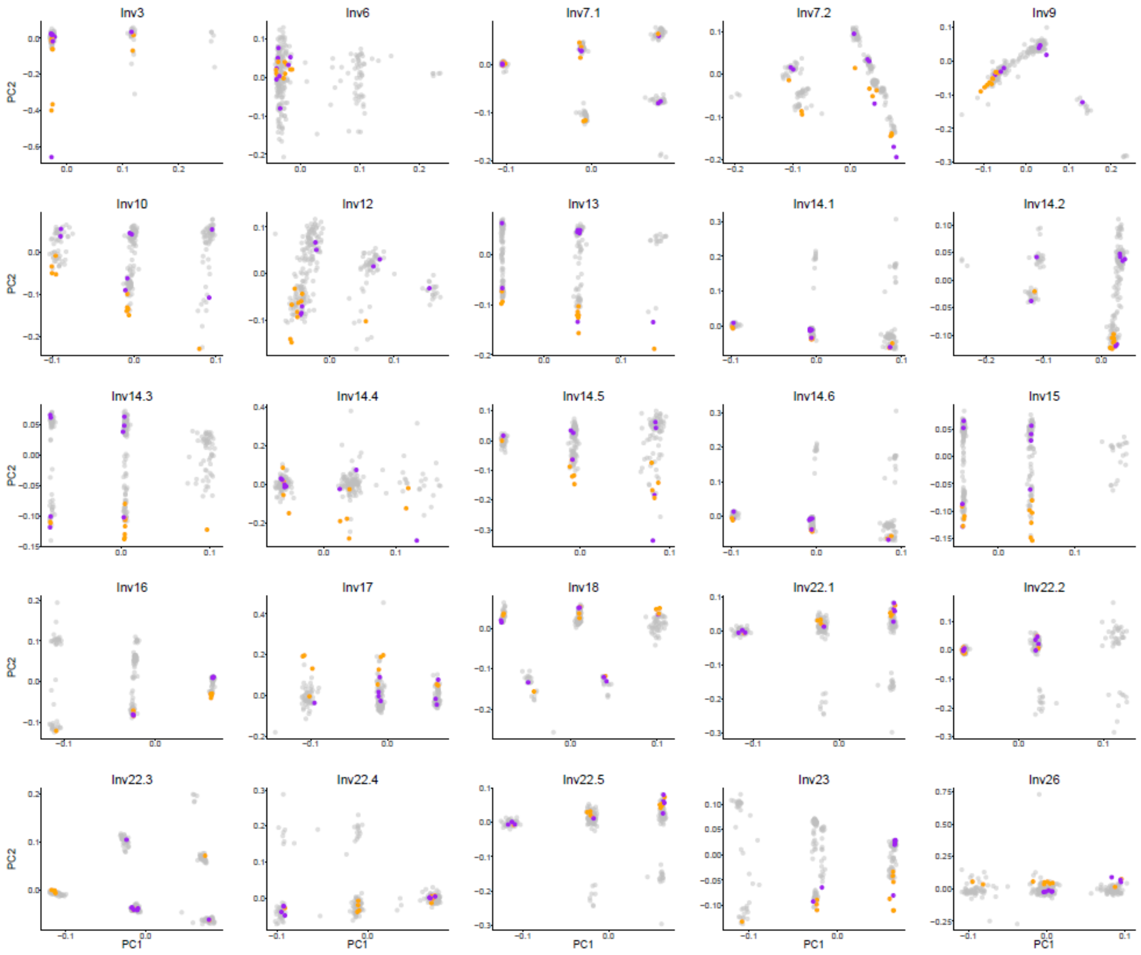

**Figure S10** Genome-wide genetic differentiation ($F_{ST}$) along 36 contigs from the *Ips typographus* genome. Contigs are ordered by length and contigs boundaries are indicated with vertical lines. Red line shows $F_{ST}$ between individuals with facultative and obligate diapause and blue line shows $F_{ST}$ between southern and northern beetle populations.

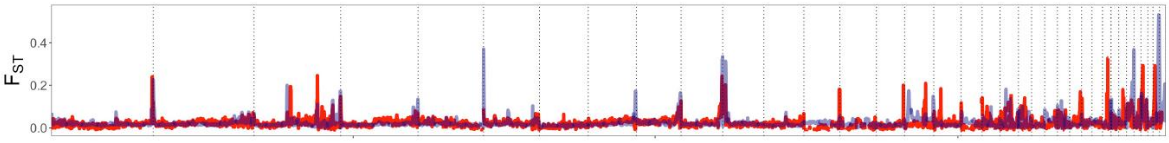

**Figure S11** Correlation between inversion haplotype frequency and longitude for inversions detected in *Ips typographus* populations collected throughout Europe. Inversions harboring odorant receptors are indicated in green. None of the correlations were significant after correction for multiple testing.

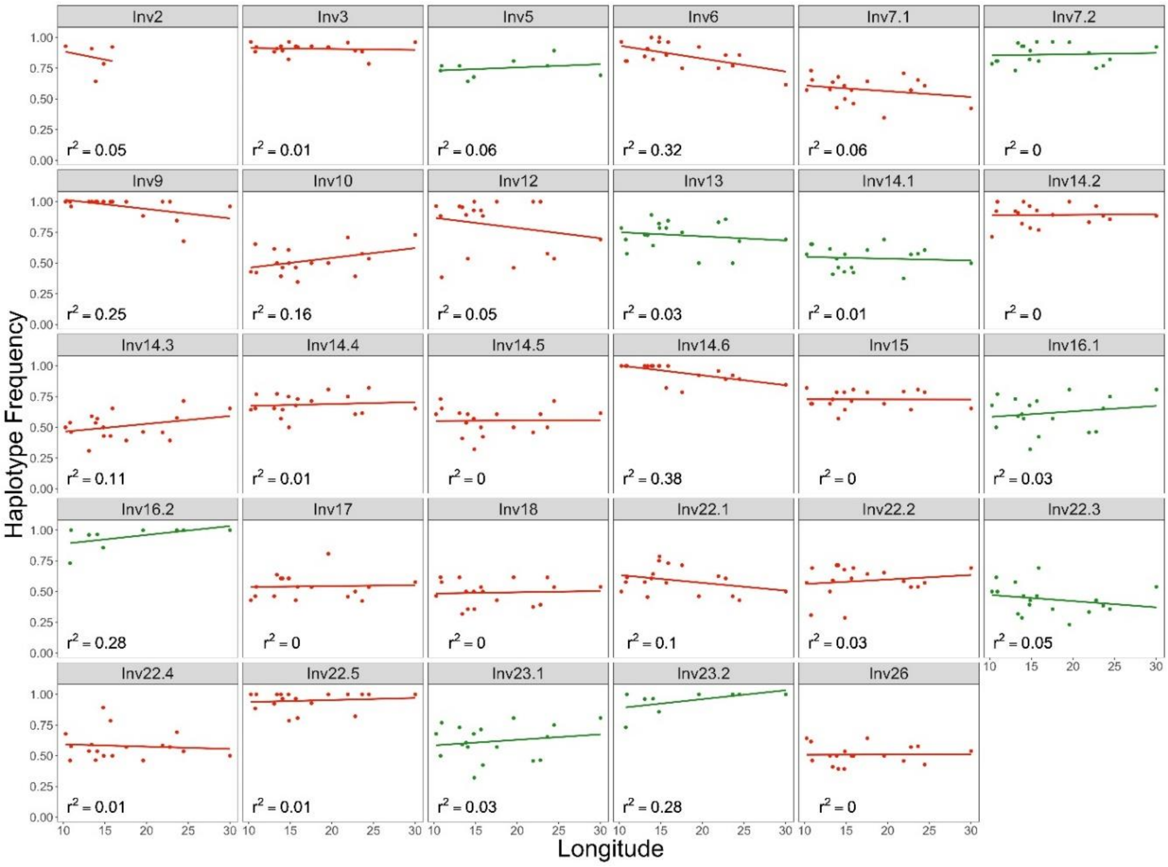

**Figure S12** Examples of inversion haplotypes in the *Ips typographus* genome with (a-b) and without (c-d) haplotype structuring. a and b show two alternative inversion haplotypes of inversion Inv16.1; c and d show two alternative inversion haplotypes of Inv26. e and f show both of inversion haplotypes on one tree, for Inv16.1 and Inv26, respectively. Colored dots correspond to geographic origin of haplotypes (red: southern populations; yellow: Polish populations; blue: northern populations).

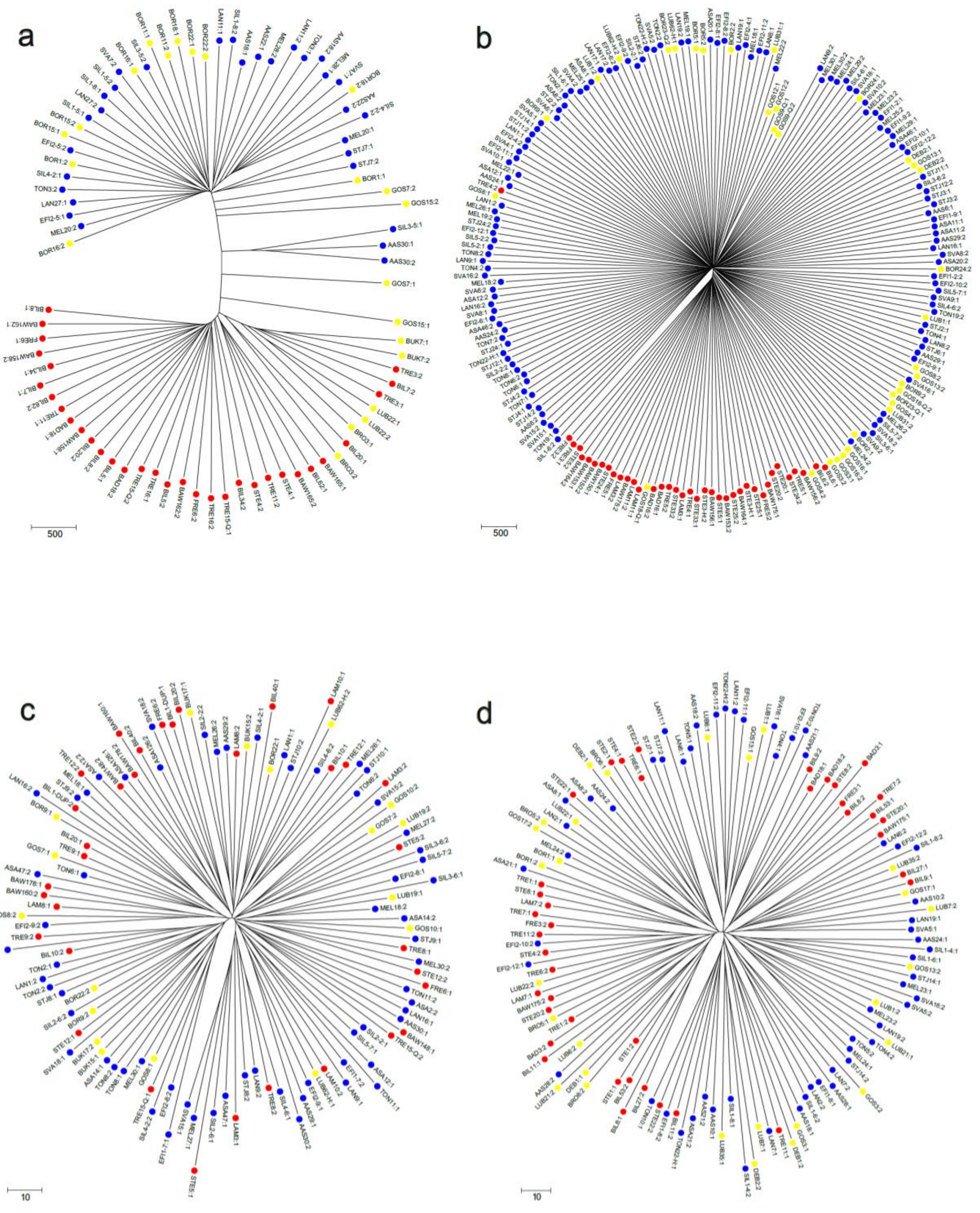

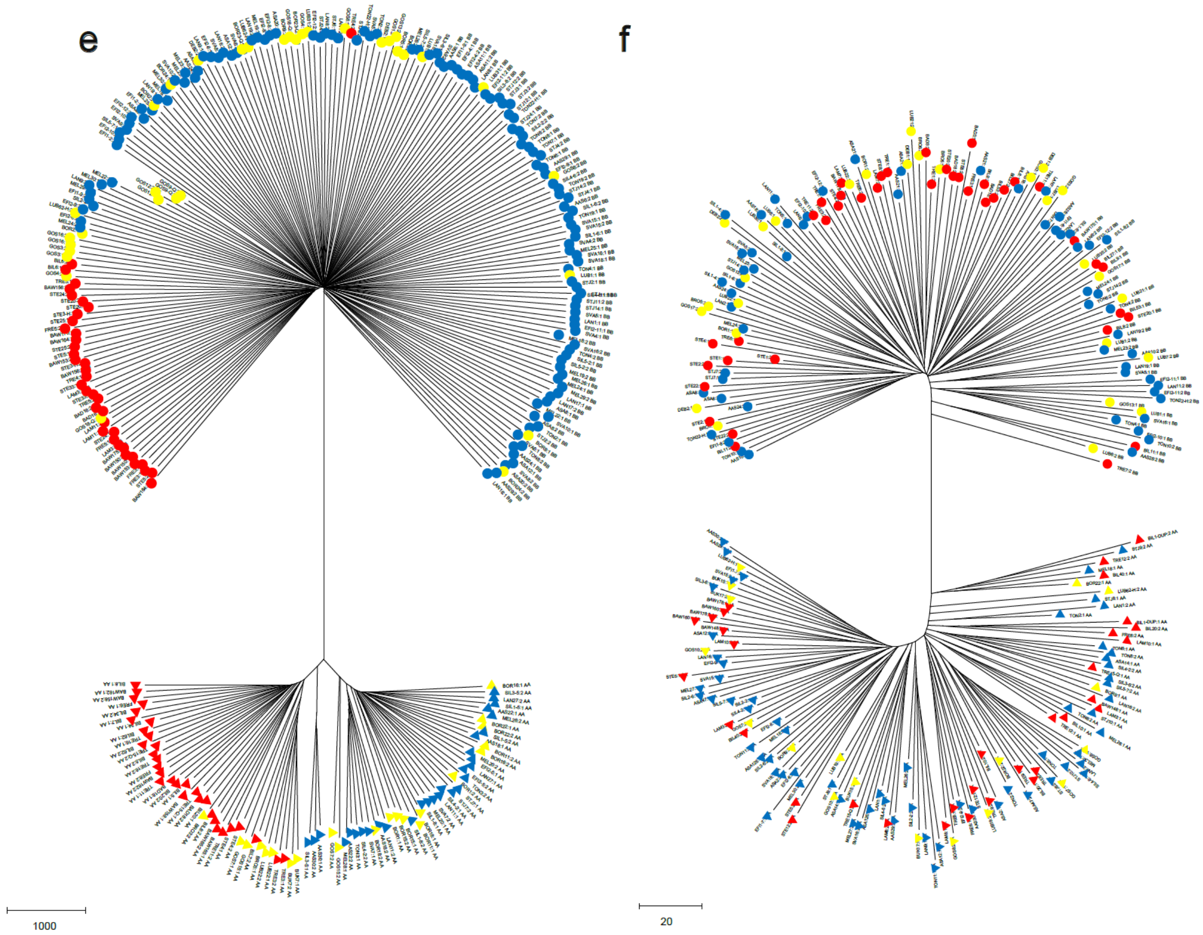
e
f
1000
20

**Figure S13** Examples of NGSadmix results for three inversion regions (a: Inv13; b: Inv15; c: Inv17) in the *Ips typographus* genome showing clear separation into inversion homozygotes (red or blue individuals) or inversion heterozygotes (individuals with red and blue ancestry in approximately equal proportions (the admixture proportions are not exactly 50:50 due to data noise, e.g. different patterns of missing data among individuals or conversion events between two haplotypes). Individuals from different beetle populations are separated by vertical black lines.

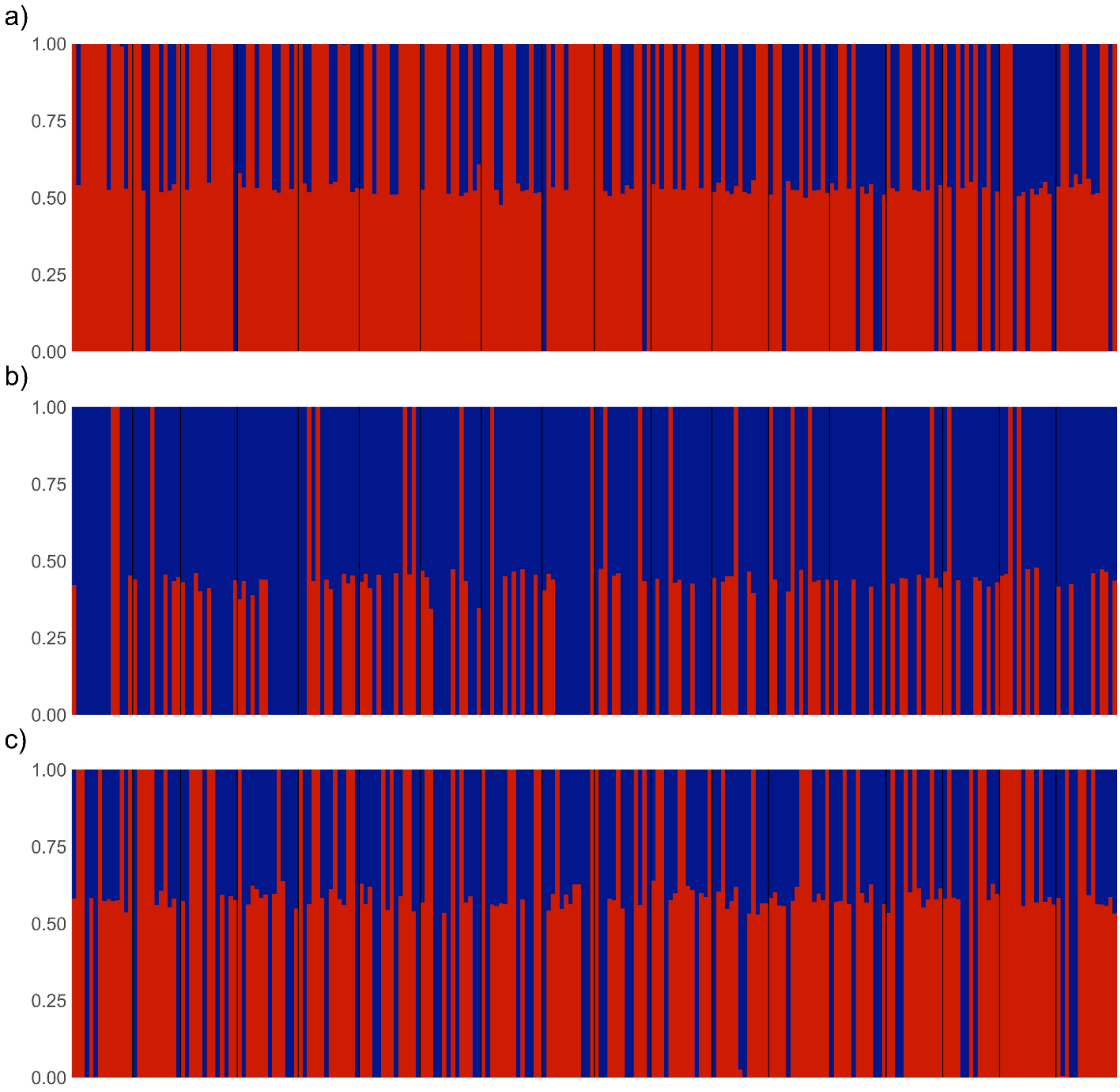

**Figure S14** Genotype-environmental association (GEA) analysis of *Ips typographus* populations collected throughout Europe. The maps show rasters of final environmental variables extracted from a 50 km radius around each population locality. BIO1 (Annual Mean Temperature), BIO2 (Mean Diurnal Range). BIO3 (Isothermality). BIO4 (Temperature Seasonality), BIO5 (Max Temperature of Warmest Month). BIO6 (Min Temperature of Coldest Month), BIO7 (Temperature Annual Range), BIO8 (Mean Temperature of Wettest Quarter). BIO9 (Mean Temperature of Driest Quarter), BIO10 (Mean Temperature of Warmest Quarter), BIO11 (Mean Temperature of Coldest Quarter), BIO12 (Annual Precipitation), BIO13 (Precipitation of Wettest Month), BIO14 (Precipitation of Driest Month), BIO15 (Precipitation Seasonality), BIO16 (Precipitation of Wettest Quarter), BIO17 (Precipitation of Driest Quarter), BIO18 (Precipitation of Warmest Quarter), and BIO19 (Precipitation of Coldest Quarter).

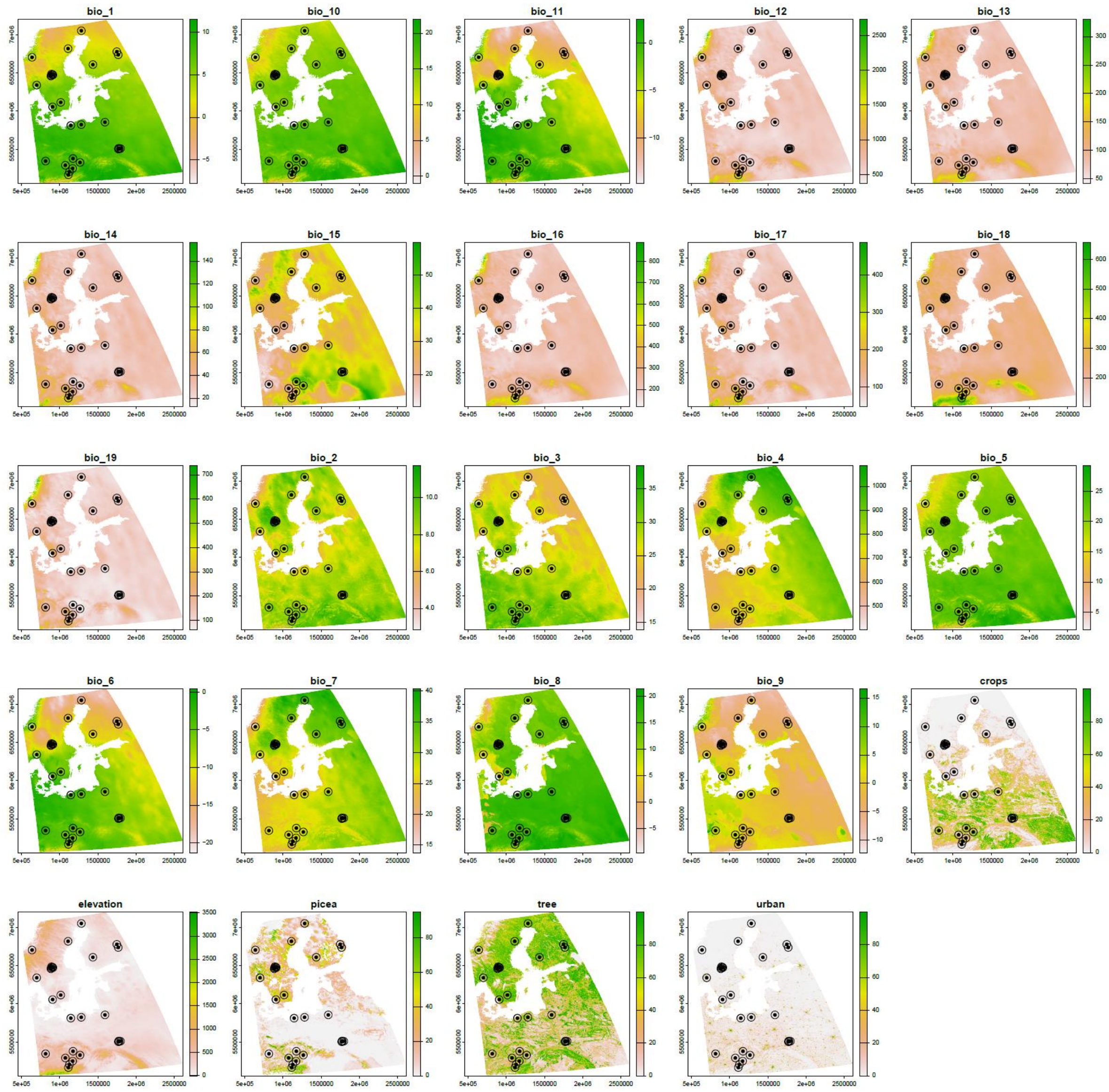
bio_1
bio_10
bio_11
bio_12
bio_13
bio_14
bio_15
bio_16
bio_17
bio_18
bio_19
bio_2
bio_3
bio_4
bio_5
bio_6
bio_7
bio_8
bio_9
crops
elevation
picea
tree
urban

**Figure S15** Genotype-environmental association (GEA) analysis for *Ips typographus* populations collected throughout Europe. Variance explained by all PCs. and loadings of environmental variables in the first two principal components. BIO1 (Annual Mean Temperature), BIO2 (Mean Diurnal Range), BIO3 (Isothermality), BIO4 (Temperature Seasonality), BIO5 (Max Temperature of Warmest Month), BIO6 (Min Temperature of Coldest Month), BIO7 (Temperature Annual Range), BIO8 (Mean Temperature of Wettest Quarter), BIO9 (Mean Temperature of Driest Quarter), BIO10 (Mean Temperature of Warmest Quarter), BIO11 (Mean Temperature of Coldest Quarter), BIO12 (Annual Precipitation), BIO13 (Precipitation of Wettest Month), BIO14 (Precipitation of Driest Month), BIO15 (Precipitation Seasonality), BIO16 (Precipitation of Wettest Quarter), BIO17 (Precipitation of Driest Quarter), BIO18 (Precipitation of Warmest Quarter), and BIO19 (Precipitation of Coldest Quarter).

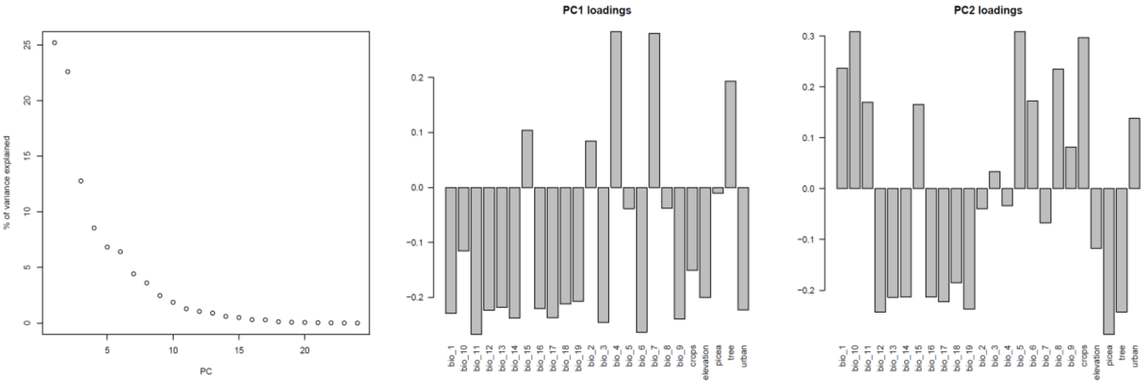
